# Speeding up interval estimation for *R*^2^-based mediation effect of high-dimensional mediators via cross-fitting

**DOI:** 10.1101/2023.02.06.527391

**Authors:** Zhichao Xu, Chunlin Li, Sunyi Chi, Tianzhong Yang, Peng Wei

**Author notes:** Correspondence to T. Yang and P. Wei. These authors contributed equally to this work (Zhichao Xu, Chunlin Li).

## Abstract

Mediation analysis is a useful tool in investigating how molecular phenotypes such as gene expression mediate the effect of exposure on health outcomes. However, commonly used mean-based total mediation effect measures may suffer from cancellation of component-wise mediation effects in opposite directions in the presence of high-dimensional omics mediators. To overcome this limitation, we recently proposed a variance-based R-squared total mediation effect measure that relies on the computationally intensive nonparametric bootstrap for confidence interval estimation. In the work described herein, we formulated a more efficient two-stage, cross-fitted estimation procedure for the *R*^2^ measure. To avoid potential bias, we performed iterative Sure Independence Screening (iSIS) in two subsamples to exclude the non-mediators, followed by ordinary least squares regressions for the variance estimation. We then constructed confidence intervals based on the newly derived closed-form asymptotic distribution of the *R*^2^ measure. Extensive simulation studies demonstrated that this proposed procedure is much more computationally efficient than the resampling-based method, with comparable coverage probability. Furthermore, when applied to the Framingham Heart Study, the proposed method replicated the established finding of gene expression mediating age-related variation in systolic blood pressure and identified the role of gene expression profiles in the relationship between sex and high-density lipoprotein cholesterol level. The proposed estimation procedure is implemented in R package `CFR2M`.

## 1 Introduction

Recent advances in high-throughput technologies have enabled researchers to measure thousands or even millions of molecular variables such as DNA methylation and gene expression in a variety of tissues and cells, providing unprecedented opportunities to study biological mechanisms. High-dimensional mediation analysis is a critical research area in which the role of molecular phenotypes such as gene expression in mediating the effect of exposure on health outcomes is explored. Most existing high-dimensional mediation analysis methods rely on mean-based total mediation effect size measures (Zhao and Luo, 2022; Huang and Pan, 2016; Dai et al., 2022; Song et al., 2020; Zeng et al., 2021). However, as shown in real data applications, component-wise mediation effects in the realm of high-dimensional genomic mediators often exhibit opposite directions. These mean-based measures may not adequately capture the entirety of the total mediation effect, as it can be obscured by the cancellation of component-wise mediation effects in opposite directions. As a complement, Yang et al. (2021) proposed a variance-based R-squared measure for the total mediation effect, denoted as 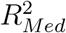, in the high-dimensional setting. It provides useful insights particularly when individual molecular mediators display mediation effects in opposite directions. In this work, we focus on the total mediation effect rather than the component-wise or path-specific mediation effect, whose identification and estimation are different topics that necessitates a more comprehensive treatment (Huber, 2019; Avin et al., 2005).

Researchers originally proposed the R-squared measure in the framework of commonality analysis under the single-mediator model (Fairchild et al., 2009). In the multiple or high-dimensional mediation analysis framework, 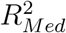 is defined as the variance of the outcome variable that is common to both exposure and mediators, or taking it a step further, explained by the exposure through the mediators (Fairchild et al., 2009; Yang et al., 2021). Such variance-based measures are well accepted in genetic and genomic research. For example, the genetic heritability measure that quantifies the proportion of phenotypic variance attributable to genetic variance is a long-standing and still active focus of research and development (Visscher and Goddard, 2019). Mirroring this,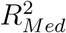 partitions the variance owing to mediation effects, providing a clear and interpretable measure for the community.

The R-squared measure is essentially an additive function of the variance of the outcome explained by the exposure, mediators, and exposure and mediators. Estimating variance under the high-dimensional setting is generally challenging and has been less explored than parameter estimation and hypothesis testing of component-wise mediation effects (Zhao and Luo, 2022; Gao et al., 2019; Dai et al., 2022; Zeng et al., 2021; Fang et al., 2020; Derkach et al., 2020; Liu et al., 2022). As demonstrated in Yang et al. (2021), 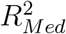 can be seriously biased when spurious mediators are included. Specifically, the estimate of 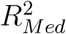 becomes inconsistent in the presence of spurious mediators that have no effect on the dependent variable. In real data analysis with high-dimensional mediators, the identity of the true mediators is rarely known *a priori*, and they are hard to distinguish from the spurious ones with a finite sample. The earlier work by Yang et al. (2021) used a variable selection method with the oracle property (Fan and Li, 2001) to filter out spurious variables based on half of the sample and estimated 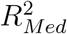 using mixed-effect models based on the remaining half. This data-splitting strategy decreases the estimation efficiency owing to insufficient usage of the whole sample. Yang et al. (2021) used a nonparametric bootstrap to compute confidence intervals, which demonstrated satisfactory coverage probability, but was computationally intensive, as each iteration of the bootstrap involved a variable selection step and an estimation step. Furthermore, Yang et al. (2021) focused on a situation in which mediators are conditionally independent given the exposure, an oversimplification in real data analysis.

We herein propose a new two-stage cross-fitted interval estimation procedure for 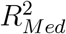 1) enhances estimation efficiency by leveraging a whole sample via cross-fitting, 2) is much faster than the nonparametric bootstrap, and 3) can improve mediator selection against spurious correlations. We derive the asymptotic distribution of the 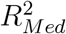 estimator and demonstrate that the resulting asymptotic confidence intervals have satisfactory coverage probabilities comparable with those of the bootstrap-based confidence intervals in extensive simulation settings. Using this newly proposed estimation procedure, we replicated a previously established mediating relationship among age, gene expression, and systolic blood pressure (BP) (Yang et al., 2021) and investigated how gene expression mediates the well-known relationship between sex and high-density lipoprotein cholesterol (HDL-C) level (Lawlor et al., 2001; Weidner et al., 1991; Wilson et al., 1983) in the Framingham Heart Study (FHS). Lastly, we implemented our new estimation procedure in `CFR2M` R package on the `CRAN`.

## 2 Methods

### 2.1 Mediation model and *R*^2^ measure

Mediation analysis is commonly used to investigate the role of intermediate variable(s) in the relationship between two variables (an exposure variable and an outcome variable), enabling researchers to understand the mechanisms underlying the effect of the exposure variable on the outcome variable. A mediation model consists of the following equations ((VanderWeele and Vansteelandt, 2014)),

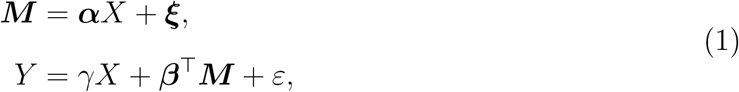

where *X* is an exposure variable; *Y* is a response variable; ***M*** is a vector of *p* potential mediators; ***ξ***, *ε* are errors; and ***α, β***, *γ* are regression coefficients. Without loss of generality, we assume that all variables are centered at 0 and scaled to have variance of 1. In addition, all measured potential confounders have been regressed out from *X, M*_*j*_’s and *Y* in the above equations. Using the counterfactual framework, the indirect or mediation effects can be identified under the following assumptions (VanderWeele and Vansteelandt, 2014; Imai and Yamamoto, 2013; Albert and Nelson, 2011): (1) no unmeasured confounders between the exposure and outcome, (2) no unmeasured confounding in the relationship between the mediator and outcome, (3) no unmeasured confounding between the exposure and mediator, and (4) no exposure-induced confounding in the relationship between the mediator and outcome. The natural indirect effect (NIE), an counterfactual entity, is a first-moment measure, i.e., NIE = ***α***^⊤^***β*** under assumptions (1)–(4) and Equation 1. Without being directly linked to the counterfactual framework, the product, proportion, and ratio mediation effect measures have been widely used in the literature (MacKinnon, 2008). Given the equations 1, the product measure is defined as ***α***^⊤^***β***, which coincides with the NIE under certain conditions.

The proportion measure is characterized as the fraction of the total effect mediated by the mediators, denoted as 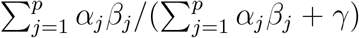, where *γ* is the direct effect. The total effect measure is defined as 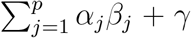. Generally speaking, the aforementioned four identifiability assumptions are required to obtain unbiased estimates for *α, β* and *γ* in causal mediation analysis.

Yang et al. (2021) adapted the concepts in commonality analysis and proposed an extension of the R-squared measure in the single-mediator model to high-dimensional mediation analysis. The measure is defined as

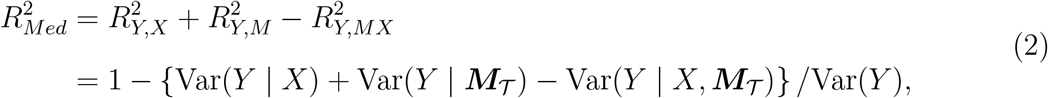

where ***M***_𝒯_ is denoted as the set of true mediators, 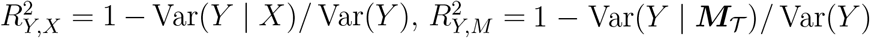, and 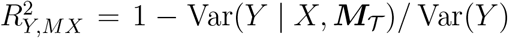 are the coefficients of determination of *Y* regressing over ***M***_𝒯_, *X*, and (*X*, ***M***_𝒯_), respectively (Fairchild et al., 2009). In equation (2), 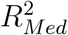 is constituted by the variance *V*_*Y*_ = Var(*Y*) and conditional variances *V*_*Y* |*X*_ = Var(*Y* | *X*), *V*_*Y* |*M*_ = Var(*Y* | ***M***_𝒯_), and *V*_*Y* |*MX*_ = Var(*Y* | *X*, ***M***_𝒯_). This observation suggests that the 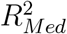 estimation can be reduced to variance estimation in regressions. When no mediator is present (i.e., 𝒯 = ∅), we have 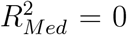. Estimation of each *R*^2^ measure in equation (2) requires control of confounding effects.

In high-dimensional mediation analysis, the identity of the true mediator is usually unknown. The potential mediators 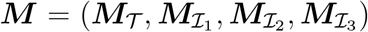 are partitioned into true mediators and three types of non-mediators, respectively. As illustrated in Figure 1, the true mediators ***M***_𝒯_ has *α*_*j*_ ≠ 0 and *β*_*j*_ ≠ 0 for *j* ∈ 𝒯), the non-mediators 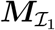 are only affecting the outcome (*α*_*j*_ = 0 and *β*_*j*_ ≠ 0 for *j* ∈ *ℐ*_1_), the non-mediators 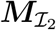 are only affected by the exposure (*α*_*j*_ ≠ 0 and *β*_*j*_ = 0 for *j* ∈ ℐ_2_), and the noise variables 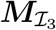 are neither affected by the exposure nor affecting the outcome (*α*_*j*_ = 0 and *β*_*j*_ = 0 for *j* ∈ ℐ_3_). Non-mediators can potentially distort the mediation effect. For example, when ***ξ*** is correlated, 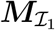 and 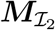 becomes the mediator-outcome and exposure-outcome confounders, respectively, violating assumptions (1) and (2). On the other hand, inclusion of 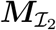 in the model has been demonstrated to bias the estimation of 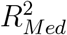 because of the model misspecification when calculating 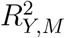 (Yang et al., 2021).

**Figure 1:**
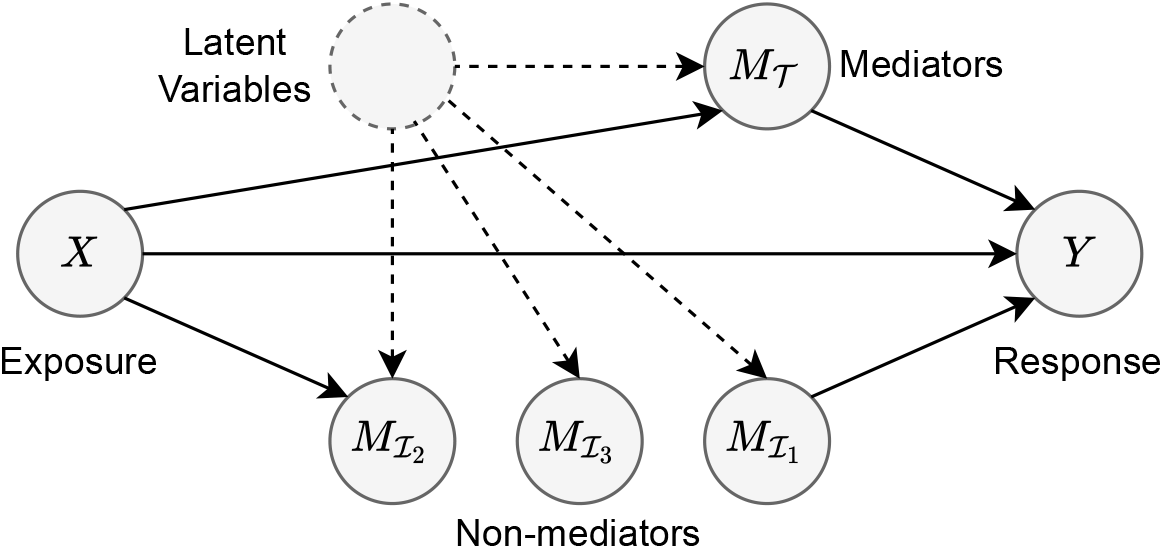
Graphical representation of a mediation model where the latent variables introduce correlations among putative mediators.

### 2.2 Cross-fitted estimation of the *R*^2^ measure

Motivated by Fan et al. (2012), we propose an estimation procedure for 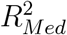 based on sample splitting and cross-fitting. To proceed, suppose that an independent and identically distributed sample 𝒟 = {(*X*_*i*_, *Y*_*i*_, ***M***_*i*_) : *i* = 1, …, *n*} is given. The procedure is summarized in Figure 2 and detailed as follows:

**Figure 2:**
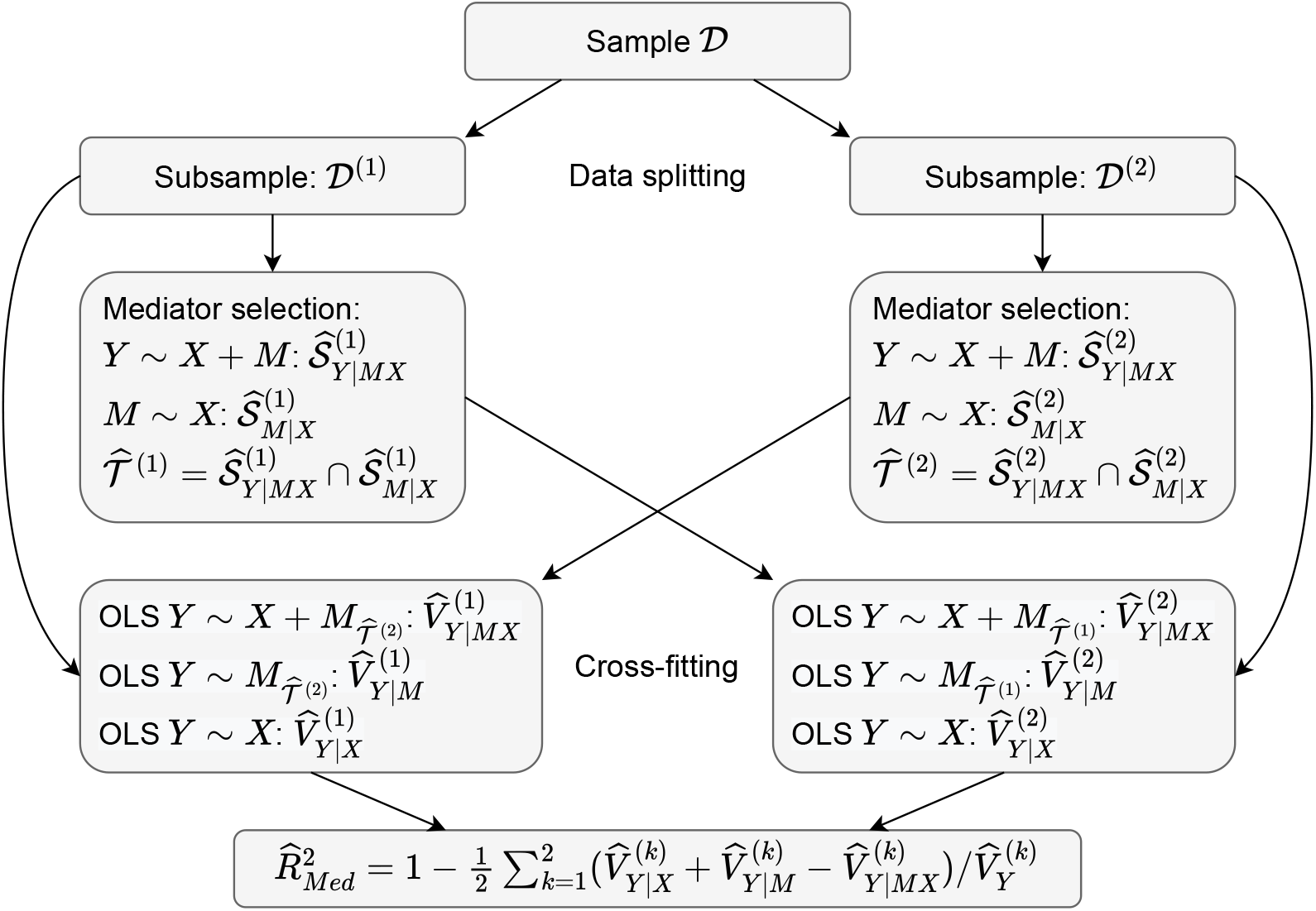
Cross-fitted estimation of 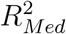. The sample 𝒟 is split into 𝒟 ^(1)^ and 𝒟^(2)^. 𝒟^(*k*)^ is then used for mediator selection 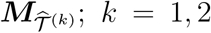. Next, 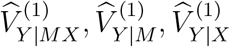, and 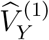 are estimated based on 𝒟 ^(1)^ and the selected mediators 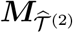, and similarly for 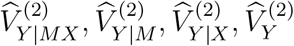. Finally, 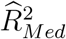 is computed.

- **(Data splitting)** The original sample 𝒟 is randomly split into two equal subsamples 𝒟^(1)^ and 𝒟^(2)^.
- **(Cross-fitting)** A mediator selection method is applied to 𝒟 ^(1)^, and *V*_*Y*_, *V*_*Y* |*X*_, *V*_*Y* |*M*_, and *V*_*Y* |*MX*_ are estimated based on 𝒟 ^(2)^. For example, iterative Sure Independence Screening (iSIS) (Fan and Lv, 2008) is used along with the Minimax Concave Penalty (MCP) (Zhang, 2010) screening procedure to select the mediator index set in each subsample. The roles of 𝒟^(1)^, 𝒟^(2)^ are then exchanged, and the procedure is repeated. Specifically, 𝒟^(1)^ is used to compute the regression of *Y* over (*X*, ***M***) and regressions of ***M*** over *X*. Let 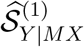 be the selected mediator index set by regressing *Y* over (*X,M*), let 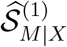 be the selected mediator index set by regressing ***M*** over *X*, and let 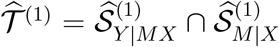 be the estimated mediator index set based on 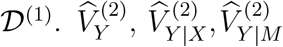, and 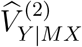 are then computed using 𝒟^(2)^, where 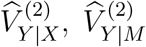, and 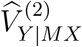 are computed by fitting ordinary least squares (OLS) regressions of *Y* over *X*, 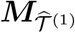, and 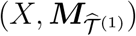, respectively. Next 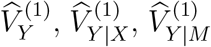, and 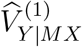 are computed in a similar way, with 𝒟 ^(1)^ and 𝒟 ^(2)^ being switched.
- The final estimate is 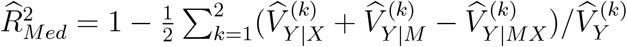.

The proposed method comprises two essential ingredients: data splitting and cross-fitting. Splitting a sample reduces the bias incurred by the mediator selection. As to be seen in Theorem 1, data splitting allows for lifting of the oracle property (i.e., asymptotically exact variable selection) (Fan and Li, 2001) for mediator selection. This significantly improves the results reported by Yang et al. (2021) because exact selection is rarely achieved in high-dimensional situations. Despite this attractive property, data splitting may result in loss of estimation efficiency when using a subset of data. The cross-fitting procedure, on the other hand, enables usage of all the data, yielding a more efficient estimator than that described by Yang et al. (2021). Importantly, according to Theorem 1, the proposed estimator achieves the same asymptotic efficiency as the hypothesized oracle estimator based on a full sample. In other words, the efficiency loss owing to data splitting becomes negligible after cross-fitting.

### 2.3 Theoretical properties and interval estimation

In this subsection, the large-sample properties of the proposed cross-fitted estimator are established. In particular, the asymptotic normality of conditional variance estimators is derived, which enables us to construct confidence intervals for the R-squared measure 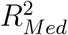.

For clarity of presentation, *X*, ***ξ***, and *ε* in equation (1) are assumed to be independently and normally distributed, where the components of ***ξ*** ∼ *N* (**0, Σ**) are allowed to be correlated. Of note is that normality is not essential and our theoretical result can be readily extended to a sub-Gaussian case (standard high-dimensional setting). However, relaxation of normality requires additional complications (see the discussion in Supplementary Materials Web Appendix A).

The cross-fitting procedure involves mediator selection that will affect the 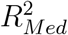 estimation quality. For our analysis, the assumptions (1-3) are described below.

#### Condition 1.

(Sure screening property) The mediator selection satisfies the property 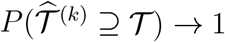 as *n* → ∞ for *k* = 1, 2.

In Assumption 1, the sure screening property (Fan and Lv, 2008) is required. Notably, the selection method does not have to possess the selection consistency or oracle property. This constitutes a significant relaxation compared with the restrictions described by Yang et al. (2021), and it aligns with our empirical results described in section 3.

#### Condition 2.

Suppose 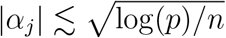 and 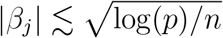 for *j* ∉ 𝒯.

Of note is that when nonzero signals 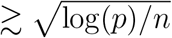, the oracle property or the sure screening property is achievable. Thus, our estimation procedure can exclude such non-mediators. On the other hand, for non-mediators with weak effects (i.e., signals 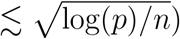, as given by Assumption 2, the exact selection may not be possible according to the information-theoretic limit. Therefore, such non-mediators are incorporated into the derivation of Theorem 2.1.

#### Condition 3.

Suppose max 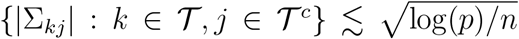 and *c*_1_ ≤ *λ*_min_(**Σ**) ≤ *λ*_max_(**Σ**) ≤ *c*_2_, where **Σ** is the covariance of ***ξ***.

Assumption 3 is a regularity condition on **Σ**, requiring that ***ξ*** is not too correlated. Notably, a correlated ***ξ*** suggests a violation of the parallel mediators assumption, which could result from uncontrolled confounding effects (Yuan and Qu, 2023). Thus, deriving the asymptotic properties of 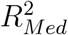 under Assumption 3 is reasonable. It is also important to note that these conditions are sufficient but not necessary. Overall, our analysis largely adheres to the assumptions. A detailed discussion of these assumptions in both the real data and simulations is provided in Supplementary Materials Web Appendix D. Remarkably, in our real data application described below, including principal components of high-dimensional genomic mediators as covariates can effectively reduce the correlations among mediators owing to residual confounding. Furthermore, 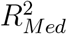 has shown to be robust to violation of this assumption under low-dimensional settings Yang et al. (2021) and under high-dimensional settings in our simulations (Section 3.1).

#### Theorem 1.

*Suppose Assumption 1, Assumption 2 and Assumption 3 are met. If* |𝒯| + |ℐ_1_| + |ℐ_2_| ≤ *s*, 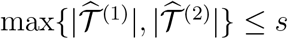 *and* 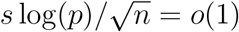, *then we have*

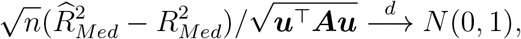

*where* 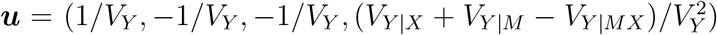 *and* ***A*** *is the (constant) covariance matrix of* (*ε*^2^, *η*^2^, ζ^2^, *Y* ^2^).

For statistical inference, the asymptotic covariance matrix ***A*** is estimated by the residuals of the corresponding least squares regressions, and the plugin estimator 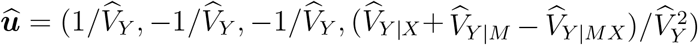 is used for ***u***. Detailed technical proof of Theorem 1 is provided in Supplementary Materials Web Appendix A.

As suggested by Theorem 1, the estimator 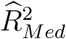 is consistent and achieves the asymptotic variance of the hypothetical oracle estimator. Thus, there is asymptotically no loss of efficiency for statistical inference.

We considered the Shared Over Simple (SOS) measure. Defined as 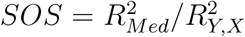, this measure represents the standardized variance in the outcome related to the exposure that intersects the mediator (Lindenberger and Potter, 1998). Derivation of the asymptotic distribution of SOS can be found in the Supplementary Materials Web Appendix A.

## 3 Simulation studies

### 3.1 Simulation settings

We first compared the proposed cross-fitted OLS estimation method (CF-OLS) with a previously established method (shortened as B-Mixed) (Yang et al., 2021), which estimates the 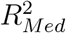 measure in a mixed model framework along with a bootstrap-based confidence interval. As shown by Yang et al. (2021), the existence of the non-mediator 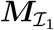 and noise variables did not affect the estimation, whereas the non-mediator 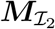 can result in a biased, inconsistent estimation when mediators are conditionally independent in high-dimensional settings. Therefore, we used the iterative Sure Independence Screening (iSIS) along with the Minimax Concave Penalty (MCP) screening procedure (iSIS-MCP) for variable selection to exclude the non-mediators 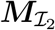. Subsequently, we assessed the performance of the CF-OLS method, increasing correlations among potential mediators to better mimic the characteristics of omics data. In this case, 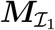 became mediator-outcome confounders, and 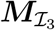 became exposure-outcome confounders. In these scenarios, we applied the false discovery rate (FDR) control along with iSIS-MCP to filter out the non-mediators 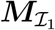 and 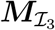. We computed the coverage probability, width of the confidence interval, bias, mean squared error (MSE), empirical standard deviation of the estimator (i.e., standard deviation of the sampling distribution of the estimator based on simulation replications), variable selection accuracy, and computational efficiency in various high-dimensional settings.

More specifically, for the B-Mixed method, we applied variable selection to the first half subsample and obtained point estimation and confidence intervals in the second half subsample. For each replication, the confidence interval for 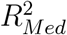 was computed from 500 nonparametric bootstrap resamplings. We then obtained the coverage probability and empirical standard deviation for the estimation from 200 replications. For the CF-OLS method, within each replication, we applied variable selection independently to two subsamples as illustrated in Figure 2. The asymptotic standard error, bias, MSE, true positive rate, and false positive rate were the mean values of their respective estimates in the subsamples. Next, we constructed the Wald confidence interval for 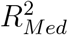 was constructed based on the asymptotic standard error. We directly reported the coverage probability and empirical standard deviation of the estimation from 200 replications. For both methods, we averaged the confidence interval width, bias, MSE, true positive rate, and false positive rate across 200 replications. We evaluated the performance of the two methods in various scenarios (A1)–(A12) that included different types or numbers of non-mediators were included. Specifically, in scenarios (A1)–(A6), we evaluated both methods under the assumption of independence, whereas in scenarios (A7)–(A12), we focused on the CF-OLS method with correlated putative mediators. In scenarios (A1), (A2), (A8), and (A9), we added a substantial number of noise variables 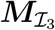 to the true mediators ***M***_𝒯_. In scenarios (A3) and (A10), we included a large quantity of non-mediators 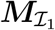. In scenarios (A4) and (A11), we simulated non-mediators 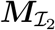.

In scenarios (A5), (A6), and (A12), we examined a combination of different types of non-mediators. Finally, in scenario (A7), we considered all variables to be non-mediators.

In each scenario, we simulated the same parameters across 200 replications so that the true 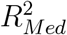 remained the same. We simulated data sets using Equation 1 with sample sizes of 750, 1500, and 3000. Also, we simulated exposure variable *X* from the standard normal distribution *N* (0, 1) and set coefficient *γ* in Equation 1 to 3. Let (*p*_0_, *p*_1_, *p*_2_, *p*_3_) denote the number of true mediators, two types of non-mediators, and noise variables 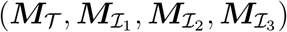, respectively. We set the total number of variables in ***M*** to 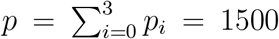. The errors in Equation 1 for scenarios (A1)–(A6) independently follow the standard normal distribution, ***ξ*** ∼ *N* (**0, *I***_*p*_) and *ε* ∼ *N* (0, 1). In scenarios (A7)–(A12), we considered two different correlation structures for the putative mediators. For the first correlation structure, ***ξ*** ∼ *N* (**0**, diag(**Σ, *I***_*p*2+*p*3_)) where **Σ**_*ij*_ = 0.2 for 1 ≤ *i* ≠ *j* ≤ *p*_0_ + *p*_1_ and **Σ**_*ij*_ = 1 for 1 ≤ *i* = *j* ≤ *p*_0_ + *p*_1_. For the second correlation structure, we considered ***ξ*** ∼ *N* (**0**, diag(**Σ, *I***_*p*2 +*p*3_)) where **Σ**_*ij*_’s are iid samples from *N* (0, 0.1^2^) for 1 ≤ *i* ≠*j* ≤ *p*_0_ + *p*_1_ and **Σ**_*ij*_ = 1 for 1 ≤ *i* = *j* ≤ *p*_0_ +*p*_1_. We set the maximum number of iterations for iSIS equal to 3. We also calculated the bias and MSE of the mean-based mediation effect measures (product, proportion, and total effect measures) and the SOS measure in these simulation scenarios.

The details of simulation scenarios (A1)–(A12) were shown as follows:

- **(A1)** (*p*_0_, *p*_1_, *p*_2_, *p*_3_) = (15, 0, 0, 1485): *α*_*i*_ ∼ *N* (0, 1.5^2^), *β*_*i*_ ∼ *N* (0, 1.5^2^) for *i* = 1, …, 15; *α*_*i*_ = *β*_*i*_ = 0 for *i* = 16, …, 1500.
- **(A2)** (*p*_0_, *p*_1_, *p*_2_, *p*_3_) = (150, 0, 0, 1350): *α*_*i*_ ∼ *N* (0, 1.5^2^), *β*_*i*_ ∼ *N* (0, 1.5^2^) for *i* = 1, …, 150; *α*_*i*_ = *β*_*i*_ = 0 for *i* = 151, …, 1500.
- **(A3)** (*p*_0_, *p*_1_, *p*_2_, *p*_3_) = (150, 1350, 0, 0): *α*_*i*_ ∼ *N* (0, 1.5^2^), *β*_*i*_ ∼ *N* (0, 1.5^2^) for *i* = 1, …, 150; *α*_*i*_ = 0, *β*_*i*_ ∼ *N* (0, 1.5^2^) for *i* = 151, …, 1500.
- **(A4)** (*p*_0_, *p*_1_, *p*_2_, *p*_3_) = (150, 0, 1350, 0): *α*_*i*_ ∼ *N* (0, 1.5^2^), *β*_*i*_ ∼ *N* (0, 1.5^2^) for *i* = 1, …, 150; *α*_*i*_ ∼ *N* (0, 1.5^2^), *β*_*i*_ = 0 for *i* = 151, …, 1500.
- **(A5)** (*p*_0_, *p*_1_, *p*_2_, *p*_3_) = (150, 150, 0, 1200): *α*_*i*_ ∼ *N* (0, 1.5^2^), *β*_*i*_ ∼ *N* (0, 1.5^2^) for *i* = 1, …, 150; *α*_*i*_ = 0, *β*_*i*_ ∼ *N* (0, 1.5^2^) for *i* = 151, …, 300; *α*_*i*_ = *β*_*i*_ = 0 for *i* = 301, …, 1500.
- **(A6)** (*p*_0_, *p*_1_, *p*_2_, *p*_3_) = (150, 150, 150, 1050): *α*_*i*_ ∼ *N* (0, 1.5^2^), *β*_*i*_ ∼ *N* (0, 1.5^2^) for *i* = 1, …, 150; *α*_*i*_ = 0, *β*_*i*_ ∼ *N* (0, 1.5^2^) for *i* = 151, …, 300; *α*_*i*_ ∼ *N* (0, 1.5^2^), *β*_*i*_ = 0 for *i* = 301, …, 450; *α*_*i*_ = *β*_*i*_ = 0 for *i* = 451, …, 1500.
- **(A7)** (*p*_0_, *p*_1_, *p*_2_, *p*_3_) = (0, 20, 20, 1460): *α*_*i*_ = 0, *β*_*i*_ ∼ *N* (0, 1.5^2^) for *i* = 1, …, 20; *α*_*i*_ ∼ *N* (0, 1.5^2^), *β*_*i*_ = 0 for *i* = 21, …, 40; *α*_*i*_ = *β*_*i*_ = 0 for *i* = 41, …, 1500.
- **(A8)** (*p*_0_, *p*_1_, *p*_2_, *p*_3_) = (5, 0, 0, 1495): *α*_*i*_ ∼ *N* (0, 1.5^2^), *β*_*i*_ ∼ *N* (0, 1.5^2^) for *i* = 1, …, 5; *α*_*i*_ = *β*_*i*_ = 0 for *i* = 6, …, 1500.
- **(A9)** (*p*_0_, *p*_1_, *p*_2_, *p*_3_) = (20, 0, 0, 1480): *α*_*i*_ ∼ *N* (0, 1.5^2^), *β*_*i*_ ∼ *N* (0, 1.5^2^) for *i* = 1, …, 20; *α*_*i*_ = *β*_*i*_ = 0 for *i* = 21, …, 1500.
- **(A10)** (*p*_0_, *p*_1_, *p*_2_, *p*_3_) = (20, 60, 0, 1420): *α*_*i*_ ∼ *N* (0, 1.5^2^), *β*_*i*_ ∼ *N* (0, 1.5^2^) for *i* = 1, …, 20; *α*_*i*_ = 0, *β*_*i*_ ∼ *N* (0, 1.5^2^) for *i* = 21, …, 80; *α*_*i*_ = *β*_*i*_ = 0 for *i* = 81, …, 1500.
- **(A11)** (*p*_0_, *p*_1_, *p*_2_, *p*_3_) = (20, 0, 60, 1420): *α*_*i*_ ∼ *N* (0, 1.5^2^), *β*_*i*_ ∼ *N* (0, 1.5^2^) for *i* = 1, …, 20; *α*_*i*_ ∼ *N* (0, 1.5^2^), *β*_*i*_ = 0 for *i* = 21, …, 80; *α*_*i*_ = *β*_*i*_ = 0 for *i* = 81, …, 1500.
- **(A12)** (*p*_0_, *p*_1_, *p*_2_, *p*_3_) = (20, 60, 60, 1360): *α*_*i*_ ∼ *N* (0, 1.5^2^), *β*_*i*_ ∼ *N* (0, 1.5^2^) for *i* = 1, …, 20; *α*_*i*_ = 0, *β*_*i*_ ∼ *N* (0, 1.5^2^) for *i* = 21, …, 80; *α*_*i*_ ∼ *N* (0, 1.5^2^), *β*_*i*_ = 0 for *i* = 81, …, 140; *α*_*i*_ = *β*_*i*_ = 0 for *i* = 141, …, 1500.

### 3.2 Simulation results

Table 1 compares the statistical inference for independent putative mediators under the high-dimensional setting with the CF-OLS and the B-Mixed methods. In general, CF-OLS performed reasonably well in all scenarios. In this section, we present the results based on the iSIS-MCP variable selection alone, whereas the results based on both iSIS-MCP and the FDR control to additionally filter out 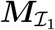 non-mediators, which were very similar to those without the FDR as shown in the Supplementary Materials Web Appendix B.

**Table 1:** Simulation results using the CF-OLS and B-Mixed methods with independent mediators in scenarios (A1)–(A6). **N** refers to the sample size. **CP** refers to coverage probability based on 200 replications. **Width** refers to half the width of the 95% confidence interval. **SE** refers to the average asymptotic standard error. **SD** refers to the empirical standard deviation of replicated estimations. MSE refers to mean squared error. **TP** refers to the average true positive rate. **FP** refers to the average false positive rate. The true value of 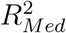 is shown in parentheses. **Time** refers to the mean computational time in minutes for each replication with its standard error shown in parentheses. The computational time for CF-OLS was observed using a single CPU core. The computational time for B-Mixed was observed using 20 cores in parallel.

| Scenario | N | CF-OLS |  |  |  |  |  |  |  |  | B-Mixed |  |  |  |  |  |  |  |  |
| --- | --- | --- | --- | --- | --- | --- | --- | --- | --- | --- | --- | --- | --- | --- | --- | --- | --- | --- | --- |
|  |  | CP | Width | SE | Bias | SD | MSE | TP | FP | Time | CP | Width | Bias | SD | MSE | TP | FP | Time |  |
| $(R^2_{Med})$ | | % | ($\times 10^{-2}$) | ($\times 10^{-2}$) | ($\times 10^{-2}$) | ($\times 10^{-2}$) | ($\times 10^{-4}$) | % | % | (mins) | % | ($\times 10^{-2}$) | ($\times 10^{-2}$) | ($\times 10^{-2}$) | ($\times 10^{-4}$) | % | % | (mins) | |
| A1<br>(0.065) | 750 | 92.0 | 3.664 | 1.870 | 0.739 | 1.940 | 4.292 | 94.5 | 2.1 | 0.12 (0.00) | 98.5 | 5.159 | 0.149 | 2.646 | 6.990 | 94.0 | 2.0 | 44.96 (2.27) |  |
|  | 1500 | 93.5 | 2.601 | 1.327 | 0.658 | 1.316 | 2.155 | 92.9 | 1.8 | 3.44 (0.04) | 95.0 | 3.615 | 0.236 | 2.084 | 4.377 | 92.3 | 1.5 | 85.09 (4.44) |  |
|  | 3000 | 93.5 | 1.844 | 0.941 | 0.133 | 0.994 | 1.001 | 96.7 | 0.8 | 4.80 (0.07) | 93.0 | 2.591 | 0.138 | 1.491 | 2.230 | 96.8 | 0.8 | 153.49 (8.12) |  |
| A2<br>(0.418) | 750 | 94.5 | 5.383 | 2.747 | -0.032 | 2.736 | 7.450 | 40.3 | 0.1 | 1.98 (0.04) | 95.0 | 7.702 | -0.263 | 3.908 | 15.266 | 40.2 | 0.1 | 51.23 (2.83) |  |
|  | 1500 | 92.0 | 3.787 | 1.932 | 0.334 | 1.956 | 3.920 | 69.4 | 0.3 | 5.30 (0.11) | 94.0 | 5.353 | 0.355 | 2.647 | 7.097 | 69.6 | 0.3 | 88.22 (4.54) |  |
|  | 3000 | 94.5 | 2.691 | 1.373 | -0.131 | 1.390 | 1.940 | 94.3 | 0.3 | 6.78 (0.04) | 94.0 | 3.777 | -0.103 | 1.953 | 3.807 | 94.3 | 0.2 | 149.68 (6.28) |  |
| A3<br>(0.064) | 750 | 93.5 | 3.494 | 1.782 | 0.269 | 1.790 | 3.259 | 31.0 | 1.1 | 2.13 (0.04) | 92.5 | 5.054 | 0.365 | 2.762 | 7.725 | 31.1 | 1.1 | 38.51 (1.56) |  |
|  | 1500 | 95.0 | 2.431 | 1.240 | 0.198 | 1.259 | 1.617 | 50.5 | 2.6 | 5.10 (0.05) | 94.0 | 3.390 | -0.008 | 1.820 | 3.297 | 50.6 | 2.6 | 74.06 (2.69) |  |
|  | 3000 | 95.0 | 1.707 | 0.871 | 0.168 | 0.817 | 0.692 | 76.2 | 6.5 | 8.62 (0.10) | 96.0 | 2.391 | 0.015 | 1.118 | 1.245 | 76.3 | 6.5 | 147.08 (4.46) |  |
| A4<br>(0.390) | 750 | 96.0 | 5.445 | 2.778 | 0.029 | 2.769 | 7.630 | 13.0 | 2.5 | 1.47 (0.03) | 93.5 | 7.781 | -0.227 | 4.088 | 16.680 | 13.1 | 2.6 | 41.79 (1.54) |  |
|  | 1500 | 95.0 | 3.845 | 1.962 | -0.255 | 1.956 | 3.873 | 38.6 | 2.2 | 4.95 (0.08) | 96.5 | 5.430 | -0.456 | 2.479 | 6.321 | 38.2 | 2.2 | 72.28 (2.57) |  |
|  | 3000 | 97.0 | 2.720 | 1.388 | 0.113 | 1.303 | 1.702 | 72.4 | 0.1 | 6.78 (0.12) | 95.0 | 3.831 | -0.011 | 1.839 | 3.367 | 72.3 | 0.2 | 125.16 (3.89) |  |
| A5<br>(0.271) | 750 | 96.0 | 5.440 | 2.776 | 0.025 | 2.615 | 6.802 | 35.2 | 0.6 | 1.39 (0.02) | 94.5 | 7.758 | -0.215 | 4.096 | 16.738 | 35.4 | 0.6 | 40.09 (1.32) |  |
|  | 1500 | 97.0 | 3.834 | 1.956 | 0.183 | 1.814 | 3.309 | 57.8 | 1.8 | 3.10 (0.08) | 95.0 | 5.376 | 0.148 | 2.617 | 6.834 | 57.9 | 1.7 | 73.34 (2.43) |  |
|  | 3000 | 97.0 | 2.714 | 1.385 | 0.046 | 1.292 | 1.664 | 87.9 | 5.1 | 8.88 (0.12) | 95.0 | 3.812 | -0.016 | 1.899 | 3.587 | 87.8 | 5.1 | 139.04 (4.40) |  |
| A6<br>(0.377) | 750 | 96.5 | 5.447 | 2.779 | 0.041 | 2.740 | 7.471 | 23.8 | 1.9 | 2.42 (0.04) | 93.5 | 7.765 | -0.313 | 4.165 | 17.359 | 23.7 | 1.9 | 36.60 (1.48) |  |
|  | 1500 | 92.5 | 3.863 | 1.971 | 0.052 | 2.113 | 4.447 | 40.0 | 3.4 | 4.14 (0.10) | 95.5 | 5.466 | -0.208 | 2.830 | 8.011 | 40.1 | 3.4 | 64.18 (2.49) |  |
|  | 3000 | 95.5 | 2.735 | 1.396 | -0.024 | 1.388 | 1.918 | 62.2 | 7.2 | 8.34 (0.12) | 94.5 | 3.837 | -0.013 | 1.959 | 3.817 | 62.4 | 7.2 | 114.23 (3.68) |  |

For mediator selection, CF-OLS and B-Mixed had comparable performance when iSIS-MCP was used. Generally, a high average true positive rate was achieved when the sample size was 3000. In particular, we identified a substantial proportion of true mediators ***M***_𝒯_ in scenario (A1). Also, iSIS-MCP controlled the average false positive rate at a low level across all scenarios. The average false positive rate increased as the sample size increased in scenarios (A3), (A5), and (A6) for both methods because 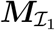 was associated with outcome *Y* given *X* and thus were not filtered out by iSIS. In Supplementary Materials Web Appendix B, we show that the average false positive rate was maintained at a low level after implementing the FDR control. However, inevitably, a small number of true mediators are excluded, as the primary aim of the FDR control is to minimize the false positive rate. Therefore, we highlight the trade-off between true positives (i.e., selecting true mediators) and false positives (i.e., falsely selecting non-mediators).

The empirical coverage probability using the CF-OLS method was satisfactory across all scenarios, and it yielded narrower confidence intervals than did the B-Mixed method. Meanwhile, we found that the empirical standard deviation of replicated estimations of CF-OLS (i.e., from its sampling distribution) was lower than that of B-Mixed. This is because the CF-OLS method makes full use of the two subsamples as illustrated in Figure 2 in contrast with the B-Mixed method, which conducts inference using only half of the data. In scenarios (A2), (A4), (A5), and (A6), we observed a relatively sizeable MSE for both methods when the sample size was 750 owing to over-selection of 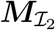 and under-selection of ***M***_𝒯_ by iSIS. The bias and MSE improved in all scenarios with increasing sample size.

Figure 3 displays asymptotic standard errors and the empirical standard deviation of replicated estimations using the CF-OLS method in scenarios (A1)–(A6). The asymptotic standard error is the mean value of 200 replications; the error bars in the figure represent one standard error of the mean. Generally, the asymptotic standard errors and empirical standard deviation tracked each other closely as the sample size increased from 500 to 3000.

**Figure 3:**
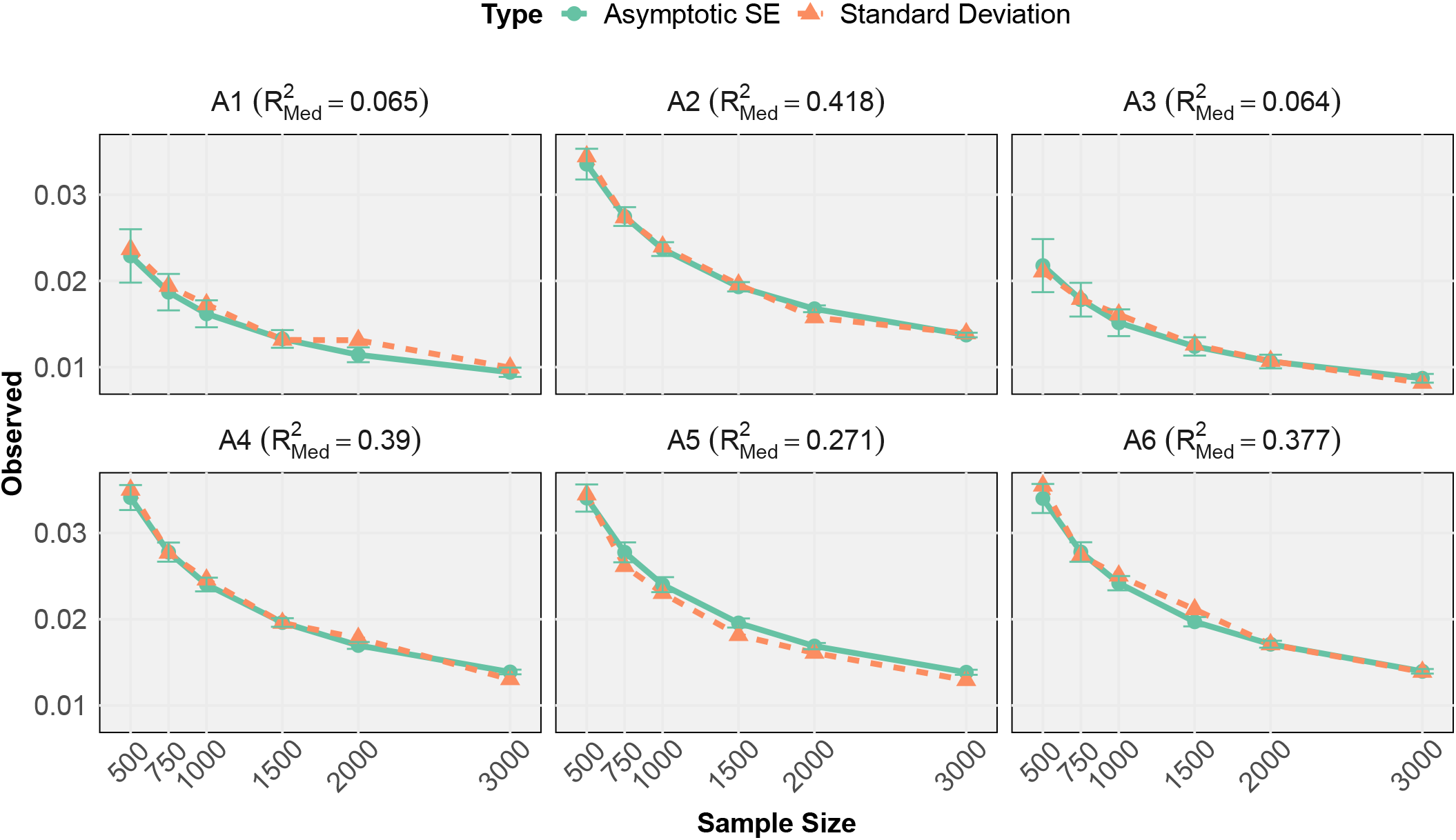
Plots of asymptotic standard error (green) and empirical standard deviation (orange) for 200 replicated estimations using the CF-OLS method for scenarios (A1)–(A6). SE refers to standard errors. The sample size increased from 500 to 3,000. The true value of 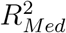 is listed within the parentheses. The error bars represent one standard error of the mean of asymptotic standard error across 200 replications in each scenario.

As expected, we observed a trend of decreasing asymptotic standard errors and empirical standard deviation with increasing sample size.

Importantly, in terms of computation, the CF-OLS method significantly outperformed the bootstrap-based B-Mixed method. Table 1 provides the means and standard errors of the computational time measured in minutes based on 200 replications using the CF-OLS and B-Mixed methods. For example, in scenario (A6) with a sample size of 750, CF-OLS spent about 2.42 minutes constructing one confidence interval using a single CPU core. In comparison, the B-Mixed method took about 36.6 minutes to achieve the same goal using 20 cores in parallel. For all the scenarios with a sample size of 3000, the proposed CF-OLS method shortened the time to compute the coverage probability based on 200 replications from longer than 380 hours to shorter than 30 hours. In practice, we found that the computational time with the B-Mixed method fluctuated highly but that with the CF-OLS method was quite stable. Of note is that the most time-consuming part of both methods was the variable selection step instead of the estimation step.

Table 2 demonstrates the robust performance of the CF-OLS method in handling correlated putative mediators across two distinct correlation structures. The true mediators ***M***_𝒯_ and the non-mediator 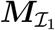 were correlated in scenarios (A7)–(A12). For mediator selection, the method consistently yielded a high average true positive rate while maintaining a low average false positive rate. Impressively, the empirical coverage probability remained favorable, even with sparse true mediators ***M***_𝒯_ and a limited sample size. In general, as the sample size increased from 500 to 3000, the asymptotic standard errors and empirical standard deviations mirrored each other closely. Consistent with expectations, both the asymptotic standard errors and empirical standard deviations exhibited a downward trend as the sample size increased.

**Table 2:** Simulation results using the CF-OLS method for correlated putative mediators in scenarios (A7)–(A12). **N** refers to the sample size. **CP** refers to coverage probability based on 200 replications. **Width** refers to half the width of the 95% confidence interval. **SE** refers to the average asymptotic standard error. **SD** refers to the empirical standard deviation of replicated estimations. MSE refers to mean squared error. **TP** refers to the average true positive rate. **FP** refers to the average false positive rate. The true value of 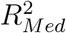 is shown in parentheses.

| Scenario | N | Correlation Structure 1 |  |  |  |  |  |  |  | Correlation Structure 2 |  |  |  |  |  |  |  |
| --- | --- | --- | --- | --- | --- | --- | --- | --- | --- | --- | --- | --- | --- | --- | --- | --- | --- |
|  |  | CP | Width | SE | Bias | SD | MSE | TP | FP | CP | Width | SE | Bias | SD | MSE | TP | FP |
| $(R^2_{Med})$ | | % | ($\times 10^{-2}$) | ($\times 10^{-2}$) | ($\times 10^{-2}$) | ($\times 10^{-2}$) | ($\times 10^{-2}$) | % | % | % | ($\times 10^{-2}$) | ($\times 10^{-2}$) | ($\times 10^{-2}$) | ($\times 10^{-2}$) | ($\times 10^{-2}$) | % | % |
| A7<br>(0) | 750 | 91.5 | 5.082 | 2.593 | 1.317 | 2.738 | 0.092 | \ | 1.3 | 91.5 | 4.942 | 2.522 | 1.313 | 2.697 | 0.090 | \ | 1.4 |
|  | 1500 | 93.0 | 3.489 | 1.780 | 0.876 | 1.946 | 0.045 | \ | 1.2 | 93.5 | 3.550 | 1.811 | 0.720 | 1.911 | 0.042 | \ | 1.3 |
|  | 3000 | 95.0 | 2.497 | 1.274 | 0.281 | 1.336 | 0.019 | \ | 1.2 | 98.0 | 2.494 | 1.272 | 0.177 | 1.146 | 0.013 | \ | 1.3 |
| A8<br>(0.128) | 750 | 95.0 | 5.667 | 2.891 | 0.455 | 2.732 | 0.076 | 100.0 | 0.0 | 93.0 | 5.162 | 2.634 | 0.037 | 2.811 | 0.079 | 100.0 | 0.0 |
|  | 1500 | 93.5 | 3.992 | 2.037 | -0.163 | 2.165 | 0.047 | 100.0 | 0.0 | 94.5 | 3.666 | 1.870 | -0.251 | 1.830 | 0.034 | 100.0 | 0.0 |
|  | 3000 | 94.5 | 2.830 | 1.444 | -0.059 | 1.484 | 0.022 | 100.0 | 0.0 | 94.5 | 2.600 | 1.327 | -0.250 | 1.299 | 0.017 | 100.0 | 0.0 |
| A9<br>(0.645) | 750 | 96.0 | 4.074 | 2.079 | -0.090 | 1.957 | 0.038 | 83.5 | 0.3 | 95.0 | 4.218 | 2.152 | -0.164 | 2.100 | 0.044 | 79.0 | 0.5 |
|  | 1500 | 96.0 | 2.878 | 1.469 | -0.147 | 1.445 | 0.021 | 86.0 | 1.6 | 95.5 | 2.991 | 1.526 | -0.028 | 1.498 | 0.022 | 79.2 | 3.6 |
|  | 3000 | 93.0 | 2.049 | 1.045 | -0.404 | 1.075 | 0.013 | 86.9 | 0.3 | 95.0 | 2.120 | 1.082 | -0.221 | 1.122 | 0.013 | 73.5 | 2.2 |
| A10<br>(0.315) | 750 | 95.0 | 5.439 | 2.775 | 0.089 | 2.960 | 0.087 | 86.4 | 2.9 | 93.5 | 5.462 | 2.787 | 0.304 | 2.849 | 0.082 | 83.5 | 2.6 |
|  | 1500 | 95.0 | 3.869 | 1.974 | -0.196 | 1.886 | 0.036 | 94.5 | 4.9 | 93.5 | 3.871 | 1.975 | 0.507 | 1.995 | 0.042 | 67.0 | 4.8 |
|  | 3000 | 93.5 | 2.749 | 1.403 | -0.087 | 1.462 | 0.021 | 95.0 | 4.3 | 95.5 | 2.742 | 1.399 | 0.205 | 1.316 | 0.018 | 62.4 | 3.3 |
| A11<br>(0.015) | 750 | 92.5 | 2.015 | 1.028 | 0.579 | 1.131 | 0.016 | 96.4 | 1.6 | 95.0 | 1.784 | 0.910 | 0.459 | 0.887 | 0.010 | 94.3 | 1.2 |
|  | 1500 | 94.5 | 1.428 | 0.729 | 0.334 | 0.764 | 0.007 | 96.8 | 1.4 | 95.0 | 1.278 | 0.652 | 0.314 | 0.706 | 0.006 | 94.5 | 0.3 |
|  | 3000 | 94.5 | 0.996 | 0.508 | 0.193 | 0.500 | 0.003 | 98.4 | 0.9 | 95.0 | 0.908 | 0.463 | 0.218 | 0.441 | 0.002 | 95.0 | 0.1 |
| A12<br>(0.003) | 750 | 95.5 | 1.057 | 0.539 | 0.533 | 0.613 | 0.007 | 73.4 | 3.0 | 93.5 | 1.167 | 0.596 | 0.542 | 0.576 | 0.006 | 65.2 | 2.7 |
|  | 1500 | 93.0 | 0.690 | 0.352 | 0.301 | 0.374 | 0.002 | 99.7 | 5.4 | 93.5 | 0.746 | 0.381 | 0.258 | 0.380 | 0.002 | 67.5 | 4.4 |
|  | 3000 | 97.5 | 0.464 | 0.237 | 0.140 | 0.247 | 0.001 | 98.6 | 5.1 | 96.5 | 0.492 | 0.251 | 0.080 | 0.261 | 0.001 | 60.2 | 3.4 |

As shown in Supplementary Materials Web Appendix B, we further evaluated the proposed CF-OLS method under scenarios (B1)–(B6) and (C1)–(C6). In scenarios (B1)–(B6), the regression coefficients ***α*** and ***β*** followed the uniform distribution *U* (−2, 2), and in scenarios (C1)–(C6), ***α*** and ***β*** followed the standard normal distribution *N* (0, 1^2^) when they were not set to 0. Overall, the coverage probability was satisfactory. When the sample size was 3000, the variable selection procedure captured an extensive number of true mediators ***M***_𝒯_, which gave a reasonable average true positive rate. Furthermore, the average false positive rate was controlled at a low level by eliminating most of the non-mediators 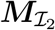. We also found that an increased average false positive rate resulted from the presence of the selected non-mediators 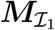 in scenarios (B3), (B5), (C3), and (C5). However, a promising finding was that the number of selected non-mediators 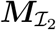 was still reasonably low, and the number of selected noise variables was nearly 0. As expected, we observed a smaller MSE with a larger sample size. Asymptotic standard errors approximated the empirical standard deviation of replicated estimations well for scenarios (B1)–(B6) and (C1)–(C6). In summary, the performance of CF-OLS under various settings was satisfactory in terms of mediator selection, coverage probability, and computational efficiency.

Additionally, we summarized the performance of the mean-based measures alongside the SOS measure across scenarios (A1) to (A12) in Supplementary Materials Web Appendix B. Overall, the bias and MSE of the SOS measure were comparable with those of the total effect measure 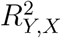 but were much lower than those of both the product and proportion measures. Importantly, in situations where the mediators were correlated and the number of true mediators was nonzero, the bias of the product and proportion measures deteriorated, whereas the SOS measure maintained a reasonable level of accuracy. Moreover, in Supplementary Materials Web Appendix C, we explored some alternative options for the iSIS procedure along with the CF-OLS method that may reduce the computational time and/or increase the accuracy of variable selection. We considered Lasso (Tibshirani, 1996), a popular alternative to MCP for sparse regression. Based on scenarios (A1)–(A6) in Table 1, we examined how our method performed with Lasso using the Akaike Information Criterion (AIC) (Akaike, 1998) for tuning the regularization parameter. We found that iSIS-Lasso kept the non-mediators 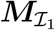 and noise variables 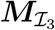 at levels similar to those for iSIS-MCP but failed to exclude the non-mediators 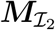. Unlike iSIS-MCP, model selection with iSIS-Lasso suffered from an increase in the average false positive rate as the sample size increased.

A possible reason for this is that Lasso regression tends to include an extensive number of false positives (Martinez et al., 2010). Despite this, we observed a minor discrepancy in the coverage probability and bias from those with iSIS-MCP using CF-OLS, which performed well across all scenarios.

## 4 Application to the Framingham Heart Study

Hypertension is a leading cause of cardiovascular disease (CVD) and mortality worldwide (Roth et al., 2018). Of the adult population worldwide in the year 2010, about 1.39 billion had hypertension, the primary symptom of which is persistently high BP, expressed as high systolic BP and diastolic BP (Mills et al., 2016). The prevalence of hypertension increases with chronological age, contributing to the current pandemic of CVD (Kearney et al., 2005). On the other hand, a higher plasma level of HDL-C was associated with a lower risk of coronary heart disease in several epidemiological studies (Castelli, 1988). A previous prospective cohort study demonstrated that the incidence and mortality of coronary heart disease among men were about threefold and fivefold greater than those among women, respectively, for which a difference in HDL-C level was the major determinant (Jousilahti et al., 1999). Our motivation was to investigate the effect of chronological age on systolic BP and the effect of sex on HDL-C level mediated by genome-wide gene expression.

We applied our proposed CF-OLS method to the individuals in the FHS Offspring Cohort who attended the 8^*th*^ and 9^*th*^ examinations and those in the FHS Third-Generation Cohort who attended the 2^*nd*^ and 3^*rd*^ examinations. BP was measured as the average value for two BP readings by physicians (to the nearest 2 mm Hg). Then BP was adjusted according to the intake of anti-hypertensive medication by adding 15 mm Hg to the measurements for treated individuals (Tobin et al., 2005). Also, HDL-C level was measured from the EDTA plasma (mg/dL) and age was recorded at the time the subject attended the examination. The covariates were body mass index (in *kg/m*^2^), smoking status (current smoker vs. current non-smoker), drinking status (never vs. ever), and the cohort the subject belonged to (Offspring Cohort vs. Generation 3 Cohort). We also incorporated the top 10 principal components (PCs) of genome-wide gene expression data, selected based on eigenvalues, as covariates in the mediation analysis models. The widespread use of PCs in genome-wide association studies underscores their importance, particularly in correcting for subtle population stratification and controlling for confounding genetic backgrounds (Price et al., 2006; Patterson et al., 2006). Age and sex were adjusted in the model, whereas the other one was considered the exposure variable of interest. High-throughput gene expression profiling of 17873 genes was performed from whole blood mRNA using an Affymetrix GeneChip Human Exon 1.0 ST (Joehanes et al., 2012). We extracted age, sex, covariates, and gene expression levels for the Offspring Cohort 8^*th*^ examination and Generation 3 Cohort 2^*nd*^ examination. Phenotypes were extracted from the Offspring Cohort 9^*th*^ examination and Generation 3 Cohort 3^*rd*^ examination, following the establishment by Kraemer et al. (2002) that the exposure affects the mediators which in turn precedes the outcome. We included a total of 4542 subjects with complete data in the systolic BP analysis and 4481 in the HDL-C analysis. For comparison, we followed Yang et al. (2021) by regressing covariates out from exposure, phenotypes, and gene expression levels to obtain the residuals for the following analyses to control for confounding effects. The descriptive statistics for the FHS samples are summarized in Supplementary Materials Web Appendix D.

The High Dimensional Multiple Testing (HDMT) method is designed to rigorously control for both the family-wise error rate (FWER) and the FDR in hypothesis testing of high-dimensional mediators (Dai et al., 2022). For comparison, we employed the HDMT method in lieu of the iSIS-MCP procedure to select variables in two subsamples independently, while keeping the inference process the same as that with the CF-OLS method. After eliminating the non-mediators 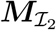, we applied the FDR control with a cutoff of 0.2 in each of the three methods to further filter out the non-mediators 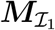. This is essential for gaining a deeper understanding of the underlying biological mechanism. We further applied the product and proportion measures based on the difference in means to the FHS data.

Table 3 compares the results of data analysis using the CF-OLS method, B-Mixed method, and HDMT methods. We found that the three methods provided comparable point estimation and confidence intervals, suggesting that the new CF-OLS method is able to provide reliable inferences. For the CF-OLS method, 20.1% of systolic BP variation could be explained by age, and 166 and 194 genes were selected in the two subsamples, respectively. Of note is that 12.6% (95% CI = (10.9%, 14.4%)) of the variance in systolic BP was attributable to the indirect effect of age through mediation by gene expression, resulting in an 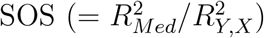 of 61.2% (95% CI = (55.9%, 66.6%)). Similarly, 16.6% of variance in HDL-C was explained by sex; 8.3% (95% CI = (6.9%, 9.8%)) of the variation was explained by sex through gene expression, with 107 and 110 genes selected in each of the two subsamples, leading to an SOS of 48.5% (95% CI = (42.1%, 54.9%)). We found that all three methods yielded similar results for the mean-based measures. However, for systolic BP, the indirect and total effects had opposite directions. This resulted in a negative value for the proportion measure, which is counterintuitive and difficult to interpret. For HDL-C level, the mean-based measures yielded interpretable results. Using the proportion measure with the CF-OLS method, we found that 55.0% and 53.6% of the total effect was mediated by gene expressions in the two subsamples, respectively. Indirect effect sizes of 8.58 and 8.05 indicated the expected change in the systolic BP for every unit increase in age mediated through the gene expression. These mean-based measures were also consistent across the CF-OLS, B-Mixed, and HDMT methods.

**Table 3:** Mediation effect sizes and their 95% confidence intervals estimated using the CF-OLS, B-Mixed, and HDMT methods with the Framingham Heart Study (FHS) data. **Exp** refers to the exposure variable. **N** refers to the sample size. **ab** refers to the indirect mediation effect. **prop** refers to the proportion measure. **total** refers to the total effect. 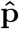 refers to the number of genes selected. The 95% confidence intervals (in parentheses) for the B-Mixed method were computed using 500 bootstrap samples. For the CF-OLS and HDMT methods, the splitting of data resulted in two sets of results for **ab, prop, total**, and 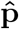 across two subsamples.

| Outcome | Exp | Method | $R^2_{Med}$ | SOS | $R^2_{Y,X}$ | ab | prop | total | $\hat{p}$ |
| --- | --- | --- | --- | --- | --- | --- | --- | --- | --- |
| Systolic BP<br>(N=4542) | Age | CF-OLS | 0.126 | 0.612 | 0.201 | -6.733/-7.052 | -10.094/-10.557 | 0.667/0.668 | 166/194 |
|  |  |  | (0.109, 0.144) | (0.559, 0.666) |  |  |  |  |  |
|  |  | B-Mixed | 0.120 | 0.601 | 0.200 | -7.268 | -10.877 | 0.668 | 200 |
|  |  |  | (0.081, 0.147) | (0.437, 0.705) | (0.174, 0.229) | (-8.108, -6.364) | (-13.177, -8.718) | (0.615, 0.730) | (149, 221) |
|  |  | HDMT | 0.042 | 0.205 | 0.201 | -6.852/-7.195 | -10.267/-10.770 | 0.667/0.668 | 7/11 |
|  |  |  | (0.034, 0.051) | (0.167, 0.243) |  |  |  |  |  |
| HDL-C<br>(N=4481) | Sex | CF-OLS | 0.083 | 0.485 | 0.166 | 8.580/8.051 | 0.550/0.536 | 15.613/15.024 | 107/110 |
|  |  |  | (0.069, 0.098) | (0.421, 0.549) |  |  |  |  |  |
|  |  | B-Mixed | 0.067 | 0.378 | 0.178 | 8.225 | 0.528 | 15.586 | 103 |
|  |  |  | (0.049, 0.169) | (0.285, 0.893) | (0.155, 0.263) | (7.282, 9.334) | (0.506, 0.553) | (14.402, 16.878) | (67, 134) |
|  |  | HDMT | 0.058 | 0.325 | 0.166 | 8.489/8.037 | 0.544/0.535 | 15.613/15.024 | 23/48 |
|  |  |  | (0.044, 0.068) | (0.265, 0.385) |  |  |  |  |  |

We further performed the canonical correlation analysis (CCA) (Harold, 1936) to evaluate the overlapping information for the two selected gene sets for each trait. More than 90% of the variance in canonical variates for systolic BP can be explained by the top eight canonical correlations. Similarly, more than 90% of the variance in canonical variates for HDL-C level can be captured by the top 12 canonical correlations. We also applied CCA to the genes identified by both the iSIS-MCP procedure and the HDMT method. Notably, even though the HDMT method was conservative in mediator selection, the top six canonical correlations still represented more than 90% of the variance in canonical variates for systolic BP. Meanwhile, the top 15 canonical correlations accounted for more than 90% of the variance in canonical variates for HDL-C level. In conclusion, regardless of whether genes were chosen from the two subsamples or via different variable selection methods, they largely captured similar biological information, likely at the pathway level, even though they did not exactly overlap. In our application to the FHS data, we also employed the CF-OLS and B-Mixed methods to assess the mediation effects for systolic BP exclusively within the FHS Offspring cohort. This approach allowed us to compare our findings with whose of prior research (Yang et al., 2021). The detailed results are included in the Supplementary Materials Table S15. Owing to the use of the full sample, the CF-OLS method yielded a narrower confidence interval than did the B-Mixed method, despite both methods yielding similar 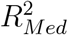 point estimates based on the OLS and linear mixed model, respectively. Specifically, the CF-OLS method attributed 4.29% (95% CI = (2.67%, 5.91%)) of the variance in systolic BP to the indirect effect of age mediated by gene expression. In contrast, the B-Mixed method’s estimate for the same mediation effect was 3.50% (95% CI = (−0.91%, 6.95%)).

To gain further insights into the mediating biological pathways, we performed pathway enrichment analysis of the selected mediating genes in all subsamples for systolic BP and HDL-C level. We identified five nominally significant pathways for systolic BP and five for HDL-C level, respectively. (See Supplementary Materials Web Appendix D). For example, rat and other studies demonstrated that the MAPK signaling pathway plays a mediatory role in the effect of the aging process on hypertension. The MAPK pathways, including extracellular signal-regulated kinase (ERK), c-Jun N-terminal Kinase (JNK), and p38 MAPK, are crucial to vascular aging and hypertension (Muslin, 2008). Aging is associated with MAPK activity in vascular tissues. Researchers showed that targeted inhibition of p38 MAPK promotes hypertrophic cardiomyopathy through upregulation of calcineurin-NFAT signaling (Braz et al., 2003). Also, oxidative stress, which increases with age, activates the MAPK pathway in endothelial cells, leading to endothelial dysfunction and a predisposition to hypertension (Son et al., 2011). The activation leads to a reduction in endothelial dependent vasodilation in humans, contributing to increased systolic BP (Seals et al., 2011). The B-Mixed method previously identified this pathway in Yang et al. (2021), underscoring the validity and efficiency of our proposed approach. Regarding the HDL-C outcome, we identified the cholesterol metabolism pathway, which encompasses the *CETP* (Cholesteryl Ester Transfer Protein) and *LDLR* (Low-Density Lipoprotein Receptor) genes. Authors reported that both *CETP* and *LDLR* were robustly associated with blood lipid levels in large-scale genome-wide association studies (Global Lipids Genetics Consortium, 2013). In addition, investigators showed that estrogen enhanced *LDLR* expression, facilitating the removal of Low-Density Lipoprotein (LDL) cholesterol from the bloodstream and thereby promoting cardiovascular health (Palmisano et al., 2018). Generally, higher *CETP* activity can lead to lower levels of HDL-C, reducing the size and number of the particles (Yamashita et al., 1991).

Finally, the computation time for CF-OLS to construct confidence intervals was substantially shorter than that for B-Mixed. In fact, the CF-OLS method can be 400 times faster than the B-Mixed method with the same computational resources. Specifically, finishing the analysis for systolic BP with CF-OLS using a single core took about 4.67 hours, whereas that with nonparametric bootstrap-based B-Mixed using 25 cores in parallel took around 75.99 hours. For the HDL-C outcome analysis, finishing the analysis with the CF-OLS method using a single core took about 5.19 hours, whereas finishing it with the B-Mixed method using 25 cores in parallel took about 54.70 hours.

## 5 Discussion

We proposed a novel two-stage interval estimation procedure for 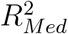 based on cross-fitting and sample-splitting to estimate the total mediation effect for high-dimensional mediators. Unlike the estimation method using nonparametric bootstrap in a mixed model framework, our proposed method relies on the asymptotic distribution of 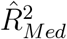 to construct confidence intervals. After splitting the data into two subsamples, we estimated 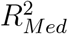 using OLS regression and conducted inference based on the asymptotic standard error. We excluded the non-mediators 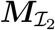 using iSIS-MCP in two subsamples separately and fitted OLS regression in the other subsample. As an optional but potentially beneficial step, we employed FDR control to further refine our list of potential mediators by excluding the non-mediators 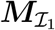.

Although Theorem 1 holds under the specific assumption on the conditional correlation of mediators and strength of spurious mediators, we found both in the simulation study and real data application, as shown in the Supplementary Materials, we found that the results did not change significantly with moderate conditional correlation and without further filtering of the non-mediators 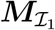. In practical settings, we rely on existing knowledge to identify confounders. However, it implicitly assumes that covariates are known and that the observed covariates adequately represent all existing confounders. In the context of high-dimensional gene expression data, confounders could be unknown or have various sources, leading to potential violation of the identifiability assumptions for causal mediation analysis as stated in Section 2.1 and elaborated on previously (Imai et al., 2010; Jérolon et al., 2020; VanderWeele et al., 2014; VanderWeele and Vansteelandt, 2009). For example, the role of 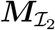 is usually unknown, and it can be considered a special type of post-treatment confounders when conditional residual correlation exists. Technical variables or batch effects are known to be difficult to correct (Leek et al., 2010), leading to the violation of the identifiability assumption. In our real data application, we performed variable selection to exclude 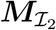 and adjusted for principal components that can be used to control for unknown confounding effects Yuan and Qu (2023). We observed much weaker residual correlation after such adjustment (Figure S3 and S4). More sophisticated methods are beyond the scope of the present study but are important topics for future work.

In addition, the point estimation improved over the original point estimation method described by Yang et al. (2021) in terms of the MSE because the new method used full data for variable selection and estimation demonstrated by our extensive simulation studies in Table 1. The CF-OLS method had narrower confidence intervals, comparable coverage probability and variable selection accuracy across various scenarios when compared with the B-Mixed method while significantly reducing the computational time. When we used iSIS-Lasso for mediator selection, the coverage probability was reasonable, but the false positive rate in some scenarios increased owing to failure in excluding the non-mediators 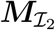.

In the FHS data analysis, treating systolic BP and HDL-C as outcomes, we applied the CF-OLS, B-Mixed, and HDMT methods to examination of the mediatory role of gene expression between exposure and phenotype. As established previously (Yang et al., 2021), a large amount of systolic BP variation can be explained by age through gene expression. In addition, we discovered that the effect of sex on HDL-C was mediated by gene expression.

Similar conclusions can be drawn after comparing the 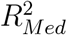 and its confidence intervals from the three methods, which corroborates the validity of the CF-OLS method. More importantly, and as expected, the CF-OLS method is very computationally efficient because it only performs the iSIS variable selection procedure twice to construct confidence intervals instead of 500 times as in the resampling-based B-Mixed method. To compute the confidence interval for systolic BP in the FHS dataset, the B-Mixed method took about 76 hours even with multicore parallel computing, whereas the CF-OLS method achieved it efficiently in about 4.5 hours using a single core. This advantage makes the CF-OLS method more practical in estimating the total mediation effect with confidence intervals under the high-dimensional setting and a relatively massive data set.

A critical research area in public health is how an exposure influences phenotypic variation. Authors have well established that exposures, including environmental (Bind et al., 2014; Timms et al., 2016), socioeconomic (Cerutti et al., 2021), and behavioral (Zong et al., 2019; Hardy and Tollefsbol, 2011; Tiffon, 2018; Maas et al., 2020) factors, are associated with changes at the molecular level (Bind et al., 2014; Timms et al., 2016; Maas et al., 2020; Huang et al., 2018; Tobi et al., 2018). Mediation analysis is a useful tool for decomposing the relationship between an exposure and an outcome into direct and mediation (indirect) effects. Over the past 3 decades, researchers have performed mediation analyses to extensively study settings in which a single mediator or a few mediators are present (Zeng et al., 2021). These methods are not generally applicable to high-dimensional molecular mediators. In the present study, we focused on the important but less explored total mediation effect, which captures the variations in outcome explained by an exposure through high-dimensional mediators. Accurate estimation of the total mediation effect improves understanding of the mediatory roles of genomic factors in various ways, including exploring the impact of a certain molecular phenotype in the exposure-outcome pathway, identifying relevant tissues or cell types, and improving the understanding of the time-varying mediatory role of a molecular phenotype. In addition to deepening our understanding of the biological mechanism at the molecular level, estimating the total mediation effect has the potential to guide outcome prediction and intervention. For example, incorporating mediators has benefited the prediction of survival outcomes (Zhou et al., 2022). Also, Tingley et al. (2014) suggested that refining interventions targeting the mechanism that explains a large proportion of an intervention’s effect on the outcome may be more desirable than the ones that do not.

The proposed method is available in `CFR2M` package on R/`CRAN`, which includes the new CF-OLS method. Lastly, whereas we have focused on continuous outcomes, we will extend our proposed approach to accommodate time-to-event and binary outcomes in the future (Chi et al., 2024).

## 6 Supplementary Materials

The Supplementary Materials contain technical proofs and additional numerical results. The proposed CF-OLS method is implemented in the R package `CFR2M`, which is publicly available on Github at https://github.com/zhichaoxu04/CFR2M. The R code for simulation and real data application is also available at https://github.com/zhichaoxu04/CFR2M-paper. The R package `CFR2M` is also contained in the updated R package `RsqMed` on `CRAN`.

## 8 Acknowledgments

This research was supported by National Institutes of Health (NIH) grant R01HL116720 (to P.W.). T.Y. was supported by the Children’s Cancer Research fund and a St Baldrick’s Career Award. The FHS was conducted and supported by the National Heart, Lung, and Blood Institute (NHLBI) in collaboration with Boston University (contract numbers N01-HC-25195, HHSN268201500001I, and 75N92019D00031). This manuscript was not prepared in collaboration with investigators in the FHS and does not necessarily reflect the opinions or views of the FHS, Boston University, or the NHLBI. The data set used for the analyses described in this manuscript was obtained from dbGaP at https://www.ncbi.nlm.nih.gov/gap/ through accession number phs000007. We acknowledge the support of the High Performance Computing for research facility at the University of Texas MD Anderson Cancer Center for providing computational resources that have contributed to the research results reported herein. We would like to thank Mr. Donald Norwood from the Research Medical Library at MD Anderson Cancer Center for editorial assistance. We are grateful to the two anonymous reviewers for their many constructive comments, which have helped substantially improve the presentation of this paper.

