## Supplementary material for "Speeding up interval estimation for *R*^2^-based mediation effect of high-dimensional mediators via cross-fitting": Supp

**Abstract**

This supplementary material contains additional information to better understand the paper and has the following sections. Web Appendix A provides technical conditions and the proof of Theorem 1. Web Appendix B gives simulation details under high dimensional settings using the CF-OLS method. In Web Appendix C, we compare the performance of various iSIS settings, and in Web Appendix D, we conduct a pathway enrichment analysis for systolic BP and HDL-C.

### 1 Web Appendix A: Technical conditions and proof of Theorem 1

#### 1.1 Technical conditions

For the proof of Theorem 1, we introduce two further assumptions beyond Assumption 1, as described in the main text. Note that when nonzero signals $\gtrsim \sqrt{\log(p)/n}$, the oracle property is achievable, and the theoretical analysis of the oracle case is straightforward (see Proof of Theorem 1, Analysis of oracle estimator). Thus, our analysis shall focus on the case where most nonzero signals $\lesssim \sqrt{\log(p)/n}$, as given by Assumption 2. In this regime, exact selection may not be possible according to the information-theoretic limit.

**Assumption 2.** Suppose $|\alpha_j| \lesssim \sqrt{\log(p)/n}$ and $|\beta_j| \lesssim \sqrt{\log(p)/n}$ for $j \notin \mathcal{T}$.

We have not explicitly imposed assumptions on $\{|\beta_j| : j \in \mathcal{T}\}$; however, to achieve the sure screening property (Assumption 1), it roughly requires $|\alpha_j|, |\beta_j| \gtrsim \sqrt{\log(p)/n}$ for $j \in \mathcal{T}$.

*Correspondence to T. Yang and P. Wei. +These authors contributed equally to this work. [1]Department of Biostatistics, The University of Texas MD Anderson Cancer Center, Houston, Texas 77030, U.S.A. [2]Department of Statistics, Iowa State University, Ames, Iowa, 50011, U.S.A. [3]Division of Biostatistics, School of Public Health, University of Minnesota, Minneapolis, Minnesota 55455, U.S.A.

**Assumption 3.** Suppose $\max\{|\Sigma_{kj}| : k \in \mathcal{T}, j \in \mathcal{T}^c\} \lesssim \sqrt{\log(p)/n}$ and $c_1 \leq \lambda_{\min}(\mathbf{\Sigma}) \leq \lambda_{\max}(\mathbf{\Sigma}) \leq c_2$, where $\mathbf{\Sigma}$ is the covariance of $\boldsymbol{\xi}$.

Assumption 3 is a regularity condition on $\mathbf{\Sigma}$, requiring that $\boldsymbol{\xi}$ is not too correlated. From the causal mediation analysis perspective, we note that correlated $\boldsymbol{\xi}$ suggests a violation of the parallel mediators assumption, which suggests the existence of uncontrolled confounding effects [Yuan and Qu, 2023]. It is thus reasonable to derive the asymptotic properties of $R^2_{Med}$ under Assumption 3.

**Relaxing normality assumption.** We have assumed the variables are jointly normal to avoid technicality and to improve the presentation. This assumption, however, is unnecessary. In fact, any conditional expectation $\mathrm{E}(\cdot \mid \star)$ in our derivation can be replaced with the "best linear approximation" operator $\mathbb{L}(\cdot \mid \star)$, defined as follows. Given random variables $U$ and $\boldsymbol{W}$, let $\mathbb{L}(U \mid \boldsymbol{W})$ be the best linear approximation of $U$ using $\boldsymbol{W}$, namely $\mathbb{L}(U \mid \boldsymbol{W}) = \widetilde{\boldsymbol{\theta}}^\top \boldsymbol{W}$ where

$$\widetilde{\boldsymbol{\theta}} = \arg\min_{\boldsymbol{\theta}} \ \mathrm{E}(U - \boldsymbol{\theta}^\top \boldsymbol{W})^2.$$

For random variables $U$, $U'$, and $\boldsymbol{W}$, we have that (a) $\mathbb{L}(U + U' \mid \boldsymbol{W}) = \mathbb{L}(U \mid \boldsymbol{W}) + \mathbb{L}(U' \mid \boldsymbol{W})$, (b) $\mathbb{L}(cU \mid \boldsymbol{W}) = c\mathbb{L}(U \mid \boldsymbol{W})$ for $c \in \mathbb{R}$, (c) $\mathbb{L}(U \mid \boldsymbol{W}) = 0$ if $\mathrm{Cov}(U, \boldsymbol{W}) = \mathbf{0}$, (d) $\mathbb{L}(U \mid \boldsymbol{W}) = U$ if $U \in \mathrm{Span}(\boldsymbol{W})$, and (e) $\mathbb{L}(U \mid \boldsymbol{W}) = \mathbb{L}(U \mid \mathbf{A}\boldsymbol{W})$ for invertible $\mathbf{A}$. Thus, $\mathbb{L}(\cdot \mid \star)$ mimics $\mathbb{E}(\cdot \mid \star)$. As a result, if the data are sub-Gaussian, the proof of Theorem 1 continues to hold with $\mathrm{E}(\cdot \mid \star)$ being replaced by $\mathbb{L}(\cdot \mid \star)$.

#### 1.2 Proof of Theorem 1

Before proceeding, we first introduce some notations used in the proof. Recalling Equation (1) in the main text, we have

$$\boldsymbol{M} = \boldsymbol{\alpha} X + \boldsymbol{\xi}, \qquad Y = \gamma X + \boldsymbol{\beta}_{\mathcal{T}}^\top \boldsymbol{M}_{\mathcal{T}} + \boldsymbol{\beta}_{\mathcal{I}_1}^\top \boldsymbol{M}_{\mathcal{I}_1} + \varepsilon.$$

Let $\mathcal{A}$ denote a generic subset $\mathcal{T} \subseteq \mathcal{A} \subseteq \{1, \ldots, p\}$ such that $\mathcal{A} \subseteq \mathcal{T} \cup \mathcal{I}_1$ and $\mathcal{A} \subseteq \mathcal{T} \cup \mathcal{I}_2$. Define

$$\begin{aligned}
\eta &= \varepsilon + \boldsymbol{\beta}^\top \{\boldsymbol{M} - \mathrm{E}(\boldsymbol{M} \mid X)\}, \\
\omega_{\mathcal{A}} &= \varepsilon + \boldsymbol{\beta}_{\mathcal{I}_1}^\top \big\{\boldsymbol{M}_{\mathcal{I}_1} - \mathrm{E}(\boldsymbol{M}_{\mathcal{I}_1} \mid X, \boldsymbol{M}_{\mathcal{A}})\big\}, \\
\zeta_{\mathcal{A}} &= \gamma\big\{X - \mathrm{E}(X \mid \boldsymbol{M}_{\mathcal{A}})\big\} + \varepsilon + \boldsymbol{\beta}_{\mathcal{I}_1}^\top \big\{\boldsymbol{M}_{\mathcal{I}_1} - \mathrm{E}(\boldsymbol{M}_{\mathcal{I}_1} \mid \boldsymbol{M}_{\mathcal{A}})\big\}.
\end{aligned}$$

Let $\boldsymbol{\eta} = (\eta_1, \ldots, \eta_n)$, $\widehat{\boldsymbol{\eta}} = (\widehat{\eta}_1, \ldots, \widehat{\eta}_n)$, $\boldsymbol{\zeta}_{\mathcal{A}} = (\zeta_{\mathcal{A},1}, \ldots, \zeta_{\mathcal{A},n})$, $\widehat{\boldsymbol{\zeta}}_{\mathcal{A}} = (\widehat{\zeta}_{\mathcal{A},1}, \ldots, \widehat{\zeta}_{\mathcal{A},n})$, $\boldsymbol{\omega}_{\mathcal{A}} = (\omega_{\mathcal{A},1}, \ldots, \omega_{\mathcal{A},n})$, and $\widehat{\boldsymbol{\omega}}_{\mathcal{A}} = (\widehat{\omega}_{\mathcal{A},1}, \ldots, \widehat{\omega}_{\mathcal{A},n})$, where $(\eta_i, \zeta_{\mathcal{A},i}, \omega_{\mathcal{A},i})$ are independent and identically distributed copies of $(\eta, \zeta_{\mathcal{A}}, \omega_{\mathcal{A}})$, and $\widehat{\eta}_i$, $\widehat{\zeta}_{\mathcal{A},i}$, and $\widehat{\omega}_{\mathcal{A},i}$ are the residuals of OLS regressions of $Y$ over $X$, over $\boldsymbol{M}_{\mathcal{A}}$, and over $(X, \boldsymbol{M}_{\mathcal{A}})$. Further, we denote $\boldsymbol{y} = (Y_1, \ldots, Y_n)$ and define

$$\rho^2_{\mathcal{A}} = 1 - \Big(\mathrm{E}\,\eta^2 + \mathrm{E}\,\zeta^2_{\mathcal{A}} - \mathrm{E}\,\omega^2_{\mathcal{A}}\Big)/\,\mathrm{E}\,Y^2,$$

$$\widetilde{\rho}^2_{\mathcal{A}} = 1 - \Big(\frac{\boldsymbol{\eta}^\top\boldsymbol{\eta}}{n} + \frac{\boldsymbol{\zeta}^\top_{\mathcal{A}}\boldsymbol{\zeta}_{\mathcal{A}}}{n} - \frac{\boldsymbol{\omega}^\top_{\mathcal{A}}\boldsymbol{\omega}_{\mathcal{A}}}{n}\Big)/\Big(\frac{\boldsymbol{y}^\top\boldsymbol{y}}{n}\Big),$$

$$\widehat{\rho}^2_{\mathcal{A}} = 1 - \Big(\frac{\widehat{\boldsymbol{\eta}}^\top\widehat{\boldsymbol{\eta}}}{n} + \frac{\widehat{\boldsymbol{\zeta}}^\top_{\mathcal{A}}\widehat{\boldsymbol{\zeta}}_{\mathcal{A}}}{n} - \frac{\widehat{\boldsymbol{\omega}}^\top_{\mathcal{A}}\widehat{\boldsymbol{\omega}}_{\mathcal{A}}}{n}\Big)/\Big(\frac{\boldsymbol{y}^\top\boldsymbol{y}}{n}\Big),$$

As a result, we have $R^2_{Med} = \rho^2_{\mathcal{A}=\mathcal{T}}$. We simplify the notation of $\rho^2_{\mathcal{A}=\mathcal{T}}$ as $\rho^2_{\mathcal{T}}$.

Now, we outline the strategy for establishing the asymptotic distribution of $\widehat{R}^2_{Med}$. Firstly, we control the difference $|\widehat{\rho}^2_{\mathcal{A}} - \widetilde{\rho}^2_{\mathcal{A}}|$ uniformly in $\mathcal{A}$. Secondly, we upper-bound $|\widetilde{\rho}^2_{\mathcal{A}} - \widetilde{\rho}^2_{\mathcal{T}}|$ uniformly in $\mathcal{A}$. Thirdly, we derive the asymptotics for the oracle estimator $\widetilde{\rho}^2_{\mathcal{T}}$. Finally, we establish the large-sample properties of $\widehat{R}^2_{Med}$ estimated by our proposed algorithm.

**Bounding the difference between $\widehat{\rho}^2_{\mathcal{A}}$ and $\widetilde{\rho}^2_{\mathcal{A}}$.** Note that

$$\widehat{\boldsymbol{\eta}}^\top\widehat{\boldsymbol{\eta}} = \boldsymbol{\eta}^\top\boldsymbol{\eta} - \boldsymbol{\eta}^\top\boldsymbol{P}_X\boldsymbol{\eta},$$
$$\widehat{\boldsymbol{\zeta}}^\top_{\mathcal{A}}\widehat{\boldsymbol{\zeta}}_{\mathcal{A}} = \boldsymbol{\zeta}^\top_{\mathcal{A}}\boldsymbol{\zeta}_{\mathcal{A}} - \boldsymbol{\zeta}^\top_{\mathcal{A}}\boldsymbol{P}_{\boldsymbol{M}_{\mathcal{A}}}\boldsymbol{\zeta}_{\mathcal{A}},$$
$$\widehat{\boldsymbol{\omega}}^\top_{\mathcal{A}}\widehat{\boldsymbol{\omega}}_{\mathcal{A}} = \boldsymbol{\omega}^\top_{\mathcal{A}}\boldsymbol{\omega}_{\mathcal{A}} - \boldsymbol{\omega}^\top_{\mathcal{A}}\boldsymbol{P}_{X,\boldsymbol{M}_{\mathcal{A}}}\boldsymbol{\omega}_{\mathcal{A}},$$

where $\boldsymbol{P}_X$, $\boldsymbol{P}_{\boldsymbol{M}_{\mathcal{A}}}$, and $\boldsymbol{P}_{X,\boldsymbol{M}_{\mathcal{A}}}$ are the projection matrices onto the column spaces of $X$, $\boldsymbol{M}_{\mathcal{A}}$, and $(X, \boldsymbol{M}_{\mathcal{A}})$, respectively. Thus, we have

$$|\widehat{\rho}^2_{\mathcal{A}} - \widetilde{\rho}^2_{\mathcal{A}}| \lesssim \frac{1}{n}\left(\boldsymbol{\eta}^\top\boldsymbol{P}_X\boldsymbol{\eta} + \boldsymbol{\zeta}^\top_{\mathcal{A}}\boldsymbol{P}_{\boldsymbol{M}_{\mathcal{A}}}\boldsymbol{\zeta}_{\mathcal{A}} + \boldsymbol{\omega}^\top_{\mathcal{A}}\boldsymbol{P}_{X,\boldsymbol{M}_{\mathcal{A}}}\boldsymbol{\omega}_{\mathcal{A}}\right).$$

By Theorem 2.1 of Hsu et al. [2012], there exists a constant $C > 0$ such that we have

$$\mathbb{P}\Big(\boldsymbol{\eta}^\top\boldsymbol{P}_X\boldsymbol{\eta} \ge C(1 + 2\sqrt{t} + 2t)\Big) \le \exp(-t),$$
$$\mathbb{P}\Big(\boldsymbol{\zeta}^\top_{\mathcal{A}}\boldsymbol{P}_{\boldsymbol{M}_{\mathcal{A}}}\boldsymbol{\zeta}_{\mathcal{A}} \ge C(|\mathcal{A}| + \sqrt{|\mathcal{A}|t} + 2t)\Big) \le \exp(-t),$$
$$\mathbb{P}\Big(\boldsymbol{\omega}^\top_{\mathcal{A}}\boldsymbol{P}_{X,\boldsymbol{M}_{\mathcal{A}}}\boldsymbol{\omega}_{\mathcal{A}} \ge C(|\mathcal{A}| + \sqrt{|\mathcal{A}|t} + 2t)\Big) \le \exp(-t),$$

for any $\mathcal{A}$ with $|\mathcal{A}| \le s$. Consequently, if $t = s\log(p)$ and if $s\log(p) \ll n$, we have

$$\sqrt{n}\sup_{\mathcal{A}}|\widehat{\rho}^2_{\mathcal{A}} - \widetilde{\rho}^2_{\mathcal{A}}| \lesssim s\log(p)/\sqrt{n}.$$

This provides a uniform bound of $|\widehat{\rho}^2_{\mathcal{A}} - \widetilde{\rho}^2_{\mathcal{A}}|$ for any $|\mathcal{A}| \le s$.

**Bounding the difference between $\widetilde{\rho}^2_{\mathcal{A}}$ and $\widetilde{\rho}^2_{\mathcal{T}}$.** Note that

$$\begin{aligned}\sqrt{n}|\widetilde{\rho}^2_{\mathcal{A}} - \widetilde{\rho}^2_{\mathcal{T}}| &\lesssim \frac{1}{\sqrt{n}}\left|(\boldsymbol{\zeta}^\top_{\mathcal{A}}\boldsymbol{\zeta}_{\mathcal{A}} - \boldsymbol{\omega}^\top_{\mathcal{A}}\boldsymbol{\omega}_{\mathcal{A}}) - (\boldsymbol{\zeta}^\top_{\mathcal{T}}\boldsymbol{\zeta}_{\mathcal{T}} - \boldsymbol{\omega}^\top_{\mathcal{T}}\boldsymbol{\omega}_{\mathcal{T}})\right| \\ &\le \frac{1}{\sqrt{n}}\left|\boldsymbol{\zeta}^\top_{\mathcal{A}}\boldsymbol{\zeta}_{\mathcal{A}} - \boldsymbol{\zeta}^\top_{\mathcal{T}}\boldsymbol{\zeta}_{\mathcal{T}}\right| + \frac{1}{\sqrt{n}}\left|\boldsymbol{\omega}^\top_{\mathcal{A}}\boldsymbol{\omega}_{\mathcal{A}} - \boldsymbol{\omega}^\top_{\mathcal{T}}\boldsymbol{\omega}_{\mathcal{T}}\right|.\end{aligned}$$

To establish its upper bound, denoting

$$D_1 = \frac{1}{\sqrt{n}}\sum_{i=1}^{n}(\zeta_{\mathcal{A},i}^2 - \zeta_{\mathcal{T},i}^2), \qquad D_2 = \frac{1}{\sqrt{n}}\sum_{i=1}^{n}(\omega_{\mathcal{A},i}^2 - \omega_{\mathcal{T},i}^2),$$

we would like to show $\mathrm{E}(D_1) = o(1)$, $\mathrm{Var}(D_1) = o(1)$, $\mathrm{E}(D_2) = o(1)$, and $\mathrm{Var}(D_2) = o(1)$. To this end, consider an independent observation, $(\zeta_{\mathcal{A}}, \zeta_{\mathcal{T}}, \omega_{\mathcal{A}}, \omega_{\mathcal{T}})$. We have

$$\begin{aligned}\mathrm{E}(D_1) &= \sqrt{n}\,\mathrm{E}(\zeta_{\mathcal{A}}^2 - \zeta_{\mathcal{T}}^2), \quad \mathrm{Var}(D_1) = \mathrm{Var}(\zeta_{\mathcal{A}}^2 - \zeta_{\mathcal{T}}^2),\\ \mathrm{E}(D_2) &= \sqrt{n}\,\mathrm{E}(\omega_{\mathcal{A}}^2 - \omega_{\mathcal{T}}^2), \quad \mathrm{Var}(D_2) = \mathrm{Var}(\omega_{\mathcal{A}}^2 - \omega_{\mathcal{T}}^2).\end{aligned}$$

For $\mathrm{E}(D_1)$, we have

$$\mathrm{E}(D_1) = \sqrt{n}\gamma^2(v_1 - v_2) + \sqrt{n}\boldsymbol{\beta}_{\mathcal{I}_1}^\top(\boldsymbol{O}_1 - \boldsymbol{O}_2)\boldsymbol{\beta}_{\mathcal{I}_1},$$

where

$$\begin{aligned}v_1 &= \mathrm{E}\left\{X - \mathrm{E}(X \mid \boldsymbol{M}_{\mathcal{A}})\right\}^2,\\ v_2 &= \mathrm{E}\left\{X - \mathrm{E}(X \mid \boldsymbol{M}_{\mathcal{T}})\right\}^2,\\ \boldsymbol{O}_1 &= \mathrm{E}\left\{X - \mathrm{E}(X \mid \boldsymbol{M}_{\mathcal{A}})\right\}\left\{X - \mathrm{E}(X \mid \boldsymbol{M}_{\mathcal{A}})\right\}^\top,\\ \boldsymbol{O}_2 &= \mathrm{E}\left\{X - \mathrm{E}(X \mid \boldsymbol{M}_{\mathcal{T}})\right\}\left\{X - \mathrm{E}(X \mid \boldsymbol{M}_{\mathcal{T}})\right\}^\top.\end{aligned}$$

Note that

$$\sqrt{n}(v_1 - v_2) = \frac{\sigma_X^2\boldsymbol{\alpha}_{\mathcal{A}}^\top(\boldsymbol{\Sigma}_{\mathcal{A}\mathcal{A}}^{-1} - \widetilde{\boldsymbol{\Sigma}}_{\mathcal{A}\mathcal{A}}^{-1})\boldsymbol{\alpha}_{\mathcal{A}}}{(1 + \boldsymbol{\alpha}_{\mathcal{T}}^\top\boldsymbol{\Sigma}_{\mathcal{T}\mathcal{T}}^{-1}\boldsymbol{\alpha}_{\mathcal{T}})(1 + \boldsymbol{\alpha}_{\mathcal{A}}^\top\boldsymbol{\Sigma}_{\mathcal{A}\mathcal{A}}^{-1}\boldsymbol{\alpha}_{\mathcal{A}})} \lesssim \|\boldsymbol{\alpha}_{\mathcal{B}}\|_2^2 \lesssim s\log(p)/\sqrt{n} = o(1),$$

where $\widetilde{\boldsymbol{\Sigma}}_{\mathcal{A}\mathcal{A}}^{-1}$ is the Moore-Penrose inverse of block diagonal matrix $\mathrm{Diag}(\boldsymbol{\Sigma}_{\mathcal{T}\mathcal{T}}, \boldsymbol{0})$, $\boldsymbol{\alpha}_{\mathcal{A}} = (\boldsymbol{\alpha}_{\mathcal{T}}, \boldsymbol{\alpha}_{\mathcal{B}})$, and $\mathcal{B} = \mathcal{A} \setminus \mathcal{T}$. Also, note that

$$\sqrt{n}\boldsymbol{\beta}_{\mathcal{I}_1}^\top(\boldsymbol{O}_1 - \boldsymbol{O}_2)\boldsymbol{\beta}_{\mathcal{I}_1} \lesssim \|\boldsymbol{\beta}_{\mathcal{I}_1}\|_2^2 \lesssim s\log(p)/\sqrt{n} = o(1).$$

Thus, $\mathrm{E}(D_1) = o(1)$.

For $\mathrm{Var}(D_1)$, we have

$$\mathrm{Var}(D_1) \le \mathrm{E}\{(\zeta_{\mathcal{A}} + \zeta_{\mathcal{T}})^2(\zeta_{\mathcal{A}} - \zeta_{\mathcal{T}})^2\} \lesssim \mathrm{E}(\zeta_{\mathcal{A}} - \zeta_{\mathcal{T}})^2.$$

Note that

$$|\zeta_{\mathcal{A}} - \zeta_{\mathcal{T}}| \le \gamma|\,\mathrm{E}(X \mid \boldsymbol{M}_{\mathcal{A}}) - \mathrm{E}(X \mid \boldsymbol{M}_{\mathcal{T}})| + |\boldsymbol{\beta}_{\mathcal{I}_1}^\top\{\mathrm{E}(\boldsymbol{M}_{\mathcal{I}_1} \mid \boldsymbol{M}_{\mathcal{A}}) - \mathrm{E}(\boldsymbol{M}_{\mathcal{I}_1} \mid \boldsymbol{M}_{\mathcal{T}})\}|,$$

where

$$\begin{aligned}\mathrm{E}\,|\,\mathrm{E}(X \mid \boldsymbol{M}_{\mathcal{A}}) - \mathrm{E}(X \mid \boldsymbol{M}_{\mathcal{T}})|^2 &\le \|\boldsymbol{\alpha}_{\mathcal{I}_2}\|_2^2 \lesssim s\log(p)/n = o(1),\\ \mathrm{E}\,|\boldsymbol{\beta}_{\mathcal{I}_1}^\top\{\mathrm{E}(\boldsymbol{M}_{\mathcal{I}_1} \mid \boldsymbol{M}_{\mathcal{A}}) - \mathrm{E}(\boldsymbol{M}_{\mathcal{I}_1} \mid \boldsymbol{M}_{\mathcal{T}})\}|^2 &\lesssim \|\boldsymbol{\beta}_{\mathcal{I}_1}\|_2^2 \lesssim s\log(p)/n = o(1).\end{aligned}$$

Therefore, $\mathrm{Var}(D_1) = o(1)$.

Similarly, for $\mathrm{E}(D_2)$ we have

$$\mathrm{E}(D_2) = \sqrt{n}\boldsymbol{\beta}_{\mathcal{I}_1}^\top(\boldsymbol{Q}_1 - \boldsymbol{Q}_2)\boldsymbol{\beta}_{\mathcal{I}_1} \lesssim \|\boldsymbol{\beta}_{\mathcal{I}_1}\|_2^2 \lesssim s\log(p)/\sqrt{n} = o(1),$$

where

$$\boldsymbol{Q}_1 = \mathrm{E}\left\{\boldsymbol{M}_{\mathcal{I}_1} - \mathrm{E}(\boldsymbol{M}_{\mathcal{I}_1} \mid X, \boldsymbol{M}_{\mathcal{A}})\right\}\left\{\boldsymbol{M}_{\mathcal{I}_1} - \mathrm{E}(\boldsymbol{M}_{\mathcal{I}_1} \mid X, \boldsymbol{M}_{\mathcal{A}})\right\}^\top,$$
$$\boldsymbol{Q}_2 = \mathrm{E}\left\{\boldsymbol{M}_{\mathcal{I}_1} - \mathrm{E}(\boldsymbol{M}_{\mathcal{I}_1} \mid X, \boldsymbol{M}_{\mathcal{A}})\right\}\left\{\boldsymbol{M}_{\mathcal{I}_1} - \mathrm{E}(\boldsymbol{M}_{\mathcal{I}_1} \mid X, \boldsymbol{M}_{\mathcal{A}})\right\}^\top.$$

Also, we have

$$\mathrm{Var}(D_2) \leq \mathrm{E}\{(\omega_{\mathcal{A}} + \omega_{\mathcal{T}})^2(\omega_{\mathcal{A}} - \omega_{\mathcal{T}})^2\} \lesssim \mathrm{E}(\omega_{\mathcal{A}} - \omega_{\mathcal{T}})^2.$$

Further, note that

$$|\omega_{\mathcal{A}} - \omega_{\mathcal{T}}| \lesssim |\boldsymbol{\beta}_{\mathcal{I}_1}^\top\{\mathrm{E}(\boldsymbol{M}_{\mathcal{I}_1} \mid X, \boldsymbol{M}_{\mathcal{A}}) - \mathrm{E}(\boldsymbol{M}_{\mathcal{I}_1} \mid X, \boldsymbol{M}_{\mathcal{T}})\}| \lesssim \|\boldsymbol{\beta}_{\mathcal{I}_1}\|_2 \lesssim \sqrt{s\log(p)/n}.$$

Thus, $\mathrm{Var}(D_2) \lesssim s\log(p)/n = o(1)$.

As a result, we have

$$\sqrt{n}|\widetilde{\rho}_{\mathcal{A}}^2 - \widetilde{\rho}_{\mathcal{T}}^2| \lesssim D_1 + D_2 = o_p(1).$$

This bound holds uniformly for $|\mathcal{A}| \leq s$.

**Analysis of the oracle estimator.** Now, we turn to the asymptotic distribution of $\sqrt{n}(\widetilde{\rho}_{\mathcal{T}}^2 - \rho_{\mathcal{T}}^2)$. Note that $\boldsymbol{\varepsilon} = \boldsymbol{\omega}_{\mathcal{T}}$ and $\boldsymbol{\zeta} = \boldsymbol{\zeta}_{\mathcal{T}}$. By the central limit theorem,

$$\frac{1}{\sqrt{n}}\begin{pmatrix} \boldsymbol{\varepsilon}^\top\boldsymbol{\varepsilon} - V_{Y|MX} \\ \boldsymbol{\eta}^\top\boldsymbol{\eta} - V_{Y|X} \\ \boldsymbol{\zeta}^\top\boldsymbol{\zeta} - V_{Y|M} \\ \boldsymbol{y}^\top\boldsymbol{y} - V_Y \end{pmatrix} \xrightarrow{d} N\left(\boldsymbol{0}, \underbrace{\begin{pmatrix} \mathrm{Var}(\varepsilon^2) & \mathrm{Cov}(\varepsilon^2, \eta^2) & \mathrm{Cov}(\varepsilon^2, \zeta^2) & \mathrm{Cov}(\varepsilon^2, Y^2) \\ \mathrm{Cov}(\varepsilon^2, \eta^2) & \mathrm{Var}(\eta^2) & \mathrm{Cov}(\eta^2, \zeta^2) & \mathrm{Cov}(\eta^2, Y^2) \\ \mathrm{Cov}(\varepsilon^2, \zeta^2) & \mathrm{Cov}(\eta^2, \zeta^2) & \mathrm{Var}(\zeta^2) & \mathrm{Cov}(\zeta^2, Y^2) \\ \mathrm{Cov}(\varepsilon^2, Y^2) & \mathrm{Cov}(\eta^2, Y^2) & \mathrm{Cov}(\zeta^2, Y^2) & \mathrm{Var}(Y^2) \end{pmatrix}}_{=\boldsymbol{A}}\right).$$

Consequently,

$$\sqrt{n}(\widetilde{\rho}_{\mathcal{T}}^2 - \rho_{\mathcal{T}}^2)/\sqrt{\boldsymbol{u}^\top\boldsymbol{A}\boldsymbol{u}} \xrightarrow{d} N(0, 1),$$

where $\boldsymbol{u} = (1/V_Y, -1/V_Y, -1/V_Y, (V_{Y|X} + V_{Y|M} - V_{Y|MX})/V_Y^2)$.

**Asymptotic distribution of $\sqrt{n}(\widehat{R}_{Med}^2 - R_{Med}^2)$.** We characterize $\widehat{\rho}_{\widehat{\mathcal{T}}^{(2)}}^2$ and $\widehat{\rho}_{\widehat{\mathcal{T}}^{(1)}}^2$ as $1 - (\widehat{V}_{Y|X}^{(1)} + \widehat{V}_{Y|M}^{(1)} - \widehat{V}_{Y|MX}^{(1)})/\widehat{V}_Y^{(1)}$ and $1 - (\widehat{V}_{Y|X}^{(2)} + \widehat{V}_{Y|M}^{(2)} - \widehat{V}_{Y|MX}^{(2)})/\widehat{V}_Y^{(2)}$, respectively. Following the above analysis, with $n$ being replaced by $n/2$, we have $\widehat{\rho}_{(1)}^2 = 1 - (\widehat{V}_{Y|X}^{(1)} + \widehat{V}_{Y|M}^{(1)} - \widehat{V}_{Y|MX}^{(1)})/\widehat{V}_Y^{(1)}$ and $\widehat{\rho}_{(2)}^2 = 1 - (\widehat{V}_{Y|X}^{(2)} + \widehat{V}_{Y|M}^{(2)} - \widehat{V}_{Y|MX}^{(2)})/\widehat{V}_Y^{(2)}$, both are asymptotically independent and normal in that

$$\sqrt{n/2}(\widehat{\rho}_{(1)}^2 - R_{Med}^2)/\sqrt{\boldsymbol{u}^\top\boldsymbol{A}\boldsymbol{u}} \xrightarrow{d} N(0, 1),$$
$$\sqrt{n/2}(\widehat{\rho}_{(2)}^2 - R_{Med}^2)/\sqrt{\boldsymbol{u}^\top\boldsymbol{A}\boldsymbol{u}} \xrightarrow{d} N(0, 1).$$

Consequently,

$$\sqrt{n}(\widehat{R}^2_{Med} - R^2_{Med})/\sqrt{\boldsymbol{u}^\top \boldsymbol{A}\boldsymbol{u}} = \sqrt{n/2}((\widehat{\rho}^2_{(1)} + \widehat{\rho}^2_{(2)})/2 - R^2_{Med})/\sqrt{\boldsymbol{u}^\top \boldsymbol{A}\boldsymbol{u}} \xrightarrow{d} N(0,1),$$

which completes the proof.

**Asymptotic distribution of SOS.** The shared over simple effect (SOS) is defined as

$$\text{SOS} = \frac{R^2_{Mediated}}{R^2_{Y,X}} = \frac{V_Y - V_{Y|X} - V_{Y|M} + V_{Y|MX}}{V_Y - V_{Y|X}} = 1 - \frac{V_{Y|M} - V_{Y|MX}}{V_Y - V_{Y|X}}.$$

Let the estimate for SOS be $\widehat{\text{SOS}} = \frac{1}{2}\sum_{k=1}^{2}(1 - (\widehat{V}^{(k)}_{Y|M} - \widehat{V}^{(k)}_{Y|MX})/(\widehat{V}^{(k)}_{Y} - \widehat{V}^{(k)}_{Y|X}))$. Similarly, we can show that

$$\sqrt{n}(\widehat{\text{SOS}}^2 - \text{SOS})/\sqrt{\boldsymbol{v}^\top \boldsymbol{A}\boldsymbol{v}} \xrightarrow{d} N(0,1),$$

where $\boldsymbol{v} = (1/(V_Y - V_{Y|X}), -(V_{Y|M} - V_{Y|MX})/(V_Y - V_{Y|X})^2, -1/(V_Y - V_{Y|X}), (V_{Y|M} - V_{Y|MX})/(V_Y - V_{Y|X})^2)$.

### 2 Web Appendix B: Simulation details under high dimensional settings using the CF-OLS method

In this section, we present the details of our simulation studies conducted in high-dimensional settings using the CF-OLS method. We specifically report the simulation outcomes employing CF-OLS, along with False Discovery Rate (FDR) control mechanisms, for both independent and correlated mediators across two distinct correlation structures. Additionally, we report the exact number of selected mediators of each type. Finally, we describe the simulation results for various scenarios where the parameters $\alpha$ and $\beta$ were drawn from Uniform distributions and different normal distributions. To conclude, we illustrate the asymptotic standard errors and the standard deviation of replications in the accompanying figures.

#### 2.1 Simulation results using the CF-OLS method with the FDR control

Table 1 presents the simulation results for the CF-OLS method applied to high-dimensional settings with independent mediators, post-FDR control, across scenarios (A1)–(A6). The results indicate that the coverage probability remains robust, and the average false positive rate is maintained at a low level. However, there is a slight decrease in the average true positive rate in certain instances after performing the FDR control. This may be attributed to the fact that FDR control aims to minimize the false positive rate as much as possible, which inevitably results in the loss of some true mediators during the selection process.

Similarly, Table 2 and Table 3 display the results of the CF-OLS method when selecting correlated mediators. An acceptable coverage rate has been maintained across all scenarios, and the average false positive rate is close to zero. Additionally, the average true positive rate is promising and improves with increasing sample sizes as anticipated.

#### 2.2 Details of Tables in the main manuscript

In this subsection, the effectiveness of mean-based measures across three simulation scenarios is presented in the main text. We specifically evaluated the bias and mean squared error (MSE) of the mean-based mediation effect measures (product measure, proportion measure, and total effect measure) along with the SOS measure in these simulation contexts.

In the subsequent sections, Table 4, Table 5, and Table 6, we apply proportional scaling to normalize the bias relative to the true value or an established benchmark value, thereby expressing the bias as a percentage or fraction of this reference value. It's important to highlight that in scenarios (A7) Table 5 and Table 6, the actual value of the SOS measure is 0, which means there is no scaled bias for this specific measure. The total effect measure was estimated via the regression between $X$ and $Y$.

Table 7 presents the detailed results from the simulations delineated in Table 1 of the main manuscript, which pertain to the CF-OLS method across scenarios (A1)–(A6).

Table 1: Simulation results using the CF-OLS method for independent mediators after FDR control.

| **Scenario** ($R^2_{Med}$) | **N** | **CP** % | **Width** ($\times 10^{-2}$) | **SE** ($\times 10^{-2}$) | **Bias** ($\times 10^{-2}$) | **SD** ($\times 10^{-2}$) | **MSE** ($\times 10^{-2}$) | **M** | **M-FDR** | **TP** % | **FP** % |
|---|---|---|---|---|---|---|---|---|---|---|---|
| A1 | 750 | **92.5** | 3.673 | 1.874 | 0.670 | 1.951 | 0.042 | 46 | 20 | 92.5 | 2.2 |
| (0.065) | 1500 | **93.0** | 2.604 | 1.328 | 0.627 | 1.322 | 0.021 | 40 | 19 | 92.1 | 1.8 |
| | 3000 | **93.0** | 1.845 | 0.941 | 0.126 | 0.995 | 0.010 | 27 | 17 | 97.4 | 0.8 |
| A2 | 750 | **95.0** | 5.384 | 2.747 | -0.031 | 2.732 | 0.074 | 61 | 60 | 40.0 | 0.1 |
| (0.418) | 1500 | **92.0** | 3.787 | 1.932 | 0.333 | 1.957 | 0.039 | 108 | 104 | 69.6 | 0.3 |
| | 3000 | **94.5** | 2.691 | 1.373 | -0.131 | 1.390 | 0.019 | 145 | 140 | 93.4 | 0.4 |
| A3 | 750 | **95.5** | 5.027 | 2.565 | 0.042 | 2.447 | 0.060 | 61 | 52 | 34.4 | 0.7 |
| (0.064) | 1500 | **96.0** | 3.553 | 1.813 | 0.283 | 1.689 | 0.029 | 110 | 86 | 57.2 | 1.8 |
| | 3000 | **96.5** | 2.504 | 1.277 | 0.156 | 1.147 | 0.013 | 202 | 143 | 88.1 | 5.2 |
| A4 | 750 | **96.0** | 5.445 | 2.778 | 0.029 | 2.769 | 0.076 | 54 | 54 | 35.7 | 0.0 |
| (0.390) | 1500 | **95.0** | 3.845 | 1.962 | -0.255 | 1.957 | 0.039 | 88 | 87 | 58.1 | 0.0 |
| | 3000 | **97.0** | 2.720 | 1.388 | 0.113 | 1.303 | 0.017 | 110 | 108 | 72.3 | 0.1 |
| A5 | 750 | **96.0** | 5.448 | 2.780 | 0.009 | 2.616 | 0.068 | 61 | 54 | 36.3 | 0.5 |
| (0.271) | 1500 | **97.0** | 3.836 | 1.957 | 0.174 | 1.813 | 0.033 | 110 | 91 | 60.8 | 1.4 |
| | 3000 | **97.0** | 2.715 | 1.385 | 0.030 | 1.294 | 0.017 | 201 | 145 | 90.5 | 4.8 |
| A6 | 750 | **96.5** | 5.452 | 2.782 | 0.020 | 2.739 | 0.075 | 61 | 57 | 38.0 | 0.3 |
| (0.377) | 1500 | **93.5** | 3.866 | 1.972 | 0.032 | 2.117 | 0.045 | 106 | 90 | 59.9 | 1.2 |
| | 3000 | **95.5** | 2.737 | 1.396 | -0.040 | 1.388 | 0.019 | 190 | 128 | 69.9 | 6.3 |

$N$ refers to sample size. CP refers to coverage probability based on 200 replications. Width refers to half the width of the 95% confidence interval. SE refers to the average asymptotic standard error. SD refers to the empirical standard deviation of replicated estimations. MSE refers to mean squared error. M refers to the number of selected mediators. M-FDR refers to the number of selected mediators after the FDR control. TP refers to the average true positive rate. FP refers to the average false positive rate. True value of $R^2_{Med}$ is listed within the parentheses.

Table 2: Simulation results using the CF-OLS method for correlated mediators under correlation structure 1 after FDR control.

| **Scenario** ($R^2_{Med}$) | **N** | **CP** % | **Width** ($\times10^{-2}$) | **SE** ($\times10^{-2}$) | **Bias** ($\times10^{-2}$) | **SD** ($\times10^{-2}$) | **MSE** ($\times10^{-2}$) | **M** | **M-FDR** | **TP** % | **FP** % |
|---|---|---|---|---|---|---|---|---|---|---|---|
| A8 | 750 | **92.0** | 5.776 | 2.947 | -0.674 | 2.971 | 0.092 | 5 | 5 | 97.2 | 0.0 |
| (0.128) | 1500 | **93.5** | 3.997 | 2.039 | -0.241 | 2.192 | 0.048 | 5 | 5 | 99.8 | 0.0 |
| | 3000 | **94.5** | 2.830 | 1.444 | -0.060 | 1.484 | 0.022 | 5 | 5 | 100.0 | 0.0 |
| A9 | 750 | **96.0** | 4.075 | 2.079 | -0.092 | 1.957 | 0.038 | 55 | 50 | 82.4 | 0.1 |
| (0.645) | 1500 | **96.0** | 2.879 | 1.469 | -0.146 | 1.445 | 0.021 | 75 | 56 | 85.4 | 0.3 |
| | 3000 | **93.0** | 2.049 | 1.045 | -0.403 | 1.074 | 0.013 | 57 | 53 | 86.7 | 0.1 |
| A10 | 750 | **95.0** | 5.504 | 2.808 | -0.018 | 2.995 | 0.089 | 60 | 26 | 85.9 | 0.6 |
| (0.315) | 1500 | **95.0** | 3.913 | 1.996 | -0.271 | 1.903 | 0.037 | 91 | 33 | 94.4 | 1.0 |
| | 3000 | **92.5** | 2.777 | 1.417 | -0.115 | 1.479 | 0.022 | 82 | 31 | 95.0 | 0.8 |
| A11 | 750 | **92.5** | 2.016 | 1.028 | 0.565 | 1.131 | 0.016 | 42 | 27 | 95.8 | 0.5 |
| (0.015) | 1500 | **94.5** | 1.428 | 0.729 | 0.326 | 0.764 | 0.007 | 40 | 25 | 96.8 | 0.4 |
| | 3000 | **94.5** | 0.996 | 0.508 | 0.190 | 0.500 | 0.003 | 32 | 24 | 98.4 | 0.3 |

$N$ refers to sample size. CP refers to coverage probability based on 200 replications. Width refers to half the width of the 95% confidence interval. SE refers to the average asymptotic standard error. SD refers to the empirical standard deviation of replicated estimations. MSE refers to mean squared error. M refers to the number of selected mediators. M-FDR refers to the number of selected mediators after the FDR control. TP refers to the average true positive rate. FP refers to the average false positive rate. True value of $R^2_{Med}$ is listed within the parentheses.

Table 3: Simulation results using the CF-OLS method for correlated mediators under correlation structure 2 after FDR control.

| **Scenario** ($R^2_{Med}$) | **N** | **CP** % | **Width** ($\times10^{-2}$) | **SE** ($\times10^{-2}$) | **Bias** ($\times10^{-2}$) | **SD** ($\times10^{-2}$) | **MSE** ($\times10^{-2}$) | **M** | **M-FDR** | **TP** % | **FP** % |
|---|---|---|---|---|---|---|---|---|---|---|---|
| A8 | 750 | **92.5** | 5.169 | 2.637 | 0.054 | 2.832 | 0.080 | 5 | 5 | 97.4 | 0.0 |
| (0.128) | 1500 | **94.5** | 3.666 | 1.871 | -0.255 | 1.827 | 0.034 | 5 | 5 | 99.7 | 0.0 |
| | 3000 | **94.5** | 2.600 | 1.327 | -0.250 | 1.299 | 0.017 | 5 | 5 | 100.0 | 0.0 |
| A9 | 750 | **95.0** | 4.220 | 2.153 | -0.204 | 2.107 | 0.045 | 54 | 48 | 78.0 | 0.1 |
| (0.645) | 1500 | **95.5** | 2.992 | 1.527 | -0.053 | 1.498 | 0.022 | 99 | 58 | 78.7 | 0.7 |
| | 3000 | **95.0** | 2.120 | 1.082 | -0.224 | 1.124 | 0.013 | 75 | 50 | 73.3 | 0.4 |
| A10 | 750 | **93.5** | 5.682 | 2.899 | -0.609 | 2.958 | 0.091 | 55 | 24 | 83.2 | 0.5 |
| (0.315) | 1500 | **95.5** | 4.012 | 2.047 | -0.240 | 2.054 | 0.043 | 84 | 27 | 66.9 | 0.9 |
| | 3000 | **92.5** | 2.834 | 1.446 | -0.440 | 1.416 | 0.022 | 61 | 22 | 62.4 | 0.7 |
| A11 | 750 | **95.0** | 1.784 | 0.910 | 0.449 | 0.888 | 0.010 | 36 | 24 | 93.8 | 0.4 |
| (0.015) | 1500 | **95.0** | 1.278 | 0.652 | 0.312 | 0.707 | 0.006 | 23 | 20 | 94.4 | 0.1 |
| | 3000 | **95.0** | 0.908 | 0.463 | 0.217 | 0.441 | 0.002 | 21 | 20 | 95.0 | 0.1 |

$N$ refers to sample size. CP refers to coverage probability based on 200 replications. Width refers to half the width of the 95% confidence interval. SE refers to the average asymptotic standard error. SD refers to the empirical standard deviation of replicated estimations. MSE refers to mean squared error. M refers to the number of selected mediators. M-FDR refers to the number of selected mediators after the FDR control. TP refers to the average true positive rate. FP refers to the average false positive rate. True value of $R^2_{Med}$ is listed within the parentheses.

Table 4: Details of simulation results using the mean-based measures via CF-OLS method for independent mediators in Table 1 in the main manuscript.

| Scenario | $N$ | SOS_bias | SOS_MSE | ab_bias | ab_MSE | prop_bias | prop_MSE | total_bias | total_MSE |
|---|---|---|---|---|---|---|---|---|---|
| **A1** | 750 | $-9.46 \times 10^{-3}$ | $4.06 \times 10^{-4}$ | $2.66 \times 10^{-1}$ | $2.09 \times 10^{0}$ | $4.34 \times 10^{-1}$ | $1.04 \times 10^{1}$ | $-1.87 \times 10^{-3}$ | $1.10 \times 10^{-1}$ |
| | 1500 | $-6.00 \times 10^{-4}$ | $2.14 \times 10^{-4}$ | $2.67 \times 10^{-1}$ | $2.05 \times 10^{0}$ | $4.32 \times 10^{-1}$ | $2.86 \times 10^{0}$ | $2.31 \times 10^{-3}$ | $5.68 \times 10^{-2}$ |
| | 3000 | $-1.67 \times 10^{-3}$ | $1.09 \times 10^{-4}$ | $2.65 \times 10^{-1}$ | $1.99 \times 10^{0}$ | $4.41 \times 10^{-1}$ | $2.76 \times 10^{0}$ | $-4.47 \times 10^{-3}$ | $3.25 \times 10^{-2}$ |
| **A2** | 750 | $1.67 \times 10^{-4}$ | $5.68 \times 10^{-8}$ | $-1.26 \times 10^{-1}$ | $6.74 \times 10^{1}$ | $-2.56 \times 10^{0}$ | $4.55 \times 10^{3}$ | $6.30 \times 10^{-4}$ | $9.06 \times 10^{-1}$ |
| | 1500 | $1.28 \times 10^{-5}$ | $3.89 \times 10^{-10}$ | $-1.10 \times 10^{-1}$ | $6.23 \times 10^{1}$ | $-3.78 \times 10^{-1}$ | $3.09 \times 10^{2}$ | $4.46 \times 10^{-4}$ | $4.41 \times 10^{-1}$ |
| | 3000 | $1.34 \times 10^{-5}$ | $2.09 \times 10^{-10}$ | $6.91 \times 10^{-2}$ | $1.81 \times 10^{1}$ | $7.77 \times 10^{-1}$ | $5.96 \times 10^{-1}$ | $4.18 \times 10^{-5}$ | $2.33 \times 10^{-1}$ |
| **A3** | 750 | $2.30 \times 10^{-3}$ | $1.00 \times 10^{-5}$ | $-5.57 \times 10^{-2}$ | $9.44 \times 10^{1}$ | $5.46 \times 10^{-1}$ | $5.18 \times 10^{0}$ | $2.57 \times 10^{-3}$ | $9.40 \times 10^{0}$ |
| | 1500 | $1.10 \times 10^{-3}$ | $2.37 \times 10^{-6}$ | $-1.14 \times 10^{-1}$ | $1.00 \times 10^{2}$ | $3.08 \times 10^{-1}$ | $1.74 \times 10^{1}$ | $-1.15 \times 10^{-3}$ | $4.76 \times 10^{0}$ |
| | 3000 | $5.30 \times 10^{-4}$ | $5.83 \times 10^{-7}$ | $-1.15 \times 10^{-1}$ | $7.80 \times 10^{1}$ | $1.53 \times 10^{-1}$ | $3.57 \times 10^{1}$ | $1.33 \times 10^{-3}$ | $2.33 \times 10^{0}$ |
| **A4** | 750 | $1.95 \times 10^{-3}$ | $7.53 \times 10^{-6}$ | $-4.88 \times 10^{-1}$ | $9.63 \times 10^{1}$ | $5.41 \times 10^{-1}$ | $1.31 \times 10^{1}$ | $-1.17 \times 10^{-3}$ | $1.03 \times 10^{0}$ |
| | 1500 | $3.69 \times 10^{-4}$ | $3.93 \times 10^{-7}$ | $-3.18 \times 10^{-1}$ | $3.77 \times 10^{1}$ | $4.54 \times 10^{-1}$ | $1.46 \times 10^{1}$ | $-4.28 \times 10^{-4}$ | $4.63 \times 10^{-1}$ |
| | 3000 | $-1.69 \times 10^{-5}$ | $1.55 \times 10^{-9}$ | $-3.44 \times 10^{-1}$ | $4.00 \times 10^{1}$ | $-1.89 \times 10^{-1}$ | $2.64 \times 10^{1}$ | $-1.65 \times 10^{-4}$ | $2.60 \times 10^{-1}$ |
| **A5** | 750 | $4.80 \times 10^{-4}$ | $4.50 \times 10^{-7}$ | $-1.11 \times 10^{-1}$ | $6.72 \times 10^{1}$ | $1.08 \times 10^{-1}$ | $1.19 \times 10^{1}$ | $6.47 \times 10^{-4}$ | $1.66 \times 10^{0}$ |
| | 1500 | $9.82 \times 10^{-5}$ | $2.14 \times 10^{-8}$ | $-1.27 \times 10^{-1}$ | $6.94 \times 10^{1}$ | $3.71 \times 10^{-1}$ | $4.20 \times 10^{0}$ | $6.28 \times 10^{-4}$ | $8.78 \times 10^{-1}$ |
| | 3000 | $2.78 \times 10^{-5}$ | $1.49 \times 10^{-9}$ | $-1.08 \times 10^{-1}$ | $6.16 \times 10^{1}$ | $6.81 \times 10^{-1}$ | $3.24 \times 10^{0}$ | $8.81 \times 10^{-5}$ | $4.26 \times 10^{-1}$ |
| **A6** | 750 | $1.27 \times 10^{-3}$ | $3.46 \times 10^{-6}$ | $1.75 \times 10^{-1}$ | $6.71 \times 10^{1}$ | $1.40 \times 10^{-1}$ | $4.60 \times 10^{1}$ | $-8.05 \times 10^{-4}$ | $1.79 \times 10^{0}$ |
| | 1500 | $4.72 \times 10^{-4}$ | $5.59 \times 10^{-7}$ | $1.70 \times 10^{-1}$ | $6.20 \times 10^{1}$ | $6.58 \times 10^{-1}$ | $7.55 \times 10^{1}$ | $1.55 \times 10^{-3}$ | $9.31 \times 10^{-1}$ |
| | 3000 | $5.77 \times 10^{-5}$ | $9.65 \times 10^{-9}$ | $7.56 \times 10^{-2}$ | $6.47 \times 10^{1}$ | $9.93 \times 10^{-2}$ | $6.19 \times 10^{1}$ | $-1.40 \times 10^{-5}$ | $4.37 \times 10^{-1}$ |

$N$ refers to sample size. **ab** refers to the indirect mediation effect. **prop** refers to the proportion measure. **total** refers to the total effect. **MSE** refers to the Mean Squared Error. **bias** refers to the proportional scaled bias.

Table 5: Details of simulation results using the mean-based measures via CF-OLS method for mediators with correlation structure 1 in Table 2 in the main manuscript.

| Scenario | $N$ | SOS_bias | SOS_MSE | ab_bias | ab_MSE | prop_bias | prop_MSE | total_bias | total_MSE |
|---|---|---|---|---|---|---|---|---|---|
| **A7** | 750 | / | $2.70 \times 10^{-2}$ | $1.03 \times 10^{-2}$ | $8.18 \times 10^{-2}$ | $-4.82 \times 10^{-4}$ | $3.09 \times 10^{1}$ | $7.57 \times 10^{-3}$ | $1.08 \times 10^{-1}$ |
| | 1500 | / | $1.31 \times 10^{-2}$ | $5.05 \times 10^{-3}$ | $3.88 \times 10^{-2}$ | $4.60 \times 10^{-1}$ | $7.40 \times 10^{1}$ | $-2.38 \times 10^{-2}$ | $5.72 \times 10^{-2}$ |
| | 3000 | / | $6.13 \times 10^{-3}$ | $2.32 \times 10^{-3}$ | $1.95 \times 10^{-2}$ | $1.20 \times 10^{-1}$ | $9.24 \times 10^{-1}$ | $3.22 \times 10^{-3}$ | $3.17 \times 10^{-2}$ |
| **A8** | 750 | $-9.55 \times 10^{-3}$ | $4.27 \times 10^{-3}$ | $7.68 \times 10^{-1}$ | $6.12 \times 10^{0}$ | $-1.43 \times 10^{0}$ | $2.16 \times 10^{0}$ | $-1.05 \times 10^{-2}$ | $2.04 \times 10^{-2}$ |
| | 1500 | $1.19 \times 10^{-2}$ | $2.92 \times 10^{-3}$ | $7.75 \times 10^{-1}$ | $6.15 \times 10^{0}$ | $-1.43 \times 10^{0}$ | $2.17 \times 10^{0}$ | $8.34 \times 10^{-3}$ | $1.09 \times 10^{-2}$ |
| | 3000 | $1.91 \times 10^{-3}$ | $1.22 \times 10^{-3}$ | $7.71 \times 10^{-1}$ | $6.12 \times 10^{0}$ | $-1.43 \times 10^{0}$ | $2.16 \times 10^{0}$ | $1.55 \times 10^{-3}$ | $5.14 \times 10^{-3}$ |
| **A9** | 750 | $-4.92 \times 10^{-4}$ | $5.05 \times 10^{-7}$ | $-1.19 \times 10^{1}$ | $1.41 \times 10^{2}$ | $1.47 \times 10^{0}$ | $3.64 \times 10^{0}$ | $3.17 \times 10^{-3}$ | $1.11 \times 10^{-1}$ |
| | 1500 | $-2.61 \times 10^{-4}$ | $2.32 \times 10^{-7}$ | $-1.19 \times 10^{1}$ | $1.42 \times 10^{2}$ | $1.39 \times 10^{0}$ | $2.14 \times 10^{0}$ | $-2.46 \times 10^{-2}$ | $5.94 \times 10^{-2}$ |
| | 3000 | $-2.15 \times 10^{-5}$ | $7.78 \times 10^{-8}$ | $-1.19 \times 10^{1}$ | $1.416 \times 10^{2}$ | $1.27 \times 10^{0}$ | $1.65 \times 10^{0}$ | $1.39 \times 10^{-3}$ | $3.33 \times 10^{-2}$ |
| **A10** | 750 | $2.06 \times 10^{-4}$ | $4.67 \times 10^{-6}$ | $-1.22 \times 10^{1}$ | $1.53 \times 10^{2}$ | $1.05 \times 10^{0}$ | $4.66 \times 10^{1}$ | $-1.42 \times 10^{-2}$ | $5.12 \times 10^{-1}$ |
| | 1500 | $-4.89 \times 10^{-4}$ | $4.39 \times 10^{-7}$ | $-1.18 \times 10^{1}$ | $1.42 \times 10^{2}$ | $1.60 \times 10^{0}$ | $5.17 \times 10^{1}$ | $-5.05 \times 10^{-2}$ | $2.07 \times 10^{-1}$ |
| | 3000 | $-2.62 \times 10^{-4}$ | $1.61 \times 10^{-7}$ | $-1.19 \times 10^{1}$ | $1.41 \times 10^{2}$ | $1.17 \times 10^{0}$ | $2.68 \times 10^{1}$ | $-1.46 \times 10^{-2}$ | $1.31 \times 10^{-1}$ |
| **A11** | 750 | $-7.77 \times 10^{-2}$ | $3.95 \times 10^{-2}$ | $-4.45 \times 10^{0}$ | $2.14 \times 10^{1}$ | $3.96 \times 10^{0}$ | $1.58 \times 10^{1}$ | $-1.57 \times 10^{-3}$ | $1.29 \times 10^{-1}$ |
| | 1500 | $-6.84 \times 10^{-2}$ | $1.44 \times 10^{-2}$ | $-4.24 \times 10^{0}$ | $1.81 \times 10^{1}$ | $4.04 \times 10^{0}$ | $1.63 \times 10^{1}$ | $5.03 \times 10^{-3}$ | $7.11 \times 10^{-2}$ |
| | 3000 | $-8.08 \times 10^{-2}$ | $7.77 \times 10^{-3}$ | $-4.23 \times 10^{0}$ | $1.80 \times 10^{1}$ | $4.04 \times 10^{0}$ | $1.64 \times 10^{1}$ | $1.44 \times 10^{-3}$ | $3.14 \times 10^{-2}$ |
| **A12** | 750 | $-1.36 \times 10^{-1}$ | $5.20 \times 10^{-2}$ | $-6.35 \times 10^{0}$ | $9.46 \times 10^{1}$ | $3.47 \times 10^{0}$ | $1.61 \times 10^{1}$ | $-3.33 \times 10^{-3}$ | $6.56 \times 10^{-1}$ |
| | 1500 | $-1.48 \times 10^{-1}$ | $3.02 \times 10^{-2}$ | $-4.26 \times 10^{0}$ | $1.86 \times 10^{1}$ | $4.04 \times 10^{0}$ | $1.64 \times 10^{1}$ | $1.36 \times 10^{-2}$ | $2.93 \times 10^{-1}$ |
| | 3000 | $-1.03 \times 10^{-1}$ | $1.85 \times 10^{-2}$ | $-4.27 \times 10^{0}$ | $1.85 \times 10^{1}$ | $4.04 \times 10^{0}$ | $1.63 \times 10^{1}$ | $-1.07 \times 10^{-2}$ | $1.60 \times 10^{-1}$ |

$N$ refers to sample size. **ab** refers to the indirect mediation effect. **prop** refers to the proportion measure. **total** refers to the total effect. **MSE** refers to the Mean Squared Error. **bias** refers to the proportional scaled bias.

Table 6: Details of simulation results using the mean-based measures via CF-OLS method for mediators with correlation structure 2 in Table 2 in the main manuscript.

| Scenario | $N$ | SOS_bias | SOS_MSE | ab_bias | ab_MSE | prop_bias | prop_MSE | total_bias | total_MSE |
|---|---|---|---|---|---|---|---|---|---|
| **A7** | 750 | / | $3.08 \times 10^{-2}$ | $-1.76 \times 10^{-2}$ | $2.59 \times 10^{-2}$ | $1.61 \times 10^{-1}$ | $2.61 \times 10^{0}$ | $6.98 \times 10^{-3}$ | $1.22 \times 10^{-1}$ |
| | 1500 | / | $1.46 \times 10^{-2}$ | $-4.92 \times 10^{-3}$ | $9.79 \times 10^{-3}$ | $8.22 \times 10^{-2}$ | $1.30 \times 10^{-1}$ | $-8.85 \times 10^{-3}$ | $6.45 \times 10^{-2}$ |
| | 3000 | / | $5.46 \times 10^{-3}$ | $-3.64 \times 10^{-3}$ | $5.63 \times 10^{-3}$ | $4.06 \times 10^{-2}$ | $3.96 \times 10^{-2}$ | $1.08 \times 10^{-2}$ | $2.90 \times 10^{-2}$ |
| **A8** | 750 | $1.07 \times 10^{-2}$ | $5.38 \times 10^{-3}$ | $7.70 \times 10^{-1}$ | $6.09 \times 10^{0}$ | $-1.43 \times 10^{0}$ | $2.16 \times 10^{0}$ | $1.32 \times 10^{-2}$ | $2.75 \times 10^{-2}$ |
| | 1500 | $1.47 \times 10^{-2}$ | $2.26 \times 10^{-3}$ | $7.74 \times 10^{-1}$ | $6.11 \times 10^{0}$ | $-1.43 \times 10^{0}$ | $2.17 \times 10^{0}$ | $1.21 \times 10^{-2}$ | $1.28 \times 10^{-2}$ |
| | 3000 | $1.27 \times 10^{-2}$ | $1.30 \times 10^{-3}$ | $7.74 \times 10^{-1}$ | $6.15 \times 10^{0}$ | $-1.43 \times 10^{0}$ | $2.16 \times 10^{0}$ | $3.87 \times 10^{-3}$ | $6.32 \times 10^{-3}$ |
| **A9** | 750 | $-5.78 \times 10^{-4}$ | $6.64 \times 10^{-7}$ | $-1.19 \times 10^{1}$ | $1.42 \times 10^{2}$ | $1.50 \times 10^{0}$ | $5.18 \times 10^{0}$ | $1.13 \times 10^{-2}$ | $1.25 \times 10^{-1}$ |
| | 1500 | $-1.85 \times 10^{-4}$ | $2.29 \times 10^{-7}$ | $-1.19 \times 10^{1}$ | $1.43 \times 10^{2}$ | $1.42 \times 10^{0}$ | $2.17 \times 10^{0}$ | $-1.08 \times 10^{-2}$ | $6.50 \times 10^{-2}$ |
| | 3000 | $-3.78 \times 10^{-6}$ | $9.83 \times 10^{-8}$ | $-1.19 \times 10^{1}$ | $1.42 \times 10^{2}$ | $1.30 \times 10^{0}$ | $1.73 \times 10^{0}$ | $1.18 \times 10^{-2}$ | $2.96 \times 10^{-2}$ |
| **A10** | 750 | $3.11 \times 10^{-3}$ | $2.71 \times 10^{-5}$ | $-1.22 \times 10^{1}$ | $1.63 \times 10^{2}$ | $4.06 \times 10^{-1}$ | $3.24 \times 10^{2}$ | $-1.57 \times 10^{-2}$ | $5.01 \times 10^{-1}$ |
| | 1500 | $1.65 \times 10^{-4}$ | $9.28 \times 10^{-7}$ | $-1.17 \times 10^{1}$ | $1.43 \times 10^{2}$ | $9.26 \times 10^{-1}$ | $8.56 \times 10^{1}$ | $2.46 \times 10^{-2}$ | $2.42 \times 10^{-1}$ |
| | 3000 | $4.66 \times 10^{-4}$ | $2.71 \times 10^{-5}$ | $-1.19 \times 10^{1}$ | $1.49 \times 10^{2}$ | $1.17 \times 10^{0}$ | $3.33 \times 10^{0}$ | $1.11 \times 10^{-4}$ | $1.25 \times 10^{-1}$ |
| **A11** | 750 | $-3.72 \times 10^{-2}$ | $8.96 \times 10^{-3}$ | $-4.54 \times 10^{0}$ | $2.39 \times 10^{1}$ | $3.99 \times 10^{0}$ | $1.59 \times 10^{1}$ | $-3.33 \times 10^{-2}$ | $1.44 \times 10^{-1}$ |
| | 1500 | $-3.92 \times 10^{-2}$ | $6.38 \times 10^{-3}$ | $-4.24 \times 10^{0}$ | $1.81 \times 10^{1}$ | $4.04 \times 10^{0}$ | $1.63 \times 10^{1}$ | $-1.45 \times 10^{-2}$ | $8.16 \times 10^{-2}$ |
| | 3000 | $-4.84 \times 10^{-2}$ | $3.63 \times 10^{-3}$ | $-4.26 \times 10^{0}$ | $1.83 \times 10^{1}$ | $4.03 \times 10^{0}$ | $1.63 \times 10^{1}$ | $1.06 \times 10^{-3}$ | $3.74 \times 10^{-2}$ |
| **A12** | 750 | $4.66 \times 10^{-2}$ | $6.48 \times 10^{-3}$ | $-6.66 \times 10^{0}$ | $9.67 \times 10^{1}$ | $3.61 \times 10^{0}$ | $3.75 \times 10^{1}$ | $4.46 \times 10^{-2}$ | $5.51 \times 10^{-1}$ |
| | 1500 | $8.53 \times 10^{-3}$ | $2.65 \times 10^{-4}$ | $-5.12 \times 10^{0}$ | $4.29 \times 10^{1}$ | $3.16 \times 10^{0}$ | $8.77 \times 10^{1}$ | $1.81 \times 10^{-2}$ | $2.56 \times 10^{-1}$ |
| | 3000 | $1.16 \times 10^{-2}$ | $3.40 \times 10^{-4}$ | $-5.20 \times 10^{0}$ | $4.15 \times 10^{1}$ | $4.12 \times 10^{0}$ | $7.65 \times 10^{1}$ | $-2.28 \times 10^{-2}$ | $1.35 \times 10^{-1}$ |

$N$ refers to sample size. **ab** refers to the indirect mediation effect. **prop** refers to the proportion measure. **total** refers to the total effect. **MSE** refers to the Mean Squared Error. **bias** refers to the proportional scaled bias.

In this section, we provide the number of the correctly identified true mediators as well as various types of non-mediators as determined by the iSIS-MCP procedure. Further evaluations with increased sample sizes of 500, 1000, and 2000 have been included below, which corroborate the conclusions drawn in the main text of the manuscript.

##### 2.3 Performance of the CF-OLS method in additional scenarios

As discussed in the simulation settings section of the main manuscript, additional simulation settings were considered in the same manner as described above, with the exception of the distributions followed by $\boldsymbol{\alpha}$ and $\boldsymbol{\beta}$. Scenarios (B1)–(B6) illustrate the results of the CF-OLS method when applied to the symmetric uniform distribution $Unif(-2, 2)$, while scenarios (C1)–(C6) pertain to the standard normal distribution $N(0, 1^2)$. The simulation details for the CF-OLS method under scenarios (B1)–(B6) and (C1)–(C6) are displayed in Table 8 and Table 9, respectively.

We observed that the coverage probability in most of the scenarios from (B1)–(B6) to (C1)–(C6) was consistently promising. The asymptotic standard errors were found to be close to the empirical standard deviation of the estimations in all scenarios, regardless of the sample size. A similar pattern was noted for the Mean Squared Error (MSE), which was higher for limited sample sizes and decreased as the sample size was increased. Meanwhile, the iSIS-MCP method performed well and consistently under these various settings, providing a high average true positive rate and a low average false positive rate, all within a similar computational time frame.

Overall, the performance of our proposed method is adequate for estimation and inference purposes, with high computational efficiency.

##### 2.4 Asymptotic standard error and empirical standard deviation under different settings

Here we present figures depicting the asymptotic standard error and empirical standard deviation of estimations under two additional settings, serving as complements in Figure 1 and Figure 2. We found that our proposed CF-OLS method demonstrates robustness across diverse settings, with the asymptotic standard errors closely aligning with the empirical standard deviations in all scenarios.

#### 3 Web Appendix C: Performance of various iSIS settings

In this section, we perform simulations across various settings using the iSIS variable selection procedure to evaluate its inferential performance. Our CF-OLS method was applied using the same settings as in Table 1 of the main manuscript, specifically, the normal distribution $N(0, 1.5^2)$.

Firstly, in Table 10, we substituted the MCP penalty with the Lasso penalty to assess the performance of iSIS-Lasso. For this approach, we employed the Akaike

Table 7: Details of simulation results using the CF-OLS method for independent mediators in Table 1 in the main manuscript.

| **Scen** ($R^2_{Med}$) | **N** | **CP** % | **Width** ($\times 10^{-2}$) | **SE** ($10^{-2}$) | **Bias** ($10^{-2}$) | **SD** ($10^{-2}$) | **MSE** ($10^{-4}$) | $\boldsymbol{M}_{\mathcal{T}}$ | $\boldsymbol{M}_{\mathcal{I}_1}$ | $\boldsymbol{M}_{\mathcal{I}_2}$ | $\boldsymbol{M}_{\mathcal{I}_3}$ | **TP** | **FP** | **Time** |
|---|---|---|---|---|---|---|---|---|---|---|---|---|---|---|
| **A1** | 500 | 93.0 | 4.49 | 2.29 | 1.07 | 2.37 | 6.72 | 14.0 | 0.0 | 0.0 | 22.1 | 0.94 | 0.01 | 0.08 (0.00) |
| (0.065) | 750 | 92.0 | 3.66 | 1.87 | 0.74 | 1.94 | 4.29 | 14.2 | 0.0 | 0.0 | 31.7 | 0.94 | 0.02 | 0.12 (0.00) |
| | 1000 | 93.0 | 3.17 | 1.62 | 0.74 | 1.73 | 3.51 | 14.0 | 0.0 | 0.0 | 34.0 | 0.94 | 0.02 | 1.01 (0.03) |
| | 1500 | 93.5 | 2.60 | 1.33 | 0.66 | 1.32 | 2.16 | 13.9 | 0.0 | 0.0 | 26.1 | 0.93 | 0.02 | 3.44 (0.04) |
| | 2000 | 90.5 | 2.24 | 1.14 | 0.18 | 1.31 | 1.75 | 14.3 | 0.0 | 0.0 | 26.3 | 0.95 | 0.02 | 2.88 (0.04) |
| | 3000 | 93.5 | 1.84 | 0.94 | 0.13 | 0.99 | 1.00 | 14.5 | 0.0 | 0.0 | 12.0 | 0.97 | 0.01 | 4.80 (0.07) |
| **A2** | 500 | 94.0 | 6.57 | 3.35 | -0.07 | 3.44 | 11.80 | 42.6 | 0.0 | 0.0 | 0.8 | 0.28 | 0.00 | 2.01 (0.03) |
| (0.418) | 750 | 94.5 | 5.38 | 2.75 | -0.03 | 2.74 | 7.45 | 60.4 | 0.0 | 0.0 | 0.8 | 0.40 | 0.00 | 1.98 (0.04) |
| | 1000 | 95.5 | 4.64 | 2.37 | 0.09 | 2.40 | 5.73 | 76.9 | 0.0 | 0.0 | 1.2 | 0.51 | 0.00 | 1.56 (0.03) |
| | 1500 | 92.0 | 3.79 | 1.93 | 0.33 | 1.96 | 3.92 | 104.1 | 0.0 | 0.0 | 4.1 | 0.69 | 0.00 | 5.30 (0.11) |
| | 2000 | 96.5 | 3.29 | 1.68 | 0.11 | 1.58 | 2.48 | 126.6 | 0.0 | 0.0 | 6.0 | 0.84 | 0.00 | 6.83 (0.12) |
| | 3000 | 94.5 | 2.69 | 1.37 | -0.13 | 1.39 | 1.94 | 141.5 | 0.0 | 0.0 | 3.5 | 0.94 | 0.00 | 6.78 (0.04) |
| **A3** | 500 | 96.0 | 4.27 | 2.18 | 0.23 | 2.11 | 4.47 | 34.1 | 9.7 | 0.0 | 0.0 | 0.23 | 0.01 | 0.66 (0.03) |
| (0.064) | 750 | 93.5 | 3.49 | 1.78 | 0.27 | 1.79 | 3.26 | 46.5 | 15.0 | 0.0 | 0.0 | 0.31 | 0.01 | 2.13 (0.04) |
| | 1000 | 94.5 | 2.97 | 1.52 | 0.07 | 1.61 | 2.58 | 56.8 | 21.4 | 0.0 | 0.0 | 0.38 | 0.02 | 1.45 (0.03) |
| | 1500 | 95.0 | 2.43 | 1.24 | 0.20 | 1.26 | 1.62 | 75.8 | 34.9 | 0.0 | 0.0 | 0.51 | 0.03 | 5.10 (0.05) |
| | 2000 | 94.5 | 2.09 | 1.07 | 0.11 | 1.07 | 1.15 | 91.9 | 49.5 | 0.0 | 0.0 | 0.61 | 0.04 | 3.16 (0.05) |
| | 3000 | 95.0 | 1.71 | 0.87 | 0.17 | 0.82 | 0.69 | 114.3 | 88.0 | 0.0 | 0.0 | 0.76 | 0.07 | 8.62 (0.10) |
| **A4** | 500 | 94.5 | 6.69 | 3.41 | 0.08 | 3.50 | 12.19 | 11.5 | 0.0 | 31.4 | 0.0 | 0.08 | 0.02 | 0.68 (0.01) |
| (0.390) | 750 | 96.0 | 5.44 | 2.78 | 0.03 | 2.77 | 7.63 | 19.4 | 0.0 | 34.3 | 0.0 | 0.13 | 0.03 | 1.47 (0.03) |
| | 1000 | 95.0 | 4.71 | 2.40 | -0.12 | 2.46 | 6.03 | 31.1 | 0.0 | 33.8 | 0.0 | 0.21 | 0.03 | 2.13 (0.04) |
| | 1500 | 95.0 | 3.85 | 1.96 | -0.26 | 1.96 | 3.87 | 57.9 | 0.0 | 29.7 | 0.0 | 0.39 | 0.02 | 4.95 (0.08) |
| | 2000 | 92.5 | 3.32 | 1.70 | 0.10 | 1.78 | 3.16 | 80.9 | 0.0 | 17.8 | 0.0 | 0.54 | 0.01 | 2.91 (0.05) |
| | 3000 | 97.0 | 2.72 | 1.39 | 0.11 | 1.30 | 1.70 | 108.6 | 0.0 | 1.8 | 0.0 | 0.72 | 0.00 | 6.78 (0.12) |
| **A5** | 500 | 94.0 | 6.67 | 3.41 | 0.23 | 3.45 | 11.87 | 38.6 | 2.7 | 0.0 | 2.2 | 0.26 | 0.00 | 1.10 (0.03) |
| (0.271) | 750 | 96.0 | 5.44 | 2.78 | 0.02 | 2.61 | 6.80 | 52.9 | 5.7 | 0.0 | 2.5 | 0.35 | 0.01 | 1.39 (0.02) |
| | 1000 | 94.5 | 4.71 | 2.40 | 0.19 | 2.30 | 5.31 | 65.4 | 9.6 | 0.0 | 2.8 | 0.44 | 0.01 | 3.04 (0.08) |
| | 1500 | 97.0 | 3.83 | 1.96 | 0.18 | 1.81 | 3.31 | 86.7 | 20.2 | 0.0 | 3.5 | 0.58 | 0.02 | 3.10 (0.08) |
| | 2000 | 96.0 | 3.31 | 1.69 | 0.08 | 1.61 | 2.58 | 102.3 | 33.5 | 0.0 | 5.2 | 0.68 | 0.03 | 5.73 (0.08) |
| | 3000 | 97.0 | 2.71 | 1.38 | 0.05 | 1.29 | 1.66 | 131.9 | 61.3 | 0.0 | 7.4 | 0.88 | 0.05 | 8.88 (0.12) |
| **A6** | 500 | 93.5 | 6.66 | 3.40 | -0.07 | 3.55 | 12.53 | 25.8 | 1.6 | 15.8 | 0.4 | 0.17 | 0.01 | 0.79 (0.01) |
| (0.377) | 750 | 96.5 | 5.45 | 2.78 | 0.04 | 2.74 | 7.47 | 35.7 | 4.3 | 20.2 | 0.8 | 0.24 | 0.02 | 2.42 (0.04) |
| | 1000 | 95.0 | 4.74 | 2.42 | -0.05 | 2.50 | 6.24 | 44.6 | 7.6 | 23.7 | 1.0 | 0.30 | 0.02 | 1.93 (0.03) |
| | 1500 | 92.5 | 3.86 | 1.97 | 0.05 | 2.11 | 4.45 | 60.1 | 17.7 | 26.2 | 1.9 | 0.40 | 0.03 | 4.14 (0.10) |
| | 2000 | 95.0 | 3.35 | 1.71 | -0.01 | 1.71 | 2.90 | 72.9 | 32.8 | 25.1 | 3.3 | 0.49 | 0.05 | 4.78 (0.12) |
| | 3000 | 95.5 | 2.74 | 1.40 | -0.02 | 1.39 | 1.92 | 93.3 | 62.0 | 21.1 | 13.6 | 0.62 | 0.07 | 8.34 (0.12) |

$N$ refers to sample size. CP refers to coverage probability based on 200 replications. Width refers to half the width of the 95% confidence interval. SE refers to the average asymptotic standard error. SD refers to the empirical standard deviation of replicated estimations. MSE refers to mean squared error. $(\boldsymbol{M}_{\mathcal{T}}, \boldsymbol{M}_{\mathcal{I}_1}, \boldsymbol{M}_{\mathcal{I}_2}, \boldsymbol{M}_{\mathcal{I}_3})$ refer to the number of selected true mediators, three types of non-mediators respectively. TP refers to the average true positive rate. FP refers to the average false positive rate. True value of $R^2_{Med}$ is listed within the parentheses. Time refers to the average computational time in minutes and its standard error is listed within the parentheses. The computational times for CF-OLS is observed using a single core.

Table 8: Simulation results under the uniform distribution $Unif(-2, 2)$ setting.

| **Scen** ($R^2_{Med}$) | **N** | **CP** % | **Width** ($\times 10^{-2}$) | **SE** ($10^{-2}$) | **Bias** ($10^{-2}$) | **SD** ($10^{-2}$) | **MSE** ($10^{-4}$) | $\boldsymbol{M_{\mathcal{T}}}$ | $\boldsymbol{M_{\mathcal{I}_1}}$ | $\boldsymbol{M_{\mathcal{I}_2}}$ | $\boldsymbol{M_{\mathcal{I}_3}}$ | **TP** | **FP** | **Time** |
|---|---|---|---|---|---|---|---|---|---|---|---|---|---|---|
| **B1** | 500 | 92.5 | 5.18 | 2.64 | 0.35 | 2.67 | 7.21 | 12.1 | 0.0 | 0.0 | 24.7 | 0.80 | 0.02 | 0.07 (0.00) |
| (0.620) | 750 | 94.0 | 4.24 | 2.17 | 0.16 | 2.35 | 5.54 | 12.1 | 0.0 | 0.0 | 24.0 | 0.81 | 0.02 | 0.42 (0.01) |
| | 1000 | 93.5 | 3.70 | 1.89 | 0.12 | 1.99 | 3.97 | 12.1 | 0.0 | 0.0 | 9.0 | 0.80 | 0.01 | 0.68 (0.01) |
| | 1500 | 95.0 | 3.02 | 1.54 | 0.12 | 1.53 | 2.36 | 12.2 | 0.0 | 0.0 | 1.9 | 0.81 | 0.00 | 1.29 (0.02) |
| | 2000 | 94.5 | 2.63 | 1.34 | 0.05 | 1.33 | 1.77 | 12.4 | 0.0 | 0.0 | 0.9 | 0.83 | 0.00 | 2.52 (0.05) |
| | 3000 | 93.5 | 2.14 | 1.09 | 0.05 | 1.15 | 1.31 | 13.0 | 0.0 | 0.0 | 0.9 | 0.86 | 0.00 | 3.63 (0.12) |
| **B2** | 500 | 94.0 | 5.14 | 2.62 | 0.18 | 2.65 | 7.04 | 42.1 | 0.0 | 0.0 | 1.6 | 0.28 | 0.00 | 1.60 (0.02) |
| (0.627) | 750 | 95.5 | 4.21 | 2.15 | -0.03 | 2.25 | 5.04 | 60.2 | 0.0 | 0.0 | 1.5 | 0.40 | 0.00 | 1.68 (0.04) |
| | 1000 | 92.5 | 3.63 | 1.85 | 0.12 | 1.97 | 3.89 | 77.1 | 0.0 | 0.0 | 1.5 | 0.51 | 0.00 | 1.52 (0.06) |
| | 1500 | 97.5 | 2.99 | 1.53 | -0.04 | 1.43 | 2.04 | 108.8 | 0.0 | 0.0 | 2.8 | 0.73 | 0.00 | 2.45 (0.07) |
| | 2000 | 95.0 | 2.58 | 1.32 | -0.12 | 1.28 | 1.64 | 134.4 | 0.0 | 0.0 | 4.1 | 0.90 | 0.00 | 3.41 (0.08) |
| | 3000 | 96.0 | 2.11 | 1.08 | 0.08 | 0.99 | 0.99 | 144.1 | 0.0 | 0.0 | 2.2 | 0.96 | 0.00 | 4.64 (0.11) |
| **B3** | 500 | 92.5 | 5.92 | 3.02 | 0.30 | 3.15 | 9.98 | 33.5 | 10.2 | 0.0 | 0.0 | 0.22 | 0.01 | 0.93 (0.02) |
| (0.159) | 750 | 93.5 | 4.79 | 2.44 | -0.22 | 2.42 | 5.86 | 44.9 | 16.4 | 0.0 | 0.0 | 0.30 | 0.01 | 0.94 (0.03) |
| | 1000 | 96.0 | 4.18 | 2.13 | 0.15 | 2.20 | 4.84 | 54.7 | 23.3 | 0.0 | 0.0 | 0.36 | 0.02 | 0.72 (0.01) |
| | 1500 | 97.0 | 3.40 | 1.73 | 0.06 | 1.57 | 2.47 | 73.0 | 37.6 | 0.0 | 0.0 | 0.49 | 0.03 | 2.34 (0.04) |
| | 2000 | 95.5 | 2.95 | 1.50 | 0.02 | 1.49 | 2.21 | 90.3 | 51.0 | 0.0 | 0.0 | 0.60 | 0.04 | 3.70 (0.05) |
| | 3000 | 98.0 | 2.40 | 1.23 | 0.15 | 1.22 | 1.51 | 120.3 | 81.5 | 0.0 | 0.0 | 0.80 | 0.06 | 4.97 (0.10) |
| **B4** | 500 | 94.5 | 6.61 | 3.37 | -0.15 | 3.28 | 10.70 | 13.3 | 0.0 | 30.4 | 0.0 | 0.09 | 0.02 | 0.84 (0.01) |
| (0.412) | 750 | 94.5 | 5.42 | 2.77 | -0.15 | 2.82 | 7.93 | 21.9 | 0.0 | 39.1 | 0.0 | 0.15 | 0.03 | 1.13 (0.02) |
| | 1000 | 96.5 | 4.69 | 2.39 | 0.04 | 2.28 | 5.18 | 31.1 | 0.0 | 46.1 | 0.0 | 0.21 | 0.03 | 0.86 (0.02) |
| | 1500 | 93.5 | 3.81 | 1.95 | 0.05 | 1.94 | 3.76 | 52.5 | 0.0 | 53.2 | 0.0 | 0.35 | 0.04 | 1.83 (0.07) |
| | 2000 | 93.0 | 3.30 | 1.68 | -0.07 | 1.70 | 2.88 | 75.9 | 0.0 | 55.3 | 0.0 | 0.51 | 0.04 | 3.57 (0.04) |
| | 3000 | 94.5 | 2.69 | 1.37 | 0.00 | 1.37 | 1.86 | 112.1 | 0.0 | 38.1 | 0.0 | 0.75 | 0.03 | 5.37 (0.06) |
| **B5** | 500 | 94.5 | 6.35 | 3.24 | 0.20 | 3.22 | 10.38 | 36.9 | 3.0 | 0.0 | 3.5 | 0.25 | 0.00 | 0.90 (0.02) |
| (0.467) | 750 | 93.0 | 5.21 | 2.66 | 0.04 | 2.67 | 7.12 | 50.5 | 6.4 | 0.0 | 4.3 | 0.34 | 0.01 | 0.74 (0.02) |
| | 1000 | 93.0 | 4.51 | 2.30 | 0.12 | 2.44 | 5.92 | 63.2 | 10.7 | 0.0 | 4.2 | 0.42 | 0.01 | 1.41 (0.07) |
| | 1500 | 96.0 | 3.69 | 1.89 | -0.05 | 1.85 | 3.40 | 85.5 | 21.2 | 0.0 | 4.2 | 0.57 | 0.02 | 2.76 (0.04) |
| | 2000 | 96.5 | 3.18 | 1.62 | -0.07 | 1.65 | 2.70 | 104.8 | 33.6 | 0.0 | 3.6 | 0.70 | 0.03 | 3.21 (0.07) |
| | 3000 | 98.5 | 2.60 | 1.32 | 0.18 | 1.24 | 1.56 | 136.4 | 63.4 | 0.0 | 3.6 | 0.91 | 0.05 | 4.05 (0.05) |
| **B6** | 500 | 93.5 | 6.73 | 3.43 | 0 | 3.69 | 13.52 | 24.77 | 1.78 | 15.11 | 1.73 | 0.17 | 0.01 | 1.08 (0.02) |
| (0.353) | 750 | 95 | 5.48 | 2.8 | 0.19 | 2.73 | 7.46 | 34.08 | 4.33 | 20.25 | 2.18 | 0.23 | 0.02 | 1.36 (0.02) |
| | 1000 | 95.5 | 4.77 | 2.43 | -0.19 | 2.45 | 6.02 | 42.84 | 7.62 | 24.48 | 2.48 | 0.29 | 0.03 | 2.15 (0.05) |
| | 1500 | 96 | 3.89 | 1.98 | 0.10 | 2.04 | 4.14 | 58.09 | 16.05 | 32.61 | 2.35 | 0.39 | 0.04 | 2.68 (0.07) |
| | 2000 | 93 | 3.37 | 1.72 | -0.16 | 1.88 | 3.53 | 71.62 | 25.96 | 39.01 | 2.34 | 0.48 | 0.05 | 4.79 (0.05) |
| | 3000 | 93 | 2.75 | 1.4 | 0.13 | 1.5 | 2.25 | 97.75 | 49.12 | 46.32 | 2.4 | 0.65 | 0.07 | 7.12 (0.09) |

$N$ refers to sample size. CP refers to coverage probability based on 200 replications. Width refers to half the width of the 95% confidence interval. SE refers to the average asymptotic standard error. SD refers to the empirical standard deviation of replicated estimations. MSE refers to mean squared error. $(\boldsymbol{M_{\mathcal{T}}}, \boldsymbol{M_{\mathcal{I}_1}}, \boldsymbol{M_{\mathcal{I}_2}}, \boldsymbol{M_{\mathcal{I}_3}})$ refer to the number of selected true mediators, three types of non-mediators respectively. TP refers to the average true positive rate. FP refers to the average false positive rate. True value of $R^2_{Med}$ is listed within the parentheses. Time refers to the average computational time in minutes and its standard error is listed within the parentheses. The computational times for CF-OLS is observed using a single core.

Table 9: Simulation results under the normal distribution $N(0, 1^2)$ setting.

| **Scen** ($R^2_{Med}$) | **N** | **CP** % | **Width** ($\times 10^{-2}$) | **SE** ($10^{-2}$) | **Bias** ($10^{-2}$) | **SD** ($10^{-2}$) | **MSE** ($10^{-4}$) | $M_{\mathcal{T}}$ | $M_{\mathcal{I}_1}$ | $M_{\mathcal{I}_2}$ | $M_{\mathcal{I}_3}$ | **TP** | **FP** | **Time** |
|---|---|---|---|---|---|---|---|---|---|---|---|---|---|---|
| **C1** | 500 | 91.5 | 5.28 | 2.69 | -0.50 | 3.03 | 9.40 | 10.3 | 0.0 | 0.0 | 11.2 | 0.69 | 0.01 | 1.46 (0.03) |
| (0.611) | 750 | 95.0 | 4.32 | 2.20 | -0.30 | 2.17 | 4.79 | 10.3 | 0.0 | 0.0 | 2.3 | 0.68 | 0.00 | 1.64 (0.04) |
| | 1000 | 95.5 | 3.72 | 1.90 | -0.20 | 1.87 | 3.54 | 10.3 | 0.0 | 0.0 | 0.5 | 0.68 | 0.00 | 2.01 (0.03) |
| | 1500 | 92.0 | 3.05 | 1.56 | -0.41 | 1.73 | 3.15 | 10.5 | 0.0 | 0.0 | 0.4 | 0.70 | 0.00 | 2.00 (0.07) |
| | 2000 | 94.0 | 2.64 | 1.35 | -0.43 | 1.50 | 2.42 | 10.7 | 0.0 | 0.0 | 0.4 | 0.72 | 0.00 | 3.55 (0.10) |
| | 3000 | 92.0 | 2.16 | 1.10 | -0.43 | 1.10 | 1.38 | 11.0 | 0.0 | 0.0 | 0.4 | 0.73 | 0.00 | 5.25 (0.11) |
| **C2** | 500 | 95.0 | 6.75 | 3.44 | -0.03 | 3.50 | 12.16 | 40.2 | 0.0 | 0.0 | 3.1 | 0.27 | 0.00 | 0.93 (0.03) |
| (0.330) | 750 | 96.0 | 5.51 | 2.81 | 0.01 | 2.79 | 7.75 | 58.1 | 0.0 | 0.0 | 2.8 | 0.39 | 0.00 | 1.64 (0.02) |
| | 1000 | 95.0 | 4.75 | 2.42 | 0.13 | 2.46 | 6.02 | 74.8 | 0.0 | 0.0 | 3.2 | 0.50 | 0.00 | 2.73 (0.08) |
| | 1500 | 92.5 | 3.88 | 1.98 | 0.39 | 1.98 | 4.05 | 103.9 | 0.0 | 0.0 | 5.5 | 0.69 | 0.00 | 2.71 (0.06) |
| | 2000 | 97.0 | 3.36 | 1.72 | 0.13 | 1.63 | 2.65 | 130.4 | 0.0 | 0.0 | 4.0 | 0.87 | 0.00 | 4.25 (0.03) |
| | 3000 | 94.0 | 2.75 | 1.40 | -0.11 | 1.40 | 1.96 | 140.9 | 0.0 | 0.0 | 3.9 | 0.94 | 0.00 | 6.09 (0.08) |
| **C3** | 500 | 94.5 | 3.68 | 1.88 | 0.27 | 1.85 | 3.49 | 25.9 | 17.8 | 0.0 | 0.0 | 0.17 | 0.01 | 1.13 (0.03) |
| (0.045) | 750 | 93.0 | 3.03 | 1.55 | 0.25 | 1.56 | 2.50 | 37.4 | 24.1 | 0.0 | 0.0 | 0.25 | 0.02 | 1.81 (0.03) |
| | 1000 | 92.5 | 2.57 | 1.31 | 0.10 | 1.41 | 1.99 | 47.4 | 30.6 | 0.0 | 0.0 | 0.32 | 0.02 | 1.58 (0.02) |
| | 1500 | 95.0 | 2.10 | 1.07 | 0.18 | 1.09 | 1.21 | 66.8 | 43.5 | 0.0 | 0.0 | 0.45 | 0.03 | 2.7 (0.04) |
| | 2000 | 95.0 | 1.79 | 0.92 | 0.11 | 0.93 | 0.87 | 81.4 | 59.8 | 0.0 | 0.0 | 0.54 | 0.04 | 3.5 (0.08) |
| | 3000 | 96.0 | 1.47 | 0.75 | 0.16 | 0.71 | 0.52 | 97.4 | 104.6 | 0.0 | 0.0 | 0.65 | 0.08 | 6.3 (0.08) |
| **C4** | 500 | 94.0 | 6.73 | 3.43 | 0.13 | 3.55 | 12.53 | 12.5 | 0.0 | 30.9 | 0.0 | 0.08 | 0.02 | 0.45 (0.01) |
| (0.302) | 750 | 96.0 | 5.48 | 2.80 | 0.16 | 2.75 | 7.55 | 21.8 | 0.0 | 38.6 | 0.0 | 0.15 | 0.03 | 1.73 (0.03) |
| | 1000 | 95.0 | 4.74 | 2.42 | -0.05 | 2.44 | 5.94 | 32.3 | 0.0 | 43.5 | 0.0 | 0.22 | 0.03 | 1.2 (0.01) |
| | 1500 | 95.5 | 3.86 | 1.97 | -0.23 | 1.96 | 3.89 | 53.1 | 0.0 | 49.2 | 0.0 | 0.35 | 0.04 | 1.74 (0.03) |
| | 2000 | 92.0 | 3.35 | 1.71 | 0.14 | 1.78 | 3.16 | 74.6 | 0.0 | 44.0 | 0.0 | 0.50 | 0.03 | 2.47 (0.06) |
| | 3000 | 97.0 | 2.74 | 1.40 | 0.15 | 1.30 | 1.71 | 110.9 | 0.0 | 18.7 | 0.0 | 0.74 | 0.01 | 3.69 (0.07) |
| **C5** | 500 | 94.0 | 6.38 | 3.25 | 0.25 | 3.25 | 10.55 | 33.9 | 4.5 | 0.0 | 4.8 | 0.23 | 0.01 | 0.84 (0.02) |
| (0.204) | 750 | 95.5 | 5.19 | 2.65 | 0.01 | 2.51 | 6.26 | 47.6 | 8.2 | 0.0 | 4.9 | 0.32 | 0.01 | 0.66 (0.02) |
| | 1000 | 94.5 | 4.47 | 2.28 | 0.18 | 2.23 | 4.96 | 60.2 | 12.5 | 0.0 | 4.8 | 0.40 | 0.01 | 0.59 (0.01) |
| | 1500 | 97.0 | 3.63 | 1.85 | 0.22 | 1.73 | 3.01 | 83.3 | 22.5 | 0.0 | 4.6 | 0.56 | 0.02 | 1.3 (0.03) |
| | 2000 | 96.0 | 3.13 | 1.60 | 0.10 | 1.54 | 2.38 | 103.1 | 33.6 | 0.0 | 4.9 | 0.69 | 0.03 | 1.66 (0.03) |
| | 3000 | 97.5 | 2.57 | 1.31 | 0.07 | 1.22 | 1.48 | 130.0 | 64.6 | 0.0 | 7.1 | 0.87 | 0.05 | 3.61 (0.03) |
| **C6** | 500 | 94.5 | 5.80 | 2.96 | 0.06 | 3.09 | 9.52 | 24.4 | 3.8 | 12.9 | 2.3 | 0.16 | 0.01 | 0.91 (0.02) |
| (0.150) | 750 | 95.5 | 4.73 | 2.41 | 0.20 | 2.37 | 5.64 | 33.5 | 7.6 | 16.7 | 3.0 | 0.22 | 0.02 | 0.74 (0.02) |
| | 1000 | 95.0 | 4.09 | 2.09 | 0.09 | 2.12 | 4.48 | 42.0 | 11.9 | 19.9 | 3.3 | 0.28 | 0.03 | 1.57 (0.03) |
| | 1500 | 92.5 | 3.33 | 1.70 | 0.11 | 1.79 | 3.19 | 57.3 | 21.2 | 26.1 | 4.0 | 0.38 | 0.04 | 1.32 (0.02) |
| | 2000 | 95.0 | 2.89 | 1.47 | 0.06 | 1.45 | 2.10 | 70.0 | 31.6 | 31.8 | 4.6 | 0.47 | 0.05 | 1.8 (0.04) |
| | 3000 | 96.5 | 2.36 | 1.20 | 0.04 | 1.20 | 1.42 | 92.6 | 54.3 | 39.6 | 7.5 | 0.62 | 0.08 | 3.21 (0.03) |

$N$ refers to sample size. CP refers to coverage probability based on 200 replications. Width refers to half the width of the 95% confidence interval. SE refers to the average asymptotic standard error. SD refers to the empirical standard deviation of replicated estimations. MSE refers to mean squared error. $(\boldsymbol{M}_{\mathcal{T}}, \boldsymbol{M}_{\mathcal{I}_1}, \boldsymbol{M}_{\mathcal{I}_2}, \boldsymbol{M}_{\mathcal{I}_3})$ refer to the number of selected true mediators, three types of non-mediators respectively. TP refers to the average true positive rate. FP refers to the average false positive rate. True value of $R^2_{Med}$ is listed within the parentheses. Time refers to the average computational time in minutes and its standard error is listed within the parentheses. The computational times for CF-OLS is observed using a single core.

Information Criterion (AIC) to fine-tune the regularization parameters within the iSIS-Lasso framework. Subsequently, we increased the maximum number of iterations for iSIS from 3 to 10, with the results presented in Table 11. Finally, due to the reduced computation time for the coverage probability, it is feasible and straightforward to increase the number of replications to 500 when using the CF-OLS method. The outcomes of this adjustment are documented in Table 12.

Table 10: Simulation statistics under uniform distribution $N(0, 1.5^2)$ setting using AIC with Lasso penalty. The maximum number of iterations for iSIS is set to 3.

| **Scen** ($R^2_{Med}$) | **N** | **CP** % | **Width** ($\times 10^{-2}$) | **SE** ($10^{-2}$) | **Bias** ($10^{-2}$) | **SD** ($10^{-2}$) | **MSE** ($10^{-4}$) | $M_{\mathcal{T}}$ | $M_{\mathcal{I}_1}$ | $M_{\mathcal{I}_2}$ | $M_{\mathcal{I}_3}$ | **TP** | **FP** | **Time** |
|---|---|---|---|---|---|---|---|---|---|---|---|---|---|---|
| **A1** | 500 | 92.5 | 4.50 | 2.30 | 0.89 | 2.34 | 6.25 | 14.6 | 0.0 | 0.0 | 28.2 | 0.97 | 0.02 | 0.58 (0.02) |
| (0.065) | 750 | 93.0 | 3.67 | 1.87 | 0.59 | 1.92 | 4.02 | 14.9 | 0.0 | 0.0 | 44.9 | 0.99 | 0.03 | 1.74 (0.03) |
| | 1000 | 93.5 | 3.18 | 1.62 | 0.50 | 1.69 | 3.08 | 14.9 | 0.0 | 0.0 | 61.0 | 1.00 | 0.04 | 2.18 (0.04) |
| | 1500 | 95.5 | 2.61 | 1.33 | 0.43 | 1.31 | 1.90 | 15.0 | 0.0 | 0.0 | 93.3 | 1.00 | 0.06 | 5.3 (0.08) |
| | 2000 | 91.0 | 2.24 | 1.14 | 0.21 | 1.31 | 1.75 | 15.0 | 0.0 | 0.0 | 123.8 | 1.00 | 0.08 | 3.62 (0.03) |
| | 3000 | 93.5 | 1.84 | 0.94 | 0.23 | 0.99 | 1.03 | 15.0 | 0.0 | 0.0 | 184.6 | 1.00 | 0.12 | 8.85 (0.11) |
| **A2** | 500 | 94.5 | 6.59 | 3.36 | -0.12 | 3.46 | 11.94 | 43.4 | 0.0 | 0.0 | 0.5 | 0.29 | 0.00 | 0.68 (0.02) |
| (0.418) | 750 | 95.5 | 5.39 | 2.75 | -0.07 | 2.74 | 7.48 | 61.4 | 0.0 | 0.0 | 0.5 | 0.41 | 0.00 | 2.46 (0.03) |
| | 1000 | 95.0 | 4.65 | 2.37 | 0.07 | 2.40 | 5.74 | 78.2 | 0.0 | 0.0 | 0.7 | 0.52 | 0.00 | 1.82 (0.03) |
| | 1500 | 92.0 | 3.79 | 1.93 | 0.33 | 1.96 | 3.92 | 108.0 | 0.0 | 0.0 | 3.9 | 0.72 | 0.00 | 2.91 (0.05) |
| | 2000 | 96.5 | 3.29 | 1.68 | 0.11 | 1.58 | 2.48 | 126.9 | 0.0 | 0.0 | 15.5 | 0.85 | 0.01 | 5.22 (0.11) |
| | 3000 | 94.5 | 2.69 | 1.37 | -0.13 | 1.39 | 1.94 | 148.9 | 0.0 | 0.0 | 49.2 | 0.99 | 0.04 | 8.68 (0.09) |
| **A3** | 500 | 96.0 | 4.27 | 2.18 | 0.23 | 2.10 | 4.46 | 34.6 | 9.4 | 0.0 | 0.0 | 0.23 | 0.01 | 1.05 (0.05) |
| (0.064) | 750 | 94.0 | 3.50 | 1.78 | 0.26 | 1.79 | 3.27 | 47.9 | 14.1 | 0.0 | 0.0 | 0.32 | 0.01 | 1.72 (0.04) |
| | 1000 | 95.0 | 2.97 | 1.52 | 0.08 | 1.62 | 2.61 | 59.2 | 19.7 | 0.0 | 0.0 | 0.39 | 0.01 | 1.49 (0.04) |
| | 1500 | 95.0 | 2.43 | 1.24 | 0.19 | 1.26 | 1.61 | 79.3 | 32.4 | 0.0 | 0.0 | 0.53 | 0.02 | 3.41 (0.10) |
| | 2000 | 94.5 | 2.09 | 1.07 | 0.11 | 1.07 | 1.16 | 96.5 | 46.2 | 0.0 | 0.0 | 0.64 | 0.03 | 4.45 (0.09) |
| | 3000 | 95.0 | 1.71 | 0.87 | 0.17 | 0.82 | 0.69 | 119.4 | 84.1 | 0.0 | 0.0 | 0.80 | 0.06 | 7.86 (0.13) |
| **A4** | 500 | 94.5 | 6.68 | 3.41 | 0.10 | 3.52 | 12.34 | 11.4 | 0.0 | 32.6 | 0.0 | 0.08 | 0.02 | 0.74 (0.02) |
| (0.390) | 750 | 96.0 | 5.44 | 2.77 | 0.12 | 2.74 | 7.46 | 17.8 | 0.0 | 44.0 | 0.0 | 0.12 | 0.03 | 0.89 (0.03) |
| | 1000 | 95.0 | 4.71 | 2.40 | -0.04 | 2.45 | 5.97 | 25.0 | 0.0 | 53.6 | 0.0 | 0.17 | 0.04 | 2.03 (0.07) |
| | 1500 | 95.5 | 3.85 | 1.96 | -0.26 | 1.96 | 3.90 | 39.2 | 0.0 | 71.6 | 0.0 | 0.26 | 0.05 | 4.45 (0.10) |
| | 2000 | 92.5 | 3.32 | 1.70 | 0.09 | 1.78 | 3.17 | 51.6 | 0.0 | 88.7 | 0.0 | 0.34 | 0.07 | 4.35 (0.04) |
| | 3000 | 97.0 | 2.72 | 1.39 | 0.12 | 1.30 | 1.70 | 72.4 | 0.0 | 125.0 | 0.0 | 0.48 | 0.09 | 9.07 (0.10) |
| **A5** | 500 | 93.5 | 6.69 | 3.41 | 0.20 | 3.43 | 11.77 | 39.9 | 2.3 | 0.0 | 1.7 | 0.27 | 0.00 | 1.51 (0.04) |
| (0.271) | 750 | 96.0 | 5.45 | 2.78 | -0.02 | 2.63 | 6.90 | 55.0 | 5.0 | 0.0 | 1.9 | 0.37 | 0.01 | 1.02 (0.02) |
| | 1000 | 94.0 | 4.71 | 2.40 | 0.17 | 2.32 | 5.39 | 68.3 | 8.3 | 0.0 | 2.2 | 0.46 | 0.01 | 2.69 (0.04) |
| | 1500 | 97.0 | 3.83 | 1.96 | 0.18 | 1.82 | 3.32 | 91.1 | 17.8 | 0.0 | 2.8 | 0.61 | 0.02 | 3.73 (0.08) |
| | 2000 | 96.0 | 3.31 | 1.69 | 0.08 | 1.61 | 2.58 | 109.7 | 29.4 | 0.0 | 3.8 | 0.73 | 0.02 | 4.16 (0.10) |
| | 3000 | 97.0 | 2.71 | 1.38 | 0.05 | 1.29 | 1.66 | 141.3 | 55.8 | 0.0 | 6.7 | 0.94 | 0.05 | 7.6 (0.10) |
| **A6** | 500 | 93.0 | 6.67 | 3.40 | -0.08 | 3.55 | 12.57 | 25.9 | 1.4 | 16.3 | 0.3 | 0.17 | 0.01 | 1.04 (0.02) |
| (0.377) | 750 | 96.5 | 5.45 | 2.78 | 0.04 | 2.74 | 7.48 | 35.5 | 3.8 | 22.0 | 0.6 | 0.24 | 0.02 | 1.84 (0.03) |
| | 1000 | 95.5 | 4.74 | 2.42 | -0.05 | 2.50 | 6.23 | 44.5 | 6.5 | 27.0 | 0.7 | 0.30 | 0.03 | 0.92 (0.02) |
| | 1500 | 93.0 | 3.86 | 1.97 | 0.05 | 2.12 | 4.47 | 59.7 | 12.9 | 37.6 | 1.2 | 0.40 | 0.04 | 3.38 (0.07) |
| | 2000 | 95.0 | 3.35 | 1.71 | -0.01 | 1.71 | 2.89 | 72.5 | 21.1 | 46.7 | 1.7 | 0.48 | 0.05 | 4.67 (0.13) |
| | 3000 | 95.5 | 2.74 | 1.40 | -0.03 | 1.39 | 1.92 | 93.9 | 41.6 | 61.7 | 4.5 | 0.63 | 0.08 | 8.4 (0.19) |

$N$ refers to sample size. CP refers to coverage probability based on 200 replications. Width refers to half the width of the 95% confidence interval. SE refers to the average asymptotic standard error. SD refers to the empirical standard deviation of replicated estimations. MSE refers to mean squared error. $(\boldsymbol{M}_{\mathcal{T}}, \boldsymbol{M}_{\mathcal{I}_1}, \boldsymbol{M}_{\mathcal{I}_2}, \boldsymbol{M}_{\mathcal{I}_3})$ refer to the number of selected true mediators, three types of non-mediators respectively. TP refers to the average true positive rate. FP refers to the average false positive rate. True value of $R^2_{Med}$ is listed within the parentheses. Time refers to the average computational time in minutes and its standard error is listed within the parentheses. The computational times for CF-OLS is observed using a single core.

Table 11: Simulation statistics under normal distribution $N(0, 1.5^2)$ setting using BIC with MCP penalty. The maximum number of iterations for iSIS is set to 10.

| **Scen** ($R^2_{Med}$) | **N** | **CP** % | **Width** ($\times 10^{-2}$) | **SE** ($10^{-2}$) | **Bias** ($10^{-2}$) | **SD** ($10^{-2}$) | **MSE** ($10^{-4}$) | $M_{\mathcal{T}}$ | $M_{\mathcal{I}_1}$ | $M_{\mathcal{I}_2}$ | $M_{\mathcal{I}_3}$ | **TP** | **FP** | **Time** |
|---|---|---|---|---|---|---|---|---|---|---|---|---|---|---|
| **A1** | 500 | 93.0 | 4.49 | 2.29 | 1.04 | 2.37 | 6.67 | 14.1 | 0.0 | 0.0 | 28.3 | 0.94 | 0.02 | 2.7 (0.05) |
| (0.065) | 750 | 91.5 | 3.66 | 1.87 | 0.75 | 1.93 | 4.27 | 14.2 | 0.0 | 0.0 | 40.2 | 0.95 | 0.03 | 2.21 (0.04) |
| | 1000 | 93.5 | 3.17 | 1.62 | 0.77 | 1.72 | 3.51 | 14.1 | 0.0 | 0.0 | 44.0 | 0.94 | 0.03 | 4.56 (0.08) |
| | 1500 | 92.5 | 2.60 | 1.33 | 0.75 | 1.35 | 2.38 | 13.8 | 0.0 | 0.0 | 22.0 | 0.92 | 0.01 | 6.5 (0.14) |
| | 2000 | 90.5 | 2.24 | 1.14 | 0.20 | 1.32 | 1.78 | 14.3 | 0.0 | 0.0 | 25.0 | 0.95 | 0.02 | 6.92 (0.19) |
| | 3000 | 93.5 | 1.84 | 0.94 | 0.13 | 0.99 | 1.00 | 14.5 | 0.0 | 0.0 | 12.1 | 0.97 | 0.01 | 8.54 (0.16) |
| **A2** | 500 | 94.0 | 6.57 | 3.35 | -0.05 | 3.44 | 11.81 | 43.2 | 0.0 | 0.0 | 0.9 | 0.29 | 0.00 | 1.66 (0.04) |
| (0.418) | 750 | 94.5 | 5.38 | 2.75 | -0.02 | 2.73 | 7.42 | 61.3 | 0.0 | 0.0 | 0.8 | 0.41 | 0.00 | 2.69 (0.06) |
| | 1000 | 95.5 | 4.64 | 2.37 | 0.10 | 2.40 | 5.72 | 77.9 | 0.0 | 0.0 | 1.2 | 0.52 | 0.00 | 3.38 (0.07) |
| | 1500 | 92.0 | 3.79 | 1.93 | 0.34 | 1.96 | 3.92 | 109.1 | 0.0 | 0.0 | 2.6 | 0.73 | 0.00 | 8.26 (0.25) |
| | 2000 | 96.5 | 3.29 | 1.68 | 0.11 | 1.58 | 2.48 | 140.1 | 0.0 | 0.0 | 1.0 | 0.93 | 0.00 | 12.23 (0.14) |
| | 3000 | 94.5 | 2.69 | 1.37 | -0.13 | 1.39 | 1.94 | 141.3 | 0.0 | 0.0 | 3.5 | 0.94 | 0.00 | 12.64 (0.20) |
| **A3** | 500 | 96.0 | 4.27 | 2.18 | 0.23 | 2.10 | 4.46 | 34.2 | 9.8 | 0.0 | 0.0 | 0.23 | 0.01 | 1.57 (0.03) |
| (0.064) | 750 | 93.5 | 3.49 | 1.78 | 0.27 | 1.79 | 3.27 | 46.6 | 15.4 | 0.0 | 0.0 | 0.31 | 0.01 | 3.11 (0.06) |
| | 1000 | 94.5 | 2.97 | 1.51 | 0.08 | 1.61 | 2.57 | 56.6 | 22.5 | 0.0 | 0.0 | 0.38 | 0.02 | 2.45 (0.06) |
| | 1500 | 95.0 | 2.43 | 1.24 | 0.20 | 1.26 | 1.62 | 74.3 | 37.8 | 0.0 | 0.0 | 0.50 | 0.03 | 7.09 (0.21) |
| | 2000 | 95.5 | 2.09 | 1.06 | 0.11 | 1.07 | 1.16 | 89.4 | 53.6 | 0.0 | 0.0 | 0.60 | 0.04 | 10.39 (0.28) |
| | 3000 | 95.0 | 1.71 | 0.87 | 0.17 | 0.81 | 0.69 | 111.7 | 92.5 | 0.0 | 0.0 | 0.74 | 0.07 | 18.78 (0.45) |
| **A4** | 500 | 95.5 | 6.69 | 3.41 | 0.06 | 3.48 | 12.05 | 11.7 | 0.0 | 32.3 | 0.0 | 0.08 | 0.02 | 2.05 (0.06) |
| (0.390) | 750 | 95.5 | 5.45 | 2.78 | 0.03 | 2.77 | 7.61 | 22.5 | 0.0 | 36.3 | 0.0 | 0.15 | 0.03 | 3.8 (0.06) |
| | 1000 | 94.5 | 4.70 | 2.40 | -0.05 | 2.43 | 5.88 | 41.6 | 0.0 | 32.4 | 0.0 | 0.28 | 0.02 | 8.52 (0.15) |
| | 1500 | 95.5 | 3.84 | 1.96 | -0.22 | 1.96 | 3.87 | 99.0 | 0.0 | 4.1 | 0.0 | 0.66 | 0.00 | 13.25 (0.12) |
| | 2000 | 92.5 | 3.32 | 1.70 | 0.10 | 1.78 | 3.16 | 115.0 | 0.0 | 0.1 | 0.0 | 0.77 | 0.00 | 13.94 (0.20) |
| | 3000 | 97.0 | 2.72 | 1.39 | 0.11 | 1.30 | 1.70 | 119.8 | 0.0 | 0.1 | 0.0 | 0.80 | 0.00 | 13.35 (0.11) |
| **A5** | 500 | 93.0 | 6.67 | 3.40 | 0.25 | 3.44 | 11.82 | 38.9 | 2.8 | 0.0 | 2.4 | 0.26 | 0.00 | 2.03 (0.04) |
| (0.271) | 750 | 96.0 | 5.44 | 2.77 | 0.04 | 2.61 | 6.80 | 53.3 | 6.1 | 0.0 | 2.7 | 0.36 | 0.01 | 3.22 (0.10) |
| | 1000 | 94.5 | 4.70 | 2.40 | 0.20 | 2.30 | 5.29 | 65.6 | 10.5 | 0.0 | 3.0 | 0.44 | 0.01 | 2.57 (0.06) |
| | 1500 | 97.0 | 3.83 | 1.96 | 0.19 | 1.82 | 3.31 | 86.8 | 21.8 | 0.0 | 3.5 | 0.58 | 0.02 | 6.71 (0.18) |
| | 2000 | 96.0 | 3.31 | 1.69 | 0.08 | 1.61 | 2.58 | 102.4 | 35.7 | 0.0 | 5.1 | 0.68 | 0.03 | 7.36 (0.15) |
| | 3000 | 97.0 | 2.71 | 1.38 | 0.05 | 1.29 | 1.66 | 130.7 | 67.2 | 0.0 | 6.3 | 0.87 | 0.05 | 17.04 (0.36) |
| **A6** | 500 | 93.0 | 6.66 | 3.40 | -0.07 | 3.55 | 12.55 | 26.1 | 1.6 | 16.0 | 0.4 | 0.17 | 0.01 | 1.62 (0.04) |
| (0.377) | 750 | 96.5 | 5.45 | 2.78 | 0.05 | 2.74 | 7.46 | 36.5 | 4.6 | 20.2 | 0.8 | 0.24 | 0.02 | 1.81 (0.06) |
| | 1000 | 95.0 | 4.74 | 2.42 | -0.04 | 2.50 | 6.22 | 46.4 | 8.8 | 22.8 | 1.1 | 0.31 | 0.02 | 2.82 (0.09) |
| | 1500 | 92.5 | 3.86 | 1.97 | 0.05 | 2.11 | 4.45 | 64.0 | 23.8 | 22.2 | 2.1 | 0.43 | 0.04 | 4.38 (0.08) |
| | 2000 | 95.0 | 3.35 | 1.71 | 0.00 | 1.71 | 2.90 | 78.0 | 42.6 | 19.4 | 3.1 | 0.52 | 0.05 | 5.91 (0.11) |
| | 3000 | 95.5 | 2.74 | 1.40 | -0.02 | 1.39 | 1.92 | 104.9 | 81.0 | 9.7 | 8.0 | 0.70 | 0.07 | 16.56 (0.25) |

$N$ refers to sample size. CP refers to coverage probability based on 200 replications. Width refers to half the width of the 95% confidence interval. SE refers to the average asymptotic standard error. SD refers to the empirical standard deviation of replicated estimations. MSE refers to mean squared error. $(\boldsymbol{M}_{\mathcal{T}}, \boldsymbol{M}_{\mathcal{I}_1}, \boldsymbol{M}_{\mathcal{I}_2}, \boldsymbol{M}_{\mathcal{I}_3})$ refer to the number of selected true mediators, three types of non-mediators respectively. TP refers to the average true positive rate. FP refers to the average false positive rate. True value of $R^2_{Med}$ is listed within the parentheses. Time refers to the average computational time in minutes and its standard error is listed within the parentheses. The computational times for CF-OLS is observed using a single core.

Table 12: Simulation statistics under normal distribution $N(0, 1.5^2)$ setting using BIC with MCP penalty. The maximum number of iterations for iSIS is set to 3.

| **Scen** ($R^2_{Med}$) | **N** | **CP** % | **Width** ($\times 10^{-2}$) | **SE** ($10^{-2}$) | **Bias** ($10^{-2}$) | **SD** ($10^{-2}$) | **MSE** ($10^{-4}$) | $M_{\mathcal{T}}$ | $M_{\mathcal{I}_1}$ | $M_{\mathcal{I}_2}$ | $M_{\mathcal{I}_3}$ | **TP** | **FP** | **Time** |
|---|---|---|---|---|---|---|---|---|---|---|---|---|---|---|
| **A1** | 500 | 94.2 | 4.48 | 2.28 | 1.00 | 2.29 | 6.22 | 14.1 | 0.0 | 0.0 | 21.9 | 0.94 | 0.01 | 0.94 (0.05) |
| (0.065) | 750 | 93.4 | 3.66 | 1.87 | 0.73 | 1.92 | 4.20 | 14.2 | 0.0 | 0.0 | 31.9 | 0.94 | 0.02 | 1.7 (0.07) |
| | 1000 | 93.2 | 3.17 | 1.62 | 0.74 | 1.71 | 3.46 | 14.0 | 0.0 | 0.0 | 34.2 | 0.94 | 0.02 | 2.92 (0.04) |
| | 1500 | 94.0 | 2.59 | 1.32 | 0.57 | 1.34 | 2.11 | 13.9 | 0.0 | 0.0 | 25.1 | 0.93 | 0.02 | 3.49 (0.04) |
| | 2000 | 93.0 | 2.25 | 1.15 | 0.23 | 1.25 | 1.62 | 14.3 | 0.0 | 0.0 | 26.2 | 0.95 | 0.02 | 5.15 (0.05) |
| | 3000 | 94.8 | 1.84 | 0.94 | 0.12 | 0.98 | 0.96 | 14.5 | 0.0 | 0.0 | 11.7 | 0.97 | 0.01 | 7.65 (0.11) |
| **A2** | 500 | 94.8 | 6.59 | 3.36 | 0.10 | 3.37 | 11.37 | 42.6 | 0.0 | 0.0 | 0.8 | 0.28 | 0.00 | 1.34 (0.04) |
| (0.418) | 750 | 94.4 | 5.36 | 2.74 | 0.15 | 2.73 | 7.47 | 60.5 | 0.0 | 0.0 | 0.8 | 0.40 | 0.00 | 2.65 (0.04) |
| | 1000 | 95.6 | 4.64 | 2.37 | 0.14 | 2.44 | 5.97 | 76.8 | 0.0 | 0.0 | 1.2 | 0.51 | 0.00 | 2.84 (0.07) |
| | 1500 | 95.0 | 3.79 | 1.94 | 0.19 | 1.88 | 3.56 | 104.2 | 0.0 | 0.0 | 4.0 | 0.69 | 0.00 | 3.43 (0.05) |
| | 2000 | 96.0 | 3.29 | 1.68 | 0.11 | 1.65 | 2.73 | 126.3 | 0.0 | 0.0 | 6.1 | 0.84 | 0.00 | 5.87 (0.11) |
| | 3000 | 94.0 | 2.69 | 1.37 | 0.01 | 1.41 | 1.97 | 141.5 | 0.0 | 0.0 | 3.6 | 0.94 | 0.00 | 7.88 (0.08) |
| **A3** | 500 | 95.4 | 4.29 | 2.19 | 0.39 | 2.17 | 4.85 | 34.2 | 9.6 | 0.0 | 0.0 | 0.23 | 0.01 | 1.31 (0.03) |
| (0.064) | 750 | 93.6 | 3.48 | 1.77 | 0.23 | 1.80 | 3.27 | 46.5 | 15.0 | 0.0 | 0.0 | 0.31 | 0.01 | 1.78 (0.03) |
| | 1000 | 94.8 | 2.99 | 1.53 | 0.16 | 1.54 | 2.38 | 57.0 | 21.2 | 0.0 | 0.0 | 0.38 | 0.02 | 1.72 (0.03) |
| | 1500 | 94.8 | 2.42 | 1.24 | 0.15 | 1.25 | 1.58 | 75.8 | 35.0 | 0.0 | 0.0 | 0.51 | 0.03 | 4.3 (0.09) |
| | 2000 | 94.6 | 2.08 | 1.06 | 0.08 | 1.08 | 1.17 | 92.0 | 49.4 | 0.0 | 0.0 | 0.61 | 0.04 | 4.49 (0.08) |
| | 3000 | 96.0 | 1.70 | 0.87 | 0.08 | 0.82 | 0.69 | 113.9 | 88.3 | 0.0 | 0.0 | 0.76 | 0.07 | 8.66 (0.14) |
| **A4** | 500 | 94.0 | 6.69 | 3.41 | -0.09 | 3.47 | 12.04 | 11.5 | 0.0 | 31.4 | 0.0 | 0.08 | 0.02 | 1.58 (0.03) |
| (0.390) | 750 | 94.8 | 5.44 | 2.78 | -0.09 | 2.82 | 7.97 | 19.3 | 0.0 | 34.1 | 0.0 | 0.13 | 0.03 | 1.61 (0.03) |
| | 1000 | 95.0 | 4.71 | 2.40 | -0.02 | 2.41 | 5.78 | 30.9 | 0.0 | 33.9 | 0.0 | 0.21 | 0.03 | 2.31 (0.04) |
| | 1500 | 95.2 | 3.85 | 1.96 | -0.08 | 1.94 | 3.76 | 57.6 | 0.0 | 29.9 | 0.0 | 0.38 | 0.02 | 2.63 (0.05) |
| | 2000 | 94.2 | 3.33 | 1.70 | 0.10 | 1.74 | 3.05 | 80.8 | 0.0 | 17.5 | 0.0 | 0.54 | 0.01 | 3.55 (0.06) |
| | 3000 | 95.4 | 2.72 | 1.39 | 0.12 | 1.37 | 1.89 | 108.7 | 0.0 | 1.6 | 0.0 | 0.72 | 0.00 | 3.2 (0.11) |
| **A5** | 500 | 93.8 | 6.69 | 3.41 | 0.40 | 3.58 | 12.95 | 38.8 | 2.6 | 0.0 | 2.1 | 0.26 | 0.00 | 0.94 (0.01) |
| (0.271) | 750 | 94.4 | 5.44 | 2.78 | 0.10 | 2.76 | 7.62 | 52.9 | 5.7 | 0.0 | 2.5 | 0.35 | 0.01 | 1.79 (0.03) |
| | 1000 | 94.6 | 4.71 | 2.40 | 0.15 | 2.40 | 5.78 | 65.3 | 9.7 | 0.0 | 2.8 | 0.44 | 0.01 | 2.1 (0.03) |
| | 1500 | 96.4 | 3.84 | 1.96 | 0.14 | 1.78 | 3.18 | 86.8 | 20.1 | 0.0 | 3.6 | 0.58 | 0.02 | 3.82 (0.06) |
| | 2000 | 95.2 | 3.32 | 1.69 | 0.09 | 1.66 | 2.75 | 102.5 | 33.4 | 0.0 | 5.3 | 0.68 | 0.03 | 4.49 (0.14) |
| | 3000 | 95.8 | 2.71 | 1.38 | 0.07 | 1.35 | 1.82 | 131.5 | 61.5 | 0.0 | 7.5 | 0.88 | 0.05 | 7.38 (0.11) |
| **A6** | 500 | 94.0 | 6.69 | 3.41 | 0.01 | 3.63 | 13.14 | 25.8 | 1.6 | 15.8 | 0.4 | 0.17 | 0.01 | 0.78 (0.01) |
| (0.377) | 750 | 95.2 | 5.46 | 2.79 | 0.21 | 2.88 | 8.32 | 35.7 | 4.2 | 20.3 | 0.8 | 0.24 | 0.02 | 1.23 (0.04) |
| | 1000 | 95.2 | 4.74 | 2.42 | 0.09 | 2.44 | 5.93 | 44.6 | 7.7 | 23.6 | 1.0 | 0.30 | 0.02 | 2.26 (0.05) |
| | 1500 | 93.8 | 3.86 | 1.97 | 0.12 | 2.04 | 4.17 | 59.9 | 17.8 | 26.2 | 1.9 | 0.40 | 0.03 | 3.45 (0.04) |
| | 2000 | 94.0 | 3.35 | 1.71 | 0.06 | 1.75 | 3.06 | 73.0 | 32.7 | 25.5 | 3.3 | 0.49 | 0.05 | 5.36 (0.11) |
| | 3000 | 94.4 | 2.73 | 1.40 | 0.01 | 1.39 | 1.92 | 93.5 | 62.2 | 21.1 | 13.6 | 0.62 | 0.07 | 9.27 (0.13) |

$N$ refers to sample size. CP refers to coverage probability based on 500 replications. Width refers to half the width of the 95% confidence interval. SE refers to the average asymptotic standard errors. SD refers to the empirical standard deviation of replicated estimations. MSE refers to mean squared error. $(M_{\mathcal{T}}, M_{\mathcal{I}_1}, M_{\mathcal{I}_2}, M_{\mathcal{I}_3})$ refer to the number of selected true mediators, three types of non-mediators respectively. TP refers to the average true positive rate. FP refers to the average false positive rate. True value of $R^2_{Med}$ is listed within the parentheses. Time refers to computational time and its standard error is listed within the parentheses.

### 4 Web Appendix D: Details and Supplement of the Application for Systolic BP and HDL-C in FHS

In this section, we carried out data cleaning and descriptive statistics for the FHS dataset in preparation for applying the CF-OLS method. Additionally, we present the actual correlation matrix of selected mediators before and after FDR control in two subsamples for the two outcomes. Finally, we conducted pathway enrichment analysis for the selected genes. To conclude, we illustrate the biological significance of our novel findings and the reaffirmation of previous findings.

#### 4.1 Descriptive statistics

In this section, we provide background information on the insights gathered from analyzing the FHS datasets with our proposed CF-OLS method and the B-Mixed approach. We summarize the descriptive statistics in Table 13 for the systolic BP outcome and in Table 14 for the HDL-C outcome. We have labeled the exposure variable, the outcome variable, and the covariates to elucidate their roles in each analysis. The slight differences between the two analytical samples are primarily attributable to the missingness in systolic BP and HDL-C data.

In Figure 3, we display the correlations between mediators selected before and after FDR control. The green labels indicate mediators selected after iSIS-MCP and retained post-FDR control, while the blue labels denote mediators chosen after iSIS-MCP but filtered out by FDR control. Here we did not adjust the top PCs as the covariates. After adjusting the first 10 PCs as the covariates, we obtain in Figure 4, the correlations decreased a lot among all the potential mediators.

Table 15 presents a comparison of the mediation effect size results for systolic blood pressure (BP) outcomes in the Framingham Heart Study (FHS) Offspring cohort, utilizing both the B-Mixed and CF-OLS methods. This cohort closely mirrors that used in the study by Yang et al. [2021]. To ensure comparability of the results, we adjusted the same covariates for both methods and employed the same random seed for consistent sample splitting. The findings indicate that the CF-OLS method yields higher estimates for $R^2_{Med}$ and SOS measure, while with fewer mediators selected. Additionally, the CF-OLS method benefits from a narrower confidence interval, attributed to the utilization of the entire sample rather than just a subset, thereby leveraging the full data set to enhance estimation precision.

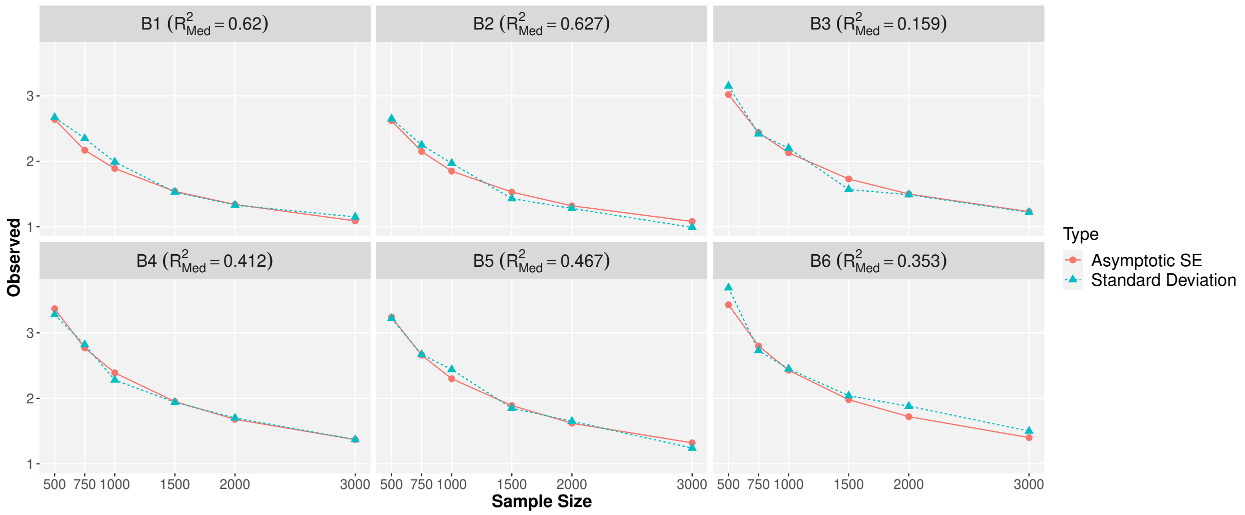


Figure 1: Plots of asymptotic standard error and empirical standard deviation of estimation under normal distribution setting $Unif(-2, 2)$ using the CF-OLS estimation. SE refers to standard errors. The sample Size is increased from 500 to 3,000. The true value of $R^2_{Med}$ is listed within the parentheses.

Table 13: Descriptive statistics for Systolic BP outcome data application.

c

| Variable | Description | Type | Min | Mean | Q1 | Median | Q3 | Max | Sd |
|---|---|---|---|---|---|---|---|---|---|
| Age | Age Attended Exam | **Exposure** | 24.0 | 54.0 | 45.0 | 54.0 | 63.0 | 90.0 | 12.6 |
| SBP–adj | Adjusted Systolic blood pressure according to Hypertensive medication, mm hg | **Outcome** | 82.0 | 128.0 | 114.0 | 127.0 | 141.0 | 211.0 | 19.0 |
| BMI | Body Mass Index | Covariate | 16.3 | 28.1 | 24.2 | 27.3 | 30.9 | 65.6 | 5.5 |
| | | | **1** | **0** | | | | | |
| Sex | Participant gender (1 = Male, 0 = Female) | Covariate | 2086 | 2456 | | | | | |
| Smoke | Current smoking status (1 = Current, 0 = Not a current smoker) | Covariate | 389 | 4153 | | | | | |
| Alcohol | Drinking Status (1 = Current or Stopped, 0 = Never) | Covariate | 4231 | 311 | | | | | |
| Cohort | Cohort (1 = Offspring, 0 = Generation 3) | Covariate | 1871 | 2671 | | | | | |

Q1 and Q3 refer to the first and third quartile, respectively. Min and Max refer to the lowest and highest observation, respectively. Sd refers to standard deviation.

Table 14: Descriptive statistics for HDL-C outcome data application.

c

| Variable | Description | Type | Min | Mean | Q1 | Median | Q3 | Max | Sd |
|---|---|---|---|---|---|---|---|---|---|
| Age | Age Attended Exam | Covariate | 24.0 | 53.8 | 45.0 | 53.0 | 63.0 | 88.0 | 12.4 |
| HDL-C | HDL cholesterol (sample type: EDTA plasma), mg/dL | **Outcome** | 7.0 | 60.5 | 46.0 | 58.0 | 72.0 | 198.0 | 19.3 |
| BMI | Body Mass Index | Covariate | 16.3 | 28.1 | 24.2 | 27.3 | 31.0 | 65.6 | 5.6 |
| | | | **1** | **0** | | | | | |
| Sex | Participant gender (1 = Male, 0 = Female) | **Exposure** | 2058 | 2423 | | | | | |
| Smoke | Current smoking status (1 = Current, 0 = Not a current smoker) | Covariate | 382 | 4099 | | | | | |
| Alcohol | Drinking Status (1 = Current or Stopped, 0 = Never) | Covariate | 4175 | 306 | | | | | |
| Cohort | Cohort (1 = Offspring, 0 = Generation 3) | Covariate | 1829 | 2652 | | | | | |

Q1 and Q3 refer to the first and third quartile, respectively. Min and Max refer to the lowest and highest observation, respectively. Sd refers to standard deviation.

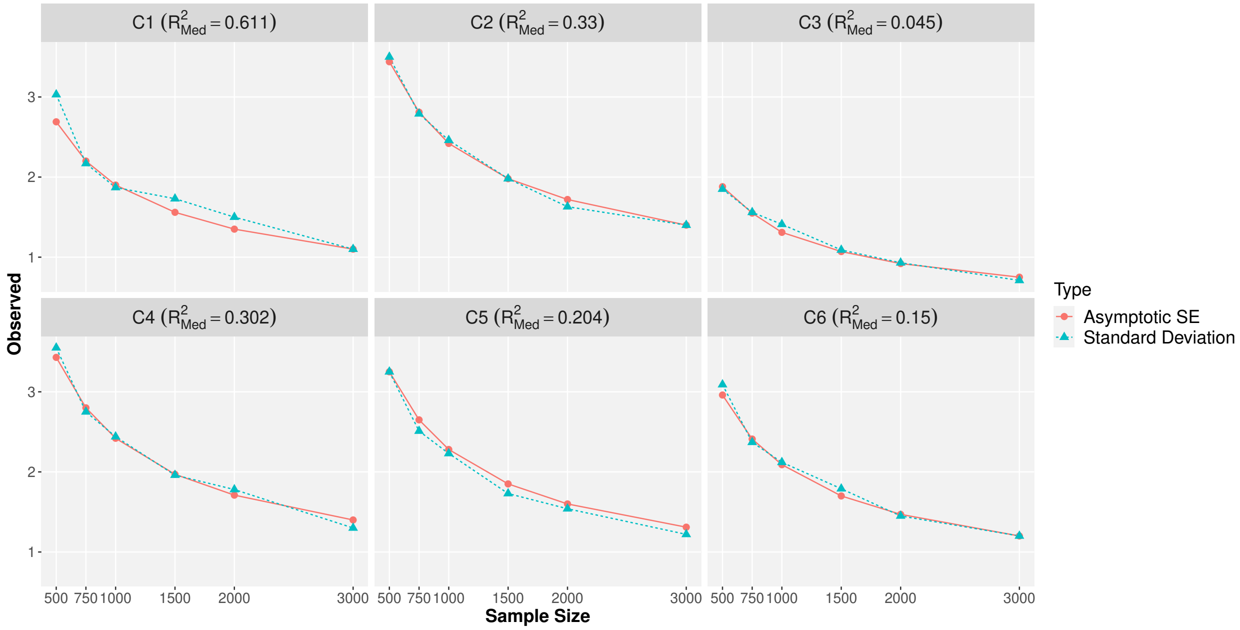


Figure 2: Plots of asymptotic standard error and empirical standard deviation of estimation under uniform distribution setting $N(0, 1^2)$ using the CF-OLS estimation. SE refers to standard errors. The sample Size is increased from 500 to 3,000. The true value of $R^2_{Med}$ is listed within the parentheses.

Figure 3: Heatmaps of the correlation of selected mediators after the iSIS-MCP procedure and the FDR control (green) and mediators filtered out by the FDR control (blue) in (A) the first subsample for systolic BP, (B) the second subsample for systolic BP, (C) the first subsample for HDL-C, and (D) the second subsample for HDL-C.

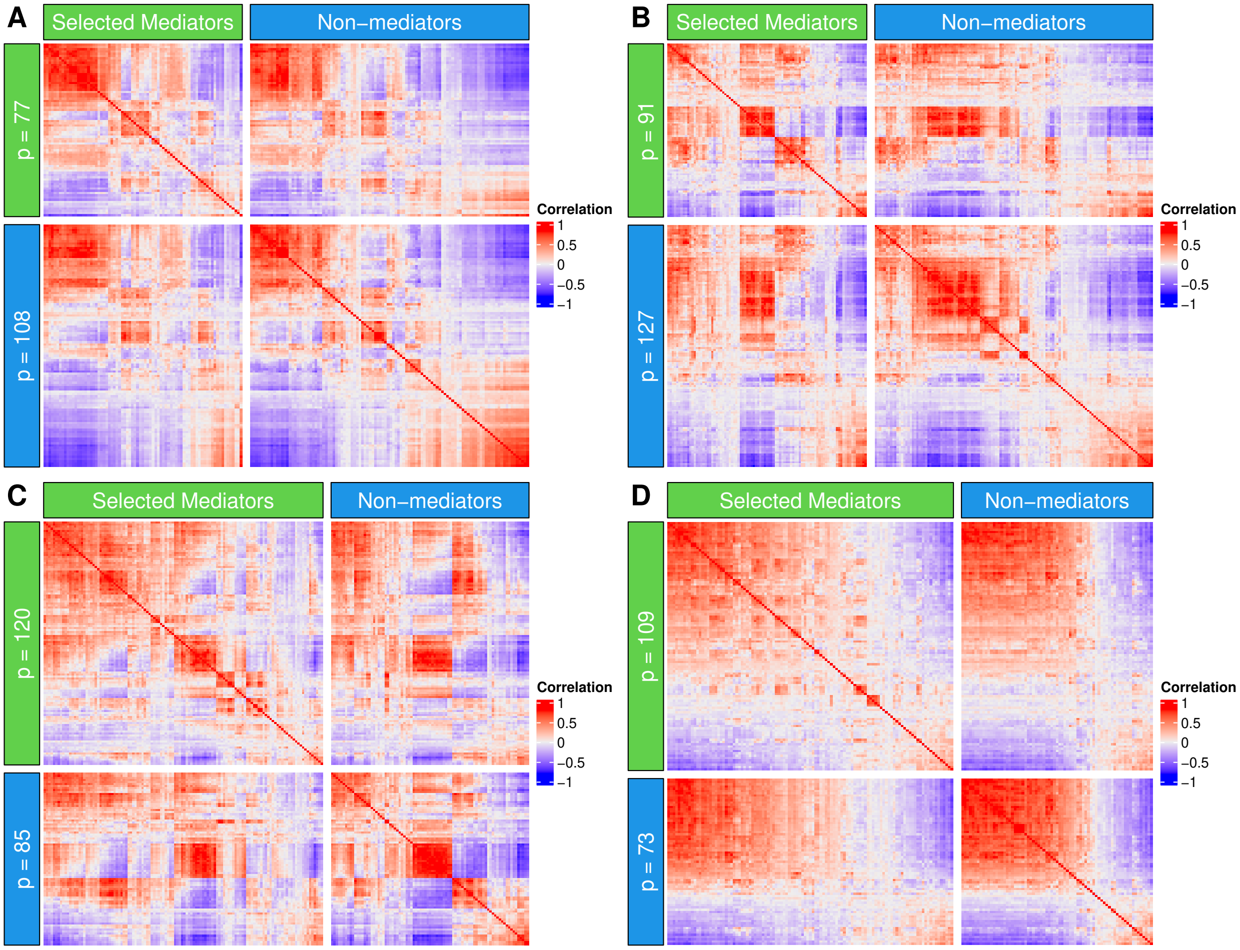

Figure 4: Heatmaps of the correlation of selected mediators (green) and mediators selected by iSIS but filtered out by the FDR control (blue) in (A) the first subsample for systolic BP, (B) the second subsample for systolic BP, (C) the first subsample for HDL-C, and (D) the second subsample for HDL-C.

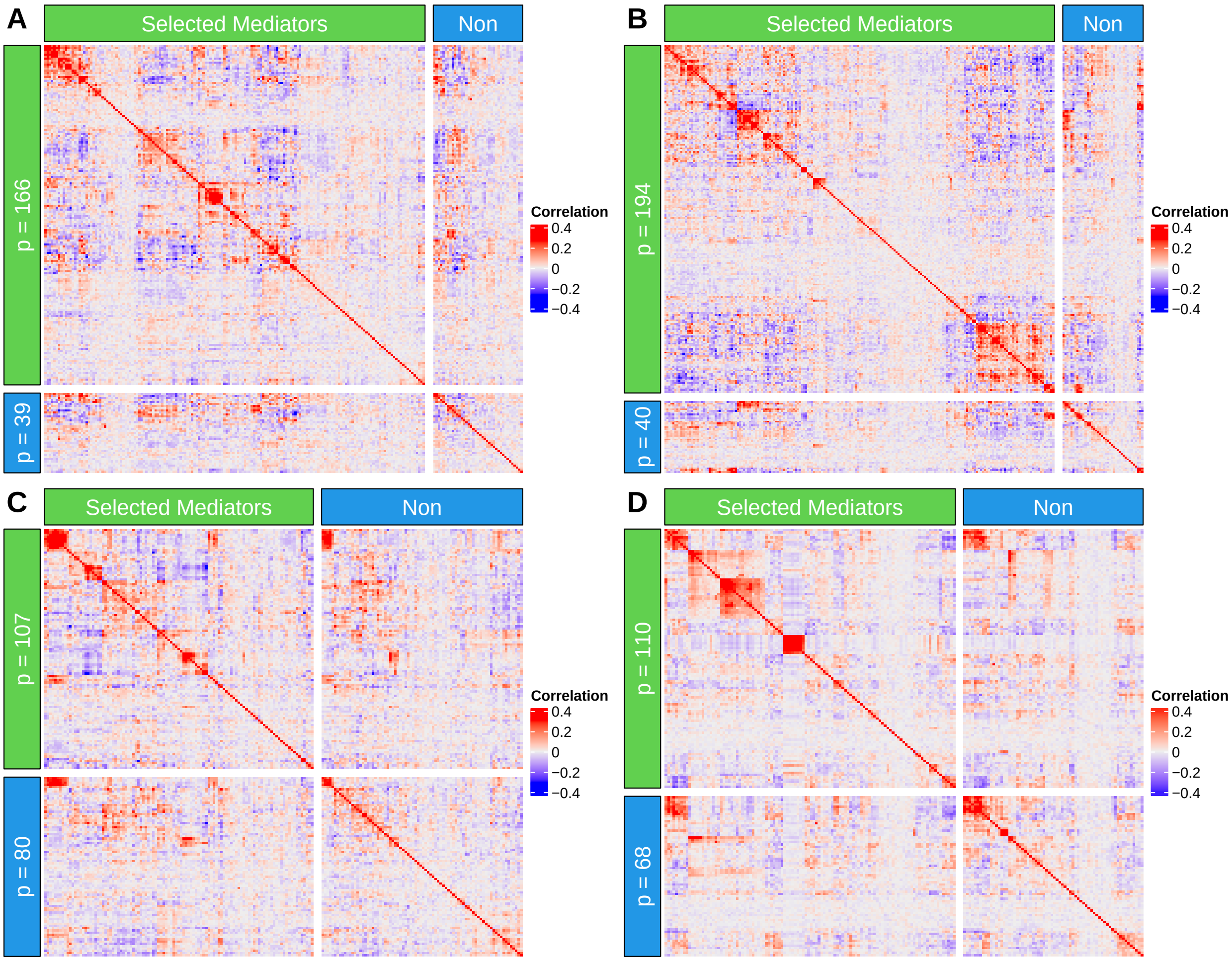

Table 15: Mediation effect size and 95% confidence interval estimated using the CF-OLS method and B-Mixed method in the Framingham Heart Study (FHS) Offspring cohort data for systolic BP outcome.

| Method | $N$ | $R^2_{Med}$ | SOS | $R^2_{Y,X}$ | ab | prop | total | p̂ |
|---|---|---|---|---|---|---|---|---|
| CF-OLS | 1888 | 0.0429<br>(0.0267, 0.0591) | 0.421<br>(0.306, 0.536) | 0.0981 | -4.853/-4.511 | -6.583/-6.161 | 0.737/0.732 | 130/131 |
| B-Mixed | 1888 | 0.0350<br>(-0.0091, 0.0695) | 0.378<br>(-0.1094, 0.6212) | 0.0924<br>(0.054, 0.139) | -4.860<br>(-6.158, -3.815) | -7.073<br>(-11.838, -4.521) | 0.687<br>(0.520, 0.844) | 167<br>(123, 194) |

**N** refers to the sample size. **ab** refers to the indirect mediation effect. **prop** refers to the proportion measure. **total** refers to the total effect. **p̂** refers to the number of genes selected. 95% confidence interval listed within the parentheses for the B-Mixed method is computed over 200 bootstrap samples. For the CF-OLS method, the splitting of data results in two sets of results for **ab**, **prop**, **total** and **p̂** across two subsamples.

#### 4.2 Discussion about the assumptions in FHS and simulations

Note that these conditions are sufficient instead of necessary, we sought to validate our theoretical assumptions using real data applications. By examining the correlations among the mediators after regressing out the principal components (PCs) and other covariates, we observed that the correlations were fairly low once adjusted for these factors to meet the Assumption 3. For the systolic blood pressure (BP) outcome (Figure 4A and Figure 4B), the $\max|\Sigma_{kj}|, k \in \mathcal{T}, j \in \mathcal{T}^c$ values are 0.556 and 0.461 for the two subsamples, respectively. The $75^{th}$ percentile of these correlations is 0.052 and 0.055, respectively. The minimum eigenvalue, $\lambda_{\min}(\mathbf{\Sigma})$, is 0.25 and 0.18, while the maximum eigenvalue, $\lambda_{\max}(\mathbf{\Sigma})$, is 6.64 and 8.37. Additionally, $\sqrt{\log(p)/n} \approx 0.046$ when $p = 17873$ and $n = 4542$.

For the HDL outcome (Figure 4C and Figure 4D), the $\max|\Sigma_{kj}|, k \in \mathcal{T}, j \in \mathcal{T}^c$ values are 0.518 and 0.724 for the two subsamples, respectively. The $75^{th}$ percentile of these correlations is 0.056 and 0.060. The minimum eigenvalue, $\lambda_{\min}(\mathbf{\Sigma})$, is 0.18 in both subsamples, while the maximum eigenvalue, $\lambda_{\max}(\mathbf{\Sigma})$, is 6.72 and 8.82. Finally, $\sqrt{\log(p)/n} \approx 0.047$ when $p = 17873$ and $n = 4481$.

We also examine the simulation settings to check whether they satisfy the conditions. In the main text for the first correlation structure, $\boldsymbol{\xi} \sim N(\mathbf{0}, \text{diag}(\mathbf{\Sigma}, \boldsymbol{I}_{p_2+p_3}))$ where $\mathbf{\Sigma}_{ij} = 0.2$ for $1 \leq i \neq j \leq p_0 + p_1$ and $\mathbf{\Sigma}_{ij} = 1$ for $1 \leq i = j \leq p_0 + p_1$. In this case, the $\max|\Sigma_{kj}|, k \in \mathcal{T}, j \in \mathcal{T}^c$ value is 0.2, which can be seen as a violation of Assumption 3. Despite this potential violation, the estimates and coverage probabilities still performed well, demonstrating the robustness of our proposed method. For the second correlation structure, we considered $\boldsymbol{\xi} \sim N(\mathbf{0}, \text{diag}(\mathbf{\Sigma}, \boldsymbol{I}_{p_2+p_3}))$ where $\mathbf{\Sigma}_{ij}$'s are iid samples from $N(0, 0.1^2)$ for $1 \leq i \neq j \leq p_0+p_1$ and $\mathbf{\Sigma}_{ij} = 1$ for $1 \leq i = j \leq p_0+p_1$. In this case, the $\max|\Sigma_{kj}|, k \in \mathcal{T}, j \in \mathcal{T}^c$ value is 0.474. The $75^{th}$ percentile of these correlations is 0.110.

For Assumption 2, we also evaluated it in the real data analysis. The mediators filtered out in the training set, where their estimated $|\alpha_j|$ and $|\beta_j|$ are zero, are consid-

Table 16: Summary statistics for the correlation $|\Sigma_{kj}|, k \in \mathcal{T}, j \in \mathcal{T}^c$ in real data analysis and simulations.

| Outcome | Statistic | min | 25% quantile | median | 75% quantile | max |
|---|---|---|---|---|---|---|
| Systolic BP | $\lvert\Sigma_{kj}\rvert$ in subsample 1 | 0.000 | 0.012 | 0.027 | 0.052 | 0.556 |
| | $\lvert\Sigma_{kj}\rvert$ in subsample 2 | 0.000 | 0.012 | 0.027 | 0.055 | 0.461 |
| HDL | $\lvert\Sigma_{kj}\rvert$ in subsample 1 | 0.000 | 0.013 | 0.029 | 0.056 | 0.518 |
| | $\lvert\Sigma_{kj}\rvert$ in subsample 2 | 0.000 | 0.013 | 0.029 | 0.060 | 0.724 |
| Simulation | $\lvert\Sigma_{kj}\rvert$ in correlation structure 1 | 0.200 | 0.200 | 0.200 | 0.200 | 0.200 |
| | $\lvert\Sigma_{kj}\rvert$ in correlation structure 2 | 0.000 | 0.035 | 0.066 | 0.110 | 0.474 |

ered non-mediators. However, their estimates in the estimation subsample using OLS are typically non-zero. In the Table 17, we observe that $|\alpha_j|$ and $|\beta_j|$ for the selected mediators are relatively large, while $|\alpha_j|$ and $|\beta_j|$ for non-selected mediators remain sufficiently small in the real data analysis for both outcomes.

In the simulation, we generated the $\alpha_j$ and $\beta_j$ for true mediators from a normal distribution $N(0, 1.5^2)$, which is sufficiently large. For the noise variables $\boldsymbol{M}_{\mathcal{I}_3}$, the $\alpha_j$ and $\beta_j$ were set to 0, ensuring they are sufficiently small to meet Assumption 2. Meanwhile, for non-mediators $\boldsymbol{M}_{\mathcal{I}_1}$ and $\boldsymbol{M}_{\mathcal{I}_2}$, the $\alpha_j$ or $\beta_j$ were also simulated from the normal distribution $N(0, 1.5^2)$. In this case, at least one of the $\alpha_j$ and $\beta_j$ was sufficiently small to meet Assumption 2.

Assumption 1 refers to the sure screening property [Fan and Lv, 2008], which ensures that the selection method can effectively reduce the dimensionality without necessarily achieving selection consistency or possessing the oracle property. As demonstrated by Fan and Lv [2008], Sure Independence Screening (SIS) possesses the sure screening property and is capable of reducing an exponentially large dimension $p$ to a more manageable size. Importantly, this reduction is done while retaining all the relevant variables in the true model with overwhelming probability.

Table 17: Summary statistics for $|\alpha_j|$, $|\beta_j|$ in real data analysis and simulations.

| Outcome | Statistic | min | 25% quantile | median | 75% quantile | max |
|---|---|---|---|---|---|---|
| Systolic BP | $|\alpha_j|$ for $j \in \mathcal{T}$ in subsample 1 | 0.002 | 0.004 | 0.006 | 0.010 | 0.026 |
| | $|\alpha_j|$ for $j \in \mathcal{T}$ in subsample 2 | 0.002 | 0.006 | 0.009 | 0.012 | 0.023 |
| | $|\alpha_j|$ for $j \in \mathcal{T}^c$ in subsample 1 | 0.000 | 0.000 | 0.002 | 0.003 | 0.006 |
| | $|\alpha_j|$ for $j \in \mathcal{T}^c$ in subsample 2 | 0.000 | 0.001 | 0.002 | 0.004 | 0.007 |
| | $|\beta_j|$ for $j \in \mathcal{T}$ in subsample 1 | 0.004 | 0.084 | 0.222 | 0.425 | 1.637 |
| | $|\beta_j|$ for $j \in \mathcal{T}$ in subsample 2 | 0.001 | 0.108 | 0.263 | 0.524 | 1.324 |
| | $|\beta_j|$ for $j \in \mathcal{T}^c$ in subsample 1 | 0.000 | 0.010 | 0.210 | 0.364 | 1.548 |
| | $|\beta_j|$ for $j \in \mathcal{T}^c$ in subsample 2 | 0.000 | 0.011 | 0.230 | 0.387 | 1.491 |
| HDL | $|\alpha_j|$ for $j \in \mathcal{T}$ in subsample 1 | 0.055 | 0.078 | 0.108 | 0.153 | 0.449 |
| | $|\alpha_j|$ for $j \in \mathcal{T}$ in subsample 2 | 0.056 | 0.081 | 0.126 | 0.215 | 0.506 |
| | $|\alpha_j|$ for $j \in \mathcal{T}^c$ in subsample 1 | 0.000 | 0.008 | 0.033 | 0.052 | 0.171 |
| | $|\alpha_j|$ for $j \in \mathcal{T}^c$ in subsample 2 | 0.000 | 0.022 | 0.044 | 0.070 | 0.163 |
| | $|\beta_j|$ for $j \in \mathcal{T}$ in subsample 1 | 0.006 | 0.161 | 0.331 | 0.510 | 3.560 |
| | $|\beta_j|$ for $j \in \mathcal{T}$ in subsample 2 | 0.006 | 0.215 | 0.446 | 0.809 | 2.862 |
| | $|\beta_j|$ for $j \in \mathcal{T}^c$ in subsample 1 | 0.000 | 0.013 | 0.027 | 0.437 | 1.666 |
| | $|\beta_j|$ for $j \in \mathcal{T}^c$ in subsample 2 | 0.000 | 0.014 | 0.028 | 0.488 | 1.380 |
| Simulation | $\boldsymbol{M}_{\mathcal{T}}$: $|\alpha_j|$ and $|\beta_j|$ | 0.001 | 0.413 | 0.968 | 1.757 | 3.717 |
| | $\boldsymbol{M}_{\mathcal{I}_1}$: $|\alpha_j|$ ($|\beta_j| = 0$) | 0.005 | 0.549 | 1.125 | 1.766 | 4.222 |
| | $\boldsymbol{M}_{\mathcal{I}_2}$: $|\beta_j|$ ($|\alpha_j| = 0$) | 0.001 | 0.384 | 0.984 | 1.683 | 3.717 |
| | $\boldsymbol{M}_{\mathcal{I}_3}$: $|\alpha_j|$ and $|\beta_j|$ | 0 | 0 | 0 | 0 | 0 |

#### 4.3 Pathway enrichment analysis results

We applied the FDR-adjusted p-value to filter out selected genes not associated with the exposures, setting the FDR cutoff point at 0.2. We conducted pathway enrichment analysis using the Database for Annotation, Visualization and Integrated Discovery (DAVID) [Dennis Jr et al., 2003] to evaluate the significance of these mediating GEs enriched in specific pathways.

Table 18 presents the pathways from the Kyoto Encyclopedia of Genes and Genomes (KEGG) [Kanehisa and Goto, 2000], BioCarta [Nishimura, 2001], and WikiPathways [Kutmon et al., 2016], which were statistically significant at a nominal significance level of 0.05. Table 19 lists the pathways identified for HDL-C.

Additionally, we calculated the product measure for the genes within the identified pathway. For instance, the MAPK pathway exhibited a positive product measure, corroborating prior research that links Aging with MAPK activity in vascular tissues. This is supported by evidence that specifically inhibiting p38 MAPK can encourage hypertrophic cardiomyopathy by enhancing calcineurin-NFAT signaling, as documented in [Braz et al., 2003].

Table 18: The significant pathways identified for systolic blood pressure. $ab$ refers to the indirect mediation effect.

| Name | Count | p-value | Gene | $ab$ | Category |
|---|---|---|---|---|---|
| MAPK signaling pathway | 9 | 2.3E-04 | ELK4, FOS, RASGRP4, CACNA2D2, DUSP1, FGF9, HSPA8, MAPK1, PPP3CB | 0.0197 | KEGG |
| Kaposi sarcoma-associated herpesvirus infection | 9 | 1.8E-03 | AKT3, FOS, GNB1, BECN1, IFNA1, MAPK1, PIK3CG, PTGS2, UBA52 | 0.0384 | KEGG |
| Chemokine signaling pathway | 8 | 6.7E-03 | AKT3, CCL27, CCL4, GNB1, DOCK2, MAPK1, PARD3, PIK3CG | 0.0263 | KEGG |
| Toll-like receptor signaling pathway | 6 | 8.2E-03 | AKT3, CCL4, FOS, IFNA1, MAPK1, MAP3K8 | 0.0266 | KEGG |
| TNF signaling pathway | 6 | 1.0E-02 | AKT3, CEBPB, FOS, MAPK1, MAP3K8, PTGS2 | 0.0359 | KEGG |

Table 19: The significant pathways identified for HDL-C. $ab$ refers to the indirect mediation effect.

| Name | Count | p-value | Gene | $ab$ | Category |
|---|---|---|---|---|---|
| Antigen processing and presentation | 4 | 1.3E-02 | CD74, CTSS, HSPA4, HLA-DOB | -0.1821 | KEGG |
| Asthma | 3 | 1.6E-02 | IL4, HLA-DOB, MS4A2 | 0.9006 | KEGG |
| Pancreatic secretion | 4 | 2.6E-02 | ATP2B4, ATP2A2, RAB27B, PRKCG | 0.1289 | KEGG |
| Cholesterol metabolism | 3 | 3.4E-02 | CETP, LDLR, MYLIP | 0.2758 | KEGG |
| Regulation of actin cytoskeleton | 5 | 4.0E-02 | DIAPH2, FGF10, ITGB8, PDGFRA, PTK2 | 0.4756 | KEGG |
